# Do closely related species interact with similar partners? Testing for phylogenetic signal in bipartite interaction networks

**DOI:** 10.1101/2021.08.30.458192

**Authors:** Benoît Perez-Lamarque, Odile Maliet, Benoît Pichon, Marc-André Selosse, Florent Martos, Hélène Morlon

## Abstract

Whether interactions between species are conserved on evolutionary time-scales has spurred the development of both correlative and process-based approaches for testing phylogenetic signal in interspecific interactions: do closely related species interact with similar partners? Here we use simulations to test the statistical performances of the two approaches that are the most widely used in the field: Mantel tests and the Phylogenetic Bipartite Linear Model (PBLM). Mantel tests investigate the correlation between phylogenetic distances and dissimilarities in sets of interacting partners, while PBLM is a process-based approach that relies on strong assumptions about how interactions evolve. We find that PBLM often detects a phylogenetic signal when it should not. Simple Mantel tests instead have infrequent false positives and moderate statistical power; however, they often artifactually detect that closely related species interact with dissimilar partners. Partial Mantel tests, which are used to partial out the phylogenetic signal in the number of partners, actually fail at correcting for this confounding effect, and we instead recommend evaluating the significance of Mantel tests with network permutations constraining the number of partners. We also explore the ability of simple Mantel tests to analyze clade-specific phylogenetic signals. We provide general guidelines and an application on an interaction network between orchids and mycorrhizal fungi.

## Introduction

Species in ecological communities engage in numerous types of interspecific interactions, such as pollination, mycorrhizal symbioses, herbivory, and parasitism (Bascompte & Jordano, 2013; Bascompte, Jordano, Melian, & Olesen, 2003; Fontaine et al., 2011; Martos et al., 2012), which are often summarized using bipartite interaction networks (Bascompte & Jordano 2013; Fig. 1). Understanding the processes that shape these interaction networks, including the role of evolutionary history, is a major focus of ecology and evolution (Braga, Janz, Nylin, Ronquist, & Landis, 2021; Elias, Fontaine, & Frank Van Veen, 2013; Futuyma & Agrawal, 2009; Gómez, Verdú, & Perfectti, 2010; Krasnov et al., 2012; Rezende, Lavabre, Guimarães, Jordano, & Bascompte, 2007; Rohr & Bascompte, 2014; Vázquez, Chacoff, & Cagnolo, 2009). One way to assess the role of evolutionary history in shaping contemporary interactions is to test for a phylogenetic signal in species interactions, *i*.*e*. whether closely related species interact with similar sets of partners (Peralta, 2016).

**Figure 1:**
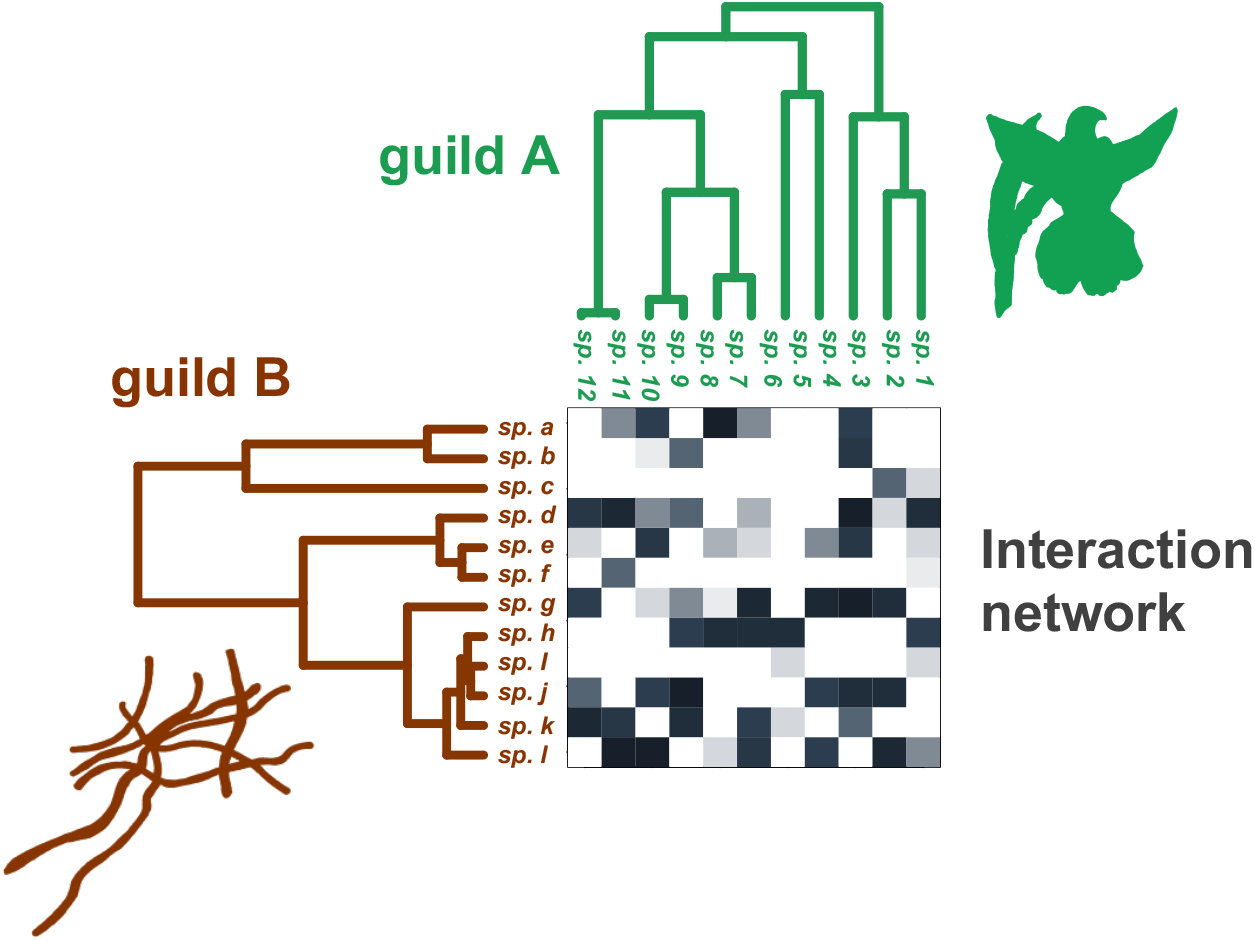
Illustration of the data used to test for phylogenetic signal in species interactions Toy example of an interaction network between orchids (in green) and mycorrhizal fungi (in brown) with associated phylogenetic trees. The bipartite interaction network between two guilds A (here the orchids) and B (the fungi) is represented by a matrix, which elements indicate either whether or not species interact (*i*.*e*. 1 if they do and 0 otherwise, “unweighted” or “binary” network) or the interaction frequency (“weighted” network; for example here we indicated the number of times a given pairwise interaction has been observed using shades of gray from white (no interaction) to dark gray (many interactions)). Each guild is also characterized by a rooted phylogenetic tree, used to compute phylogenetic distances between species pairs.

Testing for a phylogenetic signal in a trait for a given clade, *i*.*e*. whether a trait is phylogenetically conserved, is mainstream (Blomberg, Garland, & Ives, 2003; Felsenstein, 1985; Münkemüller et al., 2012). One approach (the “correlative” approach) is to perform a Mantel test between phylogenetic and trait distances (Mantel, 1967); another approach (the “process-based” approach) relies on trait evolution models such as Pagel’s λ (Pagel, 1999) or Blomberg’s *K* (Blomberg et al., 2003). The process-based approach has a higher ability to detect an existing phylogenetic signal (power) and a lower propensity to infer a phylogenetic signal when it should not (false positive; Harmon & Glor 2010): The correlative approach should therefore only be used when the process-based approach is not applicable, *e*.*g*. if the trait data is expressed in terms of pairwise distances.

Testing for a phylogenetic signal in species interactions falls in the category of cases where the trait data are pairwise distances, here the between-species dissimilarity in sets of interacting species. Simple Mantel tests have therefore been widely used in this context (*e*.*g*. Cattin *et al*. 2004; Rezende *et al*. 2007; Elias *et al*. 2013; Fontaine & Thébault 2015). Partial Mantel tests have also been used to test whether the phylogenetic signal reflects more the identity of the interacting partners than their number, as similarity in the number of partners can increase the value of similarity metrics (“phylogenetic signal in the number of partners”; Rezende *et al*. 2007; Jacquemyn *et al*. 2011; Aizen *et al*. 2016). Mantel tests, which are easy and fast to run and that do not rely on strong hypotheses, have therefore been vastly used to test for phylogenetic signal in species interactions in empirical networks (Cattin et al., 2004; Elias et al., 2013; Fontaine & Thébault, 2015; Jacquemyn et al., 2011; Rezende et al., 2007). Besides these correlative approaches, several process-based approaches have been developed (Hadfield, Krasnov, Poulin, & Nakagawa, 2014; Ives & Godfray, 2006; Li, Dinnage, Nell, Helmus, & Ives, 2020; Rafferty & Ives, 2013). The first of these approaches, the Phylogenetic Bipartite Linear Model (PBLM, Ives & Godfray 2006), has been widely used to test for phylogenetic signal in species interactions in a variety of networks, *e*.*g*. in host-parasite, plant-fungus, and pollination networks (Ives & Godfray, 2006; Martín González et al., 2015; Martos et al., 2012; Xing et al., 2020). In short, PBLM assumes that interaction strengths between species from the two guilds are determined by (unobserved) traits that evolve on the two phylogenies each following a simplified Ornstein-Uhlenbeck process (Blomberg et al., 2003). PBLM performs a phylogenetic regression to infer the Ornstein-Uhlenbeck parameters, which are then interpreted in terms of phylogenetic signal (Ives & Godfray 2006). Other models have been developed more recently (Hadfield et al., 2014; Li et al., 2020; Rafferty & Ives, 2013), including the phylogenetic generalized linear mixed model (PGLMM; Rafferty and Ives 2013) that uses linear mixed models to infer phylogenetic signals in both the number of partners and species interactions. Yet, the higher computational requirements of these methods have prevented their widespread use on empirical networks. PBLM thus remains the method frequently used in empirical studies (*e*.*g*. Xing et al. 2020; Corro et al. 2021).

Mantel tests and PBLM sometimes provide contradictory conclusions on empirical data and this is difficult to interpret because the statistical performances of the two approaches have never been compared (Peralta, 2016). Importantly, the statistical performances of PBLM have not been tested. Here, we use simulations to perform a comparative analysis of the statistical performances of these approaches. We consider both weighted and unweighted bipartite interaction networks between species from two guilds A and B (Fig. 1). Our results lead us to propose alternative approaches for measuring phylogenetic signal in interaction networks. We also investigate the ability of Mantel tests to detect the presence of phylogenetic signal in the different clades of a phylogenetic tree, as phylogenetic signal may be limited to some sub-clades. Finally, we provide general guidelines and illustrate them on an orchid-fungus mycorrhizal network identified across the oceanic island of Réunion (Martos et al., 2012).

## Methods

### Simulating bipartite interaction networks with or without phylogenetic signal in species interactions

We used *BipartiteEvol*, an individual-based eco-evolutionary model (Figure 2; see Maliet *et al*. 2020 for a complete description of the model), to generate interaction networks with or without phylogenetic signal between two guilds interacting in a mutualistic, antagonistic, or neutral way. In short, each individual from guild A (resp. B) is characterized by a multidimensional continuous trait and interacts with one individual from guild B (resp. A). The effect of this interaction on the fitness of each individual from guilds A or B is determined by the distance in trait space of the two interacting individuals, according to a classical trait matching expression parametrized by two parameters α_A_ and α_B_ (Supplementary Methods 1, Maliet *et al*. 2020). These parameters determine the nature and specificity of the interaction: positive α_A_ and α_B_ correspond to mutualistic interactions, negative α_A_ and positive α_B_ to antagonistic interactions (with guild A representing hosts/preys and guild B parasites/predators), high | α |values to scenarios with strong fitness effects (*i*.*e*. highly specialized interactions), and | α |values close to 0 to more neutral scenarios (Figure 2). *BipartiteEvol* simulates the random death of individuals that are replaced by new ones proportionally to their fitness. At birth, new individuals have a probability *μ* to mutate, leading to new trait values slightly different from the parental ones (Figure 2). Such mutations can lead to the formation of new species. Networks simulated using *BipartiteEvol* show typical structural properties observed in empirical networks, including significant nestedness and/or modularity according to the sets of simulated parameters (Maliet et al., 2020). For instance, networks simulated with antagonistic interactions (α_A_<0) tend to be significantly modular, while networks simulated with neutral or mutualistic interactions (α_A_=0 or α_A_<0) tend to be nested. Here, instead of using the species delineation of the original *BipartiteEvol* model (Maliet et al., 2020), we considered that each combination of traits corresponds to a distinct species. This increased our ability to generate phylogenetic signal in the simulated networks, and we show that it does not affect their overall structure (see below).

**Figure 2:**
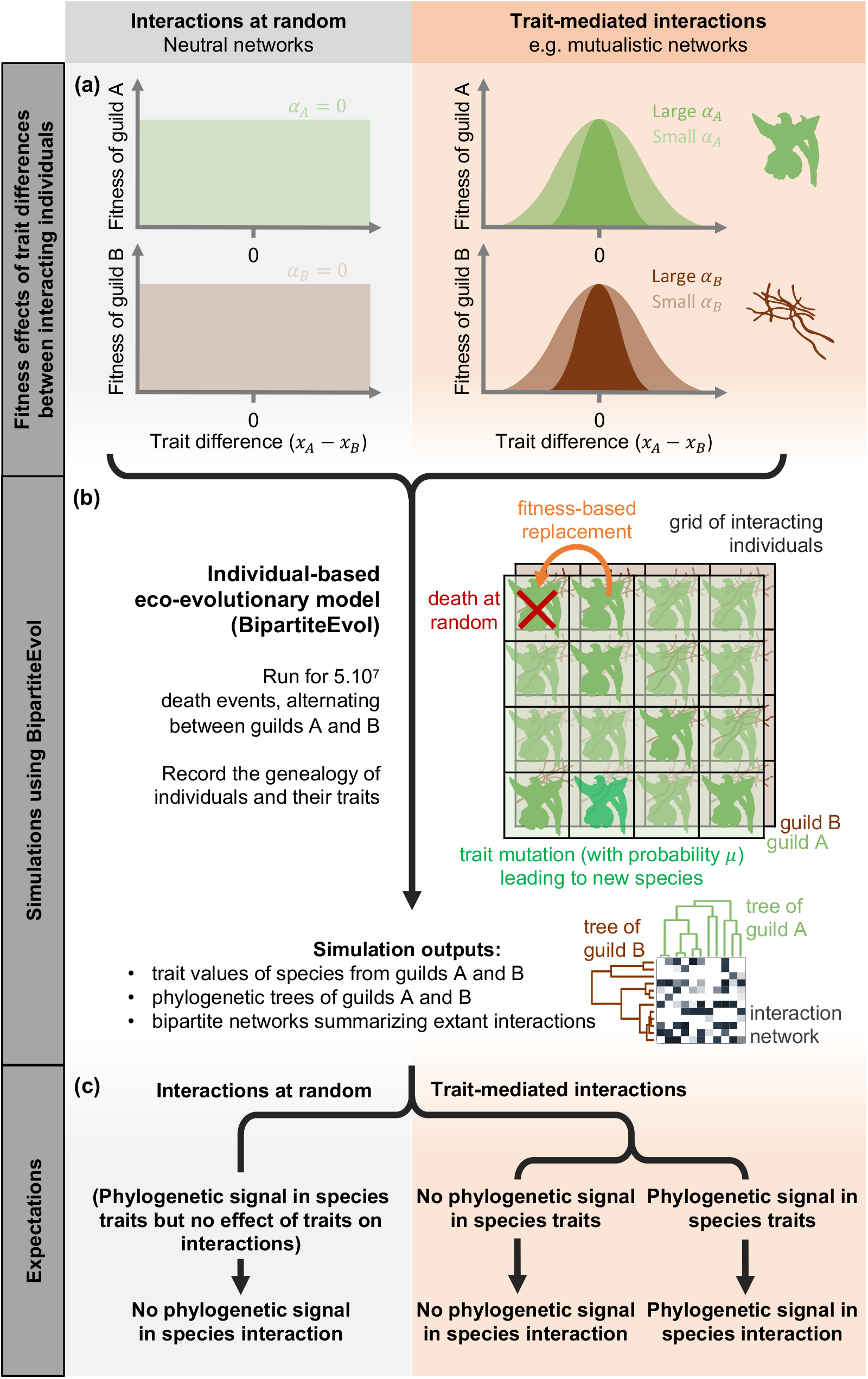
Simulation scheme used to generate interaction networks with or without phylogenetic signal in species interactions: **(a)** The fitness of a given individual is either affected by its trait value and that of the individual it interacts with (right; “mutualistic or antagonistic interactions”) or not (left; “neutral interactions”). In the first case, fitness depends on the trait matching between the pair of interacting individuals (*x*_*A*_− *x*_*B*_), where *x*_*A*_ (resp. *x*_*B*_) are the trait values of the individual from guild A (resp. B). The strength of the effect of the traits on fitness is modulated by the parameters *α* _*A*_ and *α* _*B*_ (|*α* | values close to 0 tend toward neutrality, where interactions happen at random). **(b)** *BipartiteEvol* assumes that pairs of individuals from guilds A and B interact on a grid of a given size. Each cell of the grid contains one pair of individuals. The process starts with one monomorphic species in each guild. At each time step, one individual of guild A is killed at random and replaced by another individual proportionally to its fitness in the cell. The new individual can mutate with probability *μ*, in which case the new trait value is drawn from a normal distribution centered on the parental trait. A mutation generates a new species. The same events are applied to an individual from guild B, and the process is repeated a large number of time steps. This simulation model outputs a list of species of guilds A and B with associated trait values, phylogenetic trees, and the bipartite interaction network of extant species. **(c)** A phylogenetic signal in species interactions can occur only when trait values modulate interactions (for large *α*) and there is a phylogenetic signal in trait values. When interactions are completely or quasi-random (*α* =0 or low *α*), there cannot be a phylogenetic signal in species interactions.

Under the *BipartiteEvol* model, closely related species tend to interact with similar sets of partners, *i*.*e*. there is a phylogenetic signal in species interactions, if and only if: (1) closely related species have similar traits, *i*.*e*. there is a phylogenetic signal in species traits, and (2) these traits determine who interacts with whom, *i*.*e. α* ≠ 0(Figure 2). Similarly, a negative phylogenetic signal in species interactions, *i*.*e*. the tendency for closely related species to associate with dissimilar partners, is expected if there is a negative phylogenetic signal in species traits, *i*.*e*. closely related species have dissimilar traits, and *α* ≠ 0.

We used the *sim*.*BipartiteEvol* function from the R-package RPANDA (Morlon et al., 2016; R Core Team, 2022) to simulate a total of 2,400 interaction networks. To obtain a wide range of network sizes, we considered a total number of 500, 1,000, 2,000, 3,000, 4,000, or 5,000 pairs of interacting individuals per simulation (Table 1a). For each size, we simulated the evolution of 100 neutral networks (α_A_=0 ; α_B_=0), 120 mutualistic networks (**i**: α_A_=1; α_B_=1; **ii**: α_A_=0.1; α_B_=0.1; **iii**: α_A_=0.01; α_B_=0.01; **iv**: α_A_=1; α_B_=0.1; **v**: α_A_=1; α_B_=0.01; and **vi**: α_A_=0.1; α_B_=0.01) and 180 antagonistic networks (**i**: α_A_=-1; α_B_=1; **ii**: α_A_=-0.1; α_B_=0.1; **iii**: α_A_=-0.01; α_B_=0.01; **iv**: α_A_=-1; α_B_=0.1; **v**: α_A_=-1; α_B_=0.01; **vi**: α_A_=-0.1; α_B_=1; **vii**: α_A_=-0.1; α_B_=0.01; **viii**: α_A_=-0.01; α_B_=1; **ix**: α_A_=-0.01; α_B_=0.1). Each individual was characterized by a six-dimensional trait, and trait mutation occurred at birth with a probability *μ*=0.01. Upon mutation, the new trait values were drawn independently in a normal distribution centered on the parental traits and with a variance of 1. We followed the interacting individuals during N=5.10_7_ death events. In the end, we extracted for each guild a species tree from its genealogy by randomly selecting one individual per species (Fig. S1), we also recorded the number of individuals belonging to each species, and counted the number of occurrences of each interspecific interaction; we then reconstructed the corresponding weighted interaction network.

**Table 1:**
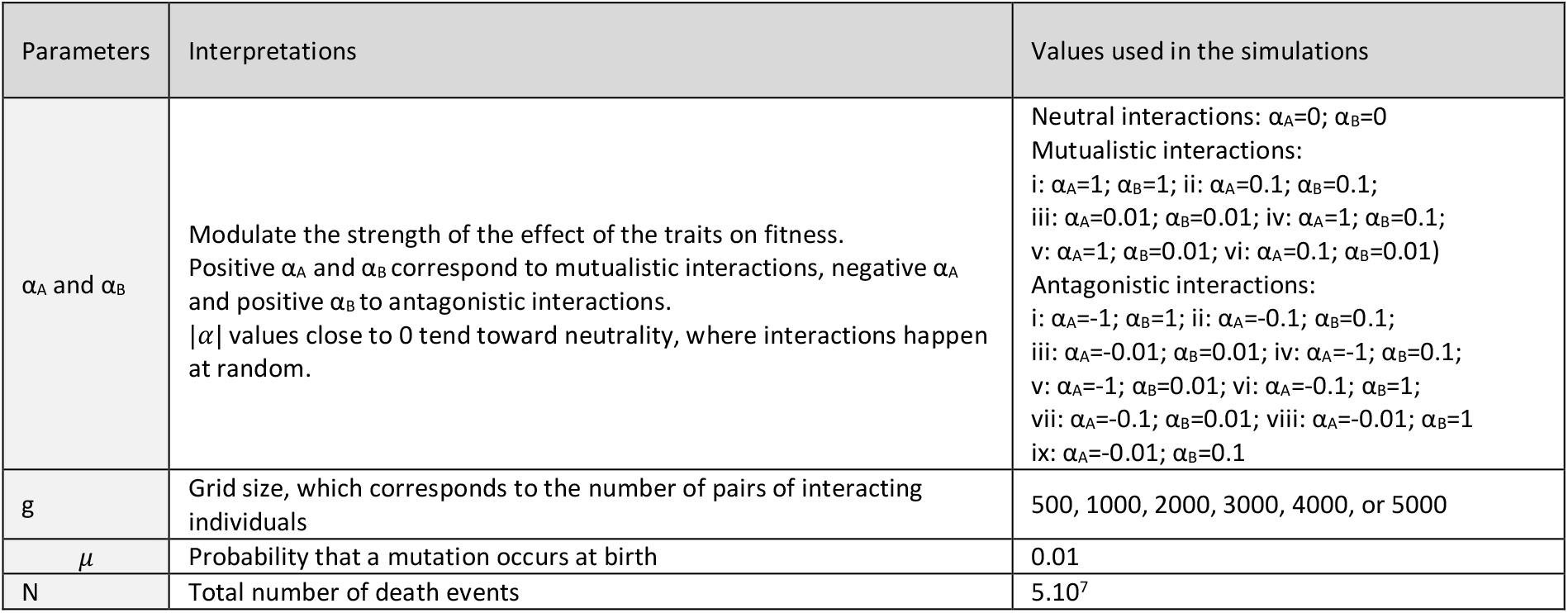
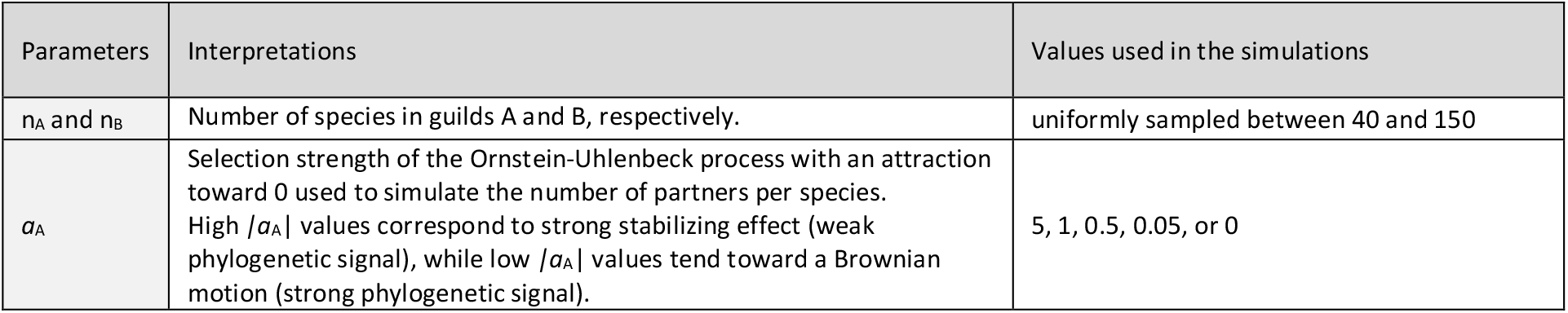
Parameters used in the simulations: **(a)** Parameters used for simulating interaction networks with or without phylogenetic signal in species interactions using BipartiteEvol **(b)** Parameters used for simulating interaction networks with or without phylogenetic signal in the number of partners:

| Parameters | Interpretations | Values used in the simulations |
| --- | --- | --- |
| $\alpha_A$ and $\alpha_B$ | Modulate the strength of the effect of the traits on fitness. Positive $\alpha_A$ and $\alpha_B$ correspond to mutualistic interactions, negative $\alpha_A$ and positive $\alpha_B$ to antagonistic interactions. $\vert \alpha\vert $ values close to 0 tend toward neutrality, where interactions happen at random. | Neutral interactions: $\alpha_A=0$; $\alpha_B=0$<br>Mutualistic interactions:<br>i: $\alpha_A=1$; $\alpha_B=1$; ii: $\alpha_A=0.1$; $\alpha_B=0.1$;<br>iii: $\alpha_A=0.01$; $\alpha_B=0.01$; iv: $\alpha_A=1$; $\alpha_B=0.1$;<br>v: $\alpha_A=1$; $\alpha_B=0.01$; vi: $\alpha_A=0.1$; $\alpha_B=0.01$)<br>Antagonistic interactions:<br>i: $\alpha_A=-1$; $\alpha_B=1$; ii: $\alpha_A=-0.1$; $\alpha_B=0.1$;<br>iii: $\alpha_A=-0.01$; $\alpha_B=0.01$; iv: $\alpha_A=-1$; $\alpha_B=0.1$;<br>v: $\alpha_A=-1$; $\alpha_B=0.01$; vi: $\alpha_A=-0.1$; $\alpha_B=1$;<br>vii: $\alpha_A=-0.1$; $\alpha_B=0.01$; viii: $\alpha_A=-0.01$; $\alpha_B=1$;<br>ix: $\alpha_A=-0.01$; $\alpha_B=0.1$ |
| $g$ | Grid size, which corresponds to the number of pairs of interacting individuals | 500, 1000, 2000, 3000, 4000, or 5000 |
| $\mu$ | Probability that a mutation occurs at birth | 0.01 |
| $N$ | Total number of death events | $5.10^7$ |

**(b) Parameters used for simulating interaction networks with or without phylogenetic signal in the number of partners:**
| Parameters | Interpretations | Values used in the simulations |
| --- | --- | --- |
| $n_A$ and $n_B$ | Number of species in guilds A and B, respectively. | uniformly sampled between 40 and 150 |
| $a_A$ | Selection strength of the Ornstein-Uhlenbeck process with an attraction toward 0 used to simulate the number of partners per species. High $\vert a_A\vert $ values correspond to strong stabilizing effect (weak phylogenetic signal), while low $\vert a_A\vert $ values tend toward a Brownian motion (strong phylogenetic signal). | 5, 1, 0.5, 0.05, or 0 |

First, we evaluated whether these simulations generated realistic networks by comparing their structure with that of empirical networks. Empirical networks were gathered from the Web of Life database (web-of-life.es (Fortuna, Ortega, & Bascompte, 2014)) and the database of Michalska-Smith & Allesina (2019). We compared the structures of simulated *versus* empirical networks in terms of connectance, nestedness, and modularity (Supplementary Methods 2).

Second, we separated the 2,400 simulated networks between those for which we should expect a phylogenetic signal in species interactions and those for which we should not (Figure 2). We did not expect any phylogenetic signal in species interactions in neutral networks and in non-neutral networks with no phylogenetic signal in species traits. Conversely, we expected phylogenetic signal in non-neutral networks with phylogenetic signal in species traits. We evaluated phylogenetic signal in species traits using two approaches. First, for simplicity and consistency with the rest of the paper, we used Mantel tests (Pearson correlation) between phylogenetic distances and trait distances computed as the Euclidian distances between trait values for each species pair. Second, given that process-based approaches usually perform better (Harmon & Glor, 2010), we used a multivariate extension of Pagel’s λ (Pagel, 1999) implemented in R (Goolsby, 2015); we assessed the significance of the phylogenetic signal in species traits with likelihood ratio tests comparing the inferred Pagel’s λ model to a null model where λ=0 (*i*.*e*. no phylogenetic signal).

### Computing phylogenetic signal in species interactions

We computed phylogenetic signal in species interactions in the simulated networks using Mantel tests and PBLM, as well as the computationally-intensive PGLMM for the smallest networks. Complete descriptions of these methods are available in Supplementary Methods 3. Mantel tests, PBLM, and PGLMM rely on different strategies to evaluate the significance of the phylogenetic signal, and it could be argued that the results of these tests are not directly comparable. Our approach is to follow the methodologies traditionally used in empirical studies and compare their conclusions (detection or not of a phylogenetic signal).

#### Mantel tests

We evaluated the phylogenetic signal in species interactions in guilds A and B separately using simple Mantel tests between phylogenetic and ecological (set of interacting partners) distances. Ecological distances were measured both without accounting for evolutionary relatedness of the interacting partners, using (weighted or unweighted) Jaccard, and accounting for relatedness using (weighted or unweighted) UniFrac distances (Supplementary Methods 3 (Lozupone, Lladser, Knights, Stombaugh, & Knight, 2011)). Accounting for evolutionary relatedness of the interacting partners can be particularly relevant for organisms with uncertain species delineations (e.g. microorganisms delineated using only molecular data (Martos et al., 2012; Sanders et al., 2014)). We used Pearson, Spearman, and Kendall correlations (R) by extending the *mantel* function in the R-package ecodist (Goslee & Urban, 2007); the significance of each correlation was evaluated using 10,000 permutations, except for the computationally intensive Kendall correlation (100 permutations only). For each network, we considered that there was a significant positive phylogenetic signal (resp. negative phylogenetic signal) if the correlation coefficient (R) was higher (resp. lower) than >95% of the randomized correlations; we computed the p-value of each one-tailed Mantel test as the fraction of the randomized correlations above (resp. below) the original value.

#### PBLM

To estimate phylogenetic signal based on PBLM, we modified the function *pblm* from the R-package picante (Kembel et al., 2010) to more efficiently perform matrix inversions and handle large interaction networks. In short, the parameters d_A_ and d_B_ of the Ornstein-Uhlenbeck processes of PBLM were estimated using generalized least squares (Ives & Godfray 2006). d_A_ and d_B_ are interpreted as a measure of phylogenetic signal in species interactions: if d=0, there is no effect of the phylogeny (similar as evolution on a star phylogeny, *i*.*e*. no phylogenetic signal); 0<d<1 generates stabilizing selection (*i*.*e*. phylogenetic signal) and d>1 disruptive selection (*i*.*e*. negative phylogenetic signal). We followed Ives & Godfray (2006; Supplementary Methods 3) by considering that the phylogenetic signal is significant when the mean square error (MSE) of the model is smaller than that obtained using star phylogenies (MSE_star_); we also used a more stringent criterion by considering that the signal is significant when the MSE is at least 5% lower than MSE_star_. Finally, we applied the bootstrapping method of Ives & Godfray (2006; Supplementary Methods 3) to the smallest networks. A single PBLM inference can take several days to run (time measured on an Intel 2.8 GHz MacOSX laptop) on networks of intermediate sizes (*e*.*g*. between 50 and 100 species per guild), which prevented us from applying the bootstrap approach to large networks; we therefore only tested this approach on networks simulated with 500 individuals (*i*.*e*. a total of 400 networks).

#### PGLMM

We performed analyses of the statistical performances of PGLMM (Rafferty & Ives, 2013) using the function *pglmm* in the R-package phyr (Li et al., 2020). Following the procedure used in Lajoie and Kembel (2021), we fitted for each network different models accounting or not for phylogenetic signal in both the number of partners and in the species interactions in both clades, using restricted maximum likelihood and evaluating significance with likelihood ratio tests. Because fitting these models can require a large amount of memory (*e*.*g*. >80 Gb for some networks with >50 species per guild) and long computation time (Fig. S2), we only tested this approach on networks simulated with 500 individuals. We fitted the PGLMM using either a Gaussian or a Poisson distribution of abundances for weighted networks, and a binomial distribution (presence/absence data) for unweighted networks (Li et al., 2020).

### Confounding effect of the phylogenetic signal in the number of partners

To test the performances of the partial Mantel test at measuring phylogenetic signal in species interactions while controlling for the number of partners (Supplementary Methods 3), we first performed partial Mantel tests between phylogenetic and ecological distances, while controlling for pairwise differences in the number of partners, on the networks simulated with *BipartiteEvol*. There is no reason that *BipartiteEvol* simulations generate a phylogenetic signal in the number of partners, and we verified this by performing Mantel tests between phylogenetic distances and pairwise differences in the number of partners. These analyses thus assess whether partial Mantel tests lose power compared to simple Mantel tests in the absence of a phylogenetic signal in the number of partners. If they do not suffer power loss, partial Mantel tests applied to *BipartiteEvol* simulations should be significant when simple Mantel tests are significant.

Second, we assessed whether partial Mantel tests successfully correct for the phylogenetic signal in the number of partners using networks simulated under a process that generates phylogenetic conservatism in the number, but not the identity, of interacting partners (*i*.*e*. partial Mantel tests should not be significant when applied to such networks). To simulate networks with only phylogenetic conservatism in the number of partners in guild A, we first simulated phylogenetic trees for guilds A and B using *pbtree* (R-package phytools; Revell 2012) with a number of species uniformly sampled between 40 and 150 by guilds. Next, we simulated the number of partners of the species from guild A using an Ornstein-Uhlenbeck process with an attraction toward 0, a variance of 0.1 (noise of the Brownian motion), and a selection strength (*a*_A_) ranging from 5 (strong stabilizing effect, weak phylogenetic signal) to 0 (Brownian motion, strong phylogenetic signal). We computed the number of partners per species by calibrating the simulated values between 1 and the number of species in guild B and taking the integer part. For each *a*_A_ value (5, 1, 0.5, 0.05, or 0), we performed 100 simulations using *mvSIM* (R-package mvMORPH; Clavel *et al*. 2015; Table 1b). Finally, for each species in A, we attributed the corresponding number of partners in B at random to obtain binary networks. We checked that our simulations indeed generated a signal in the number of partners by performing simple Mantel tests between phylogenetic distances and pairwise differences in the number of partners. Finally, we performed on each simulated network a partial Mantel test between phylogenetic and ecological distances, while controlling for pairwise differences in the number of partners.

Given the poor performances of partial Mantel tests (see Results), we tested three alternative approaches to partial out the confounding effect of the number of partners in measures of phylogenetic signal. First, we tested whether using sequential Mantel tests would provide a good alternative: based on simple Mantel tests, we consider that there is a phylogenetic signal in the identity of the partners if there is a phylogenetic signal in species interactions and no phylogenetic signal in the number of partners. Second, we tested the use of methods that directly partition ecological distance metrics into a part due to the dissimilarity in the number of partners and a part due to the dissimilarity in the identity of the partners, *i*.*e*. “species turnover” (Baselga, 2010). We used the betapart R-package (Baselga & Orme, 2012) to extract the part of the unweighted Jaccard distances due to species turnover and tested its correlation with phylogenetic distances using a simple Mantel test. Third, we designed specific network permutations to test for the significance of the Mantel correlation between phylogenetic distances and ecological distances while accounting for the number of partners. To measure whether the phylogenetic signal observed in guild A is not due to a phylogenetic signal in the number of partners, instead of shuffling the distance matrix as in a regular Mantel test (Supplementary Methods 3), we randomized the interaction network by keeping constant the number of partners per species from guild A while permuting the partner identities. Because this third approach requires recomputing the ecological distances for each permutation, it is much slower than regular Mantel tests (Fig. S2) and we thus used only 1,000 permutations. We applied these three methods to all our simulated networks.

### Effect of phylogenetic uncertainty, sampling asymmetry, and network heterogeneity on measures of phylogenetic signal in species interactions

Unlike simulations, such as those provided by *BipartiteEvol*, empirical bipartite networks suffer from (i) uncertainty in the phylogenetic reconstructions, *e*.*g*. in the microbial partners’ tree when studying host-associated microbiota, which often prevents accounting for evolutionary relatedness (*i*.*e*. using UniFrac distances), (ii) sampling asymmetry, *i*.*e*. one side of the network is more thoroughly sampled than the other, and (iii) network heterogeneity, *i*.*e*. different sub-clades in the network have different levels of phylogenetic signal. We performed additional analyses to investigate the effect of these aspects on phylogenetic signals in species interactions measured using simple Mantel tests.

First, we tested the effect of phylogenetic uncertainty in the partners’ tree on the measure of phylogenetic signal when evolutionary relatedness is accounted for (*i*.*e*. using UniFrac distances). We performed these analyses to assess whether accounting for the partners’ evolutionary relatedness remains advantageous (see Results) when phylogenetic uncertainty is high. To add some variability in the phylogenetic tree of guild B (resp. A) used to compute the UniFrac distances between species pairs from guild A (resp. B), we first simulated, on the original partners tree, the evolution of a short DNA sequence and then reconstructed the tree from the simulated DNA alignment using neighbor-joining (*nj* function, R-package APE (Paradis, Claude, & Strimmer, 2004)). We used *simulate_alignment* (R-package HOME; Perez-Lamarque & Morlon 2019) to simulate sequences of length 75, 150, 300, 600, or 1,200 base-pairs, with 30% of variable sites, and a substitution rate of 1.5 (shorter fragments should result in noisier phylogenies). To assess the uncertainty of these reconstructed phylogenetic trees compared with the original trees, we computed the correlations between the pairwise phylogenetic distances in both trees.

Second, we tested the influence of sampling asymmetry on measures of phylogenetic signal. Empirical networks are often an incomplete representation of the actual interactions between two guilds because they are under-sampled, and frequently, in an asymmetrical way. For instance, by sampling targeted species from guild A, observed networks are constituted by a few species from guild A which have the complete set of their partners and by often more species from guild B which have an incomplete set of their partners (as they likely interact with unsampled species from guild A). We tested the influence of such sampling asymmetry by selecting only 10% of the most abundant species from guild A in each simulated network (while retaining at least 10 species) and computed phylogenetic signals in these asymmetrically-subsampled networks.

Third, both Mantel tests and PBLM neglect the heterogeneity within networks. Indeed, a non-significant phylogenetic signal at the level of the entire network can potentially hide a sub-clade of species presenting a significant phylogenetic signal. Alternatively, a phylogenetic signal in the entire network may be driven by only two sub-clades of guilds A and B, while the other sub-clades present no significant phylogenetic signal. To explore the potential heterogeneity of the phylogenetic signal within one guild, one possibility is to apply Mantel tests to the sub-networks formed by a given sub-clade (*e*.*g*. Song *et al*. 2020). For each node of the tree of guild A having at least 10 descendants, we estimated the clade-specific phylogenetic signal using a Mantel test investigating whether closely related species from this sub-clade of A tend to interact with similar partners (and *vice-versa* for guild B). Using UniFrac distances, we performed the Mantel tests with 100,000 permutations and introduced a Bonferroni correction for multiple testing to keep a global risk of false positives of 5%. To test this approach, we simulated networks with known sub-clade signals by artificially combining networks simulated under neutrality with networks simulated with the mutualistic parameters **v** (see Results). We grafted each “mutualistic” phylogenetic tree from guilds A and B within a “neutral” phylogenetic tree by randomly selecting a branch, such that it creates a separate module with a strong phylogenetic signal. Such simulations could correspond to the evolution of a different niche, *e*.*g*. terrestrial *versus* epiphytic plants associating with different mycorrhizal fungi (Martos et al., 2012). We then performed our clade-specific analysis of phylogenetic signals and investigated in which nodes we recovered a significant phylogenetic signal.

### General guidelines and illustration with application on the orchid-fungus mycorrhizal network from La Réunion

We used our results and other empirical considerations to provide general guidelines for testing for phylogenetic signal in interaction networks. We illustrated these guidelines by applying them in a network between orchids and mycorrhizal fungi from La Réunion island (Martos et al., 2012). This network encompasses 70 orchid species (either terrestrial or epiphytic species) and 93 molecularly-identified fungal partners (defined according to 97% sequence similarity; Martos *et al*. 2012). We gathered the maximum-likelihood plant and fungal phylogenies on TreeBASE (Study Accession S12721), calibrated the orchid phylogeny using a relaxed clock with *chronos* (Paradis, 2013), and set the divergence between Orchidoideae and Epidendroideae at 65 million years (Givnish et al., 2015). To obtain a species-level orchid phylogeny, missing species were grafted in the phylogeny by arbitrarily adding 4 million-year-old polytomies in the corresponding unresolved genera, namely *Habenaria, Benthamia, Cynorkis, Phaius, Liparis, Bulbophyllum*, and *Polystachya*.

## Results

### Expected phylogenetic signals in species interactions in *BipartiteEvol* networks

*BipartiteEvol* simulations resulted in interaction networks with a large range of sizes for guilds A and B, from less than 50 to more than 250 species (Fig. S3). These simulated networks had similar structural properties as empirical networks, including in terms of connectance, nestedness, and modularity (Fig. S4). This means that networks simulated using *BipartiteEvol* are realistic and cover a large range of structures encountered in natural interaction networks.

Using Mantel tests, we found a significant phylogenetic signal in species traits for most antagonistic and neutral simulations (Fig. S5A). In contrast, for many mutualistic simulations, closely related species often did not tend to have similar traits, except when α_B_=0.01 (*i*.*e*. mutualistic parameters **iii, v**, and **vi**; Fig. S5A). When α_B_ were higher (*i*.*e*. mutualistic parameters **i, ii**, and **iv)**, we suspect stabilizing selection to occur and erase the phylogenetic signal in the traits (Maliet et al., 2020): we therefore do not expect any phylogenetic signal in species interactions for these simulations, which represent ∼40% of the mutualistic simulations. In addition, we found a negative phylogenetic signal in species traits (suggesting that closely related species have dissimilar traits) in less than 1% of the simulations (Fig. S5A). Given that we do not expect *BipartiteEvol* to generate a negative phylogenetic signal in species traits and given that the risk of false positives of a Mantel test is 5%, these 1% of networks with a negative phylogenetic signal in species traits are likely false positives. We removed them when evaluating the performance of the different approaches and we therefore do not expect any negative phylogenetic signal in species interactions for the networks we tested, *i*.*e*. closely related species should not tend to associate with dissimilar partners. Results were similar when using Pagel’s λ, with a significant phylogenetic signal in species traits for almost all antagonistic and neutral simulations, and in ∼65% of the mutualistic simulations (Fig. S5B). Mantel tests and Pagel’s λ lead to identical conclusions for >95% of the simulated networks.

### Computing phylogenetic signals in species interactions in *BipartiteEvol* networks

Using Mantel tests, as expected, we did not find a significant phylogenetic signal in species interactions for most neutral networks or for networks with no signal in species traits (Figs. 3 & S6): the false positive rate was below 5%, corresponding to the risk of false positives of the test (Table S1), with one notable exception for small networks when using weighted Jaccard distances and Pearson correlations (∼ 8% of false positives). Conversely, we detected a significant unexpected negative phylogenetic signal in more than 10% of the simulated networks, in particular in the small ones (Figs. 3 & S6).

**Figure 3:**
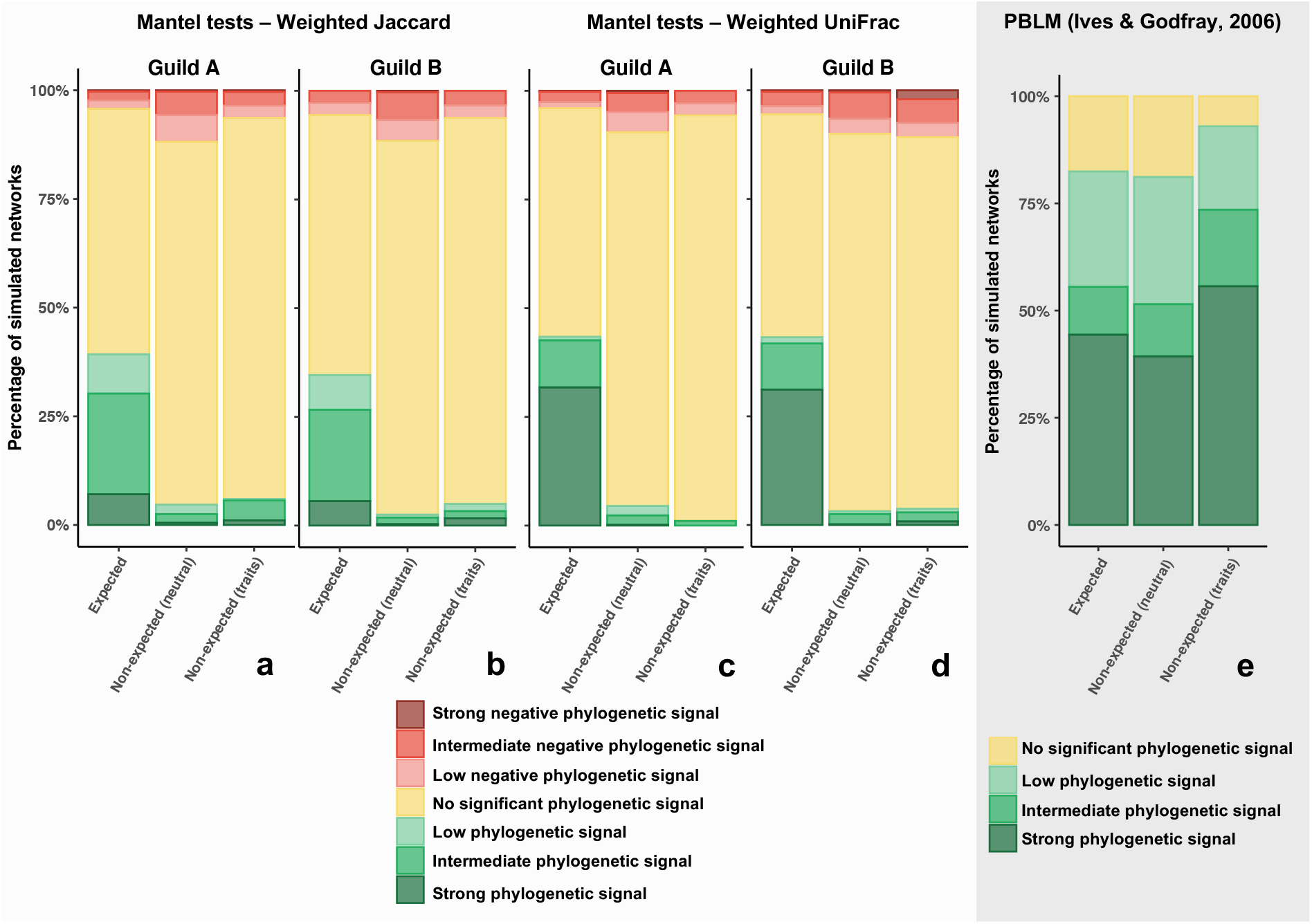
Statistical performances of the simple Mantel tests and the Phylogenetic bipartite linear model (PBLM; Ives & Godfray, 2006). For each panel, the simulations are divided between networks where a phylogenetic signal in species interactions is expected (*i*.*e*. networks (i) simulated with an effect of the traits on individual fitness - antagonistic and mutualistic simulations - and (ii) presenting traits that are phylogenetically conserved according to a Mantel test – see Fig. S5A) and networks where a phylogenetic signal in species interactions is not expected (*i*.*e*. neutral simulations (*α* = 0) or simulated networks where we observed no phylogenetic signal in the traits). Results are similar when the expectations are based on Pagel’s λ to measure the phylogenetic signals in species traits (Fig. S11). **(a-d):** Phylogenetic signals in species interactions estimated using simple Mantel tests with Pearson correlation (R) in the guilds A (a, c) and B (b, d). The different panels in rows correspond to the 2 tested ecological distances: weighted Jaccard (a, b) or weighted UniFrac (c, d) distances. One-tailed Mantel tests between phylogenetic distances and ecological distances were performed using 10,000 permutations. In each panel, the bars indicate the percentage of simulated networks that present a significant positive correlation (in green; p-value>0.05 for the test of phylogenetic signal), a significant negative correlation (in red; p-value>0.05 for the test of negative phylogenetic signal), or no significant correlation (in yellow; both p-values>0.05). Significant phylogenetic signals (resp. significant negative phylogenetic signals) are shaded from light green (resp. red) to dark green (resp. red) according to the strength of the signal: we arbitrarily considered a “low signal” when R<0.05 (resp. R>-0.05), an “intermediate signal” when 0.05<R<0.15 (resp. -0.05>R>-0.15), and a “strong signal” when R>0.15 (resp. R<-0.15). **(e):** Phylogenetic signals estimated using PBLM. For a given combination of parameters, the bar indicates the percentage of simulated networks that present no significant (in yellow; MSE<MSE_star_) or a significant (green; MSE<MSE_star_) phylogenetic signal. Phylogenetic signals are shaded from light green to dark green according to the strength of the signal: we arbitrarily considered a “low signal” when d_A_<0.05 and d_B_<0.05, an “intermediate signal” when d_A_>0.05 or d_B_>0.05, and a “strong signal” when d_A_>0.15 or d_B_>0.15. PBLM was run on the weighted networks. In each panel, the first bar indicates the statistical power of the test, whereas the second and third bars indicate the false positive rate of the test. Note that the strength of the phylogenetic signals (based on the R and d values) are not directly comparable. Results discriminating the simulated networks of different sizes and with different sets of parameters are available in Figures S6 & S8.

Many mutualistic or antagonistic networks where we expected a phylogenetic signal in species interactions (*i*.*e*. non-neutral networks with a signal in species traits) presented no significant signal with Mantel tests (Figs. 3 & S6), in particular those simulated with low α_A_ and α_B_ values (*e*.*g*. antagonism **vii;** Table 1a), where non-neutral effects were weak. Mantel tests measuring phylogenetic signals in species interactions were most often not significant unless the phylogenetic signal in species traits was strong (R>0.6; Fig. S7). Even when the phylogenetic signal in species traits was very strong (R>0.9), the phylogenetic signal in species interactions was not significant in many networks. In mutualistic networks, phylogenetic signals in species interactions were present only when there was a large asymmetry in the effects of trait matching on the fitnesses of the species from guilds A or B (case **v**: α_A_=1; α_B_=0.01), *i*.*e*. when only one guild was specialized. Conversely, in antagonistic networks, phylogenetic signals were found mainly when trait matching had a strong impact on the fitness of guild B (the obligate parasites/predators; α_B_≥ 0.1). Additionally, when the phylogenetic signal was significant in one guild, it was generally also significant in the other; in antagonistic networks, the signal was usually higher in guild A compared to guild B (Fig. S6).

The statistical power of Mantel tests measuring phylogenetic signals in species interactions seems to be modulated by network size, as phylogenetic signals were less often significant but generally stronger in smaller networks (Fig. S6). Moreover, Mantel tests based on Pearson correlations had higher power than Spearman and Kendall correlations (Fig. S6) and weighted UniFrac distances performed better than other ecological distances in terms of power in the context of these simulations (Fig. S6; Table S2).

When using mean square errors to evaluate the significance of PBLM, we found a significant phylogenetic signal in species interactions in most of the simulated networks including when we did not expect any (Fig. 3e). The strength and the significance of the inferred phylogenetic signals were independent of the strength of the phylogenetic signal in species traits (Fig. S7). The propensity of PBLM to detect phylogenetic signals decreased in large unweighted networks, but the false positives remained >30%, including when using a more stringent significance cutoff (Figs. S8). Similar results were obtained when bootstrapping to evaluate the significance (Fig. S9). PGLMM on weighted networks with a Gaussian or Poisson distribution had slightly lower but still frequent false positives (>25% or 20%, respectively) and intermediate statistical power (<50%) when measuring phylogenetic signals in species interactions (Fig. S10). PGLMM also often artifactually detected phylogenetic signals in the number of partners (Fig. S10). Conversely, PGLMM on unweighted networks never detected any significant signal (Fig. S10).

We inferred similar statistical performances of both Mantel tests and PBLM when we used Pagel’s λ to evaluate phylogenetic signals in species traits (Figs. S7 and S11).

### Confounding effect of the phylogenetic signal in the number of partners

As expected, tests of phylogenetic signals in the number of partners were non-significant in the large majority of the *BipartiteEvol* networks, especially the larger ones (Fig. S12). We did however observe significant correlations between ecological distances and differences in the number of partners (Fig. S13). Partial Mantel tests testing for phylogenetic signal in species interactions while accounting for the phylogenetic signal in the number of partners had similar false positive rates and power as simple Mantel tests (Figs. S6 & S14; Table S2). Sequential Mantel tests decreased the statistical power by less than 2% (Table S2). Partitioning the ecological distances before running the Mantel test reduced the power by only 5% and resulted in less than 1.5% of false positives, but also resulted in an artefactual detection of negative phylogenetic signals in 9% of the simulations (Table S2; Fig. S15). Finally, Mantel tests with network permutations keeping the number of partners constant increased the statistical power by ∼5% (Table S2) but resulted in an artefactual detection of (positive or negative) phylogenetic signals in ∼10% of the simulations when using Jaccard distances (Fig. S16).

Networks simulated with phylogenetic conservatism in the number, but not the identity of partners covered a realistic range of sizes (Fig. S17). As expected, Mantel tests revealed significant phylogenetic signals in the number of partners in >60% of these networks, with an increasing percentage of significant tests with decreasing a_A_ (*i*.*e*. increasing conservatism in the number of partners; Table 1b; Fig. S18). We found significant correlations between differences in the number of partners and ecological distances in most of these simulated networks (Fig. S19), generating a confounding effect. As a result, simple Mantel tests testing for phylogenetic signal in species interactions without accounting for the phylogenetic signal in the number of partners were frequently significant (>30%; Fig. S20; Table S3). Partial Mantel tests controlling for differences in the number of partners slightly decreased the proportion of false positives, but it remained very high (>25% of false positives; Fig. S21). In addition, partial Mantel tests detected a spurious significant negative phylogenetic signal in species interactions in >15% of the networks (Fig. S21). Conversely, only a few networks with a significant simple Mantel test in species interactions did not produce a significant simple Mantel test in the number of partners, such that sequential Mantel tests had only ∼7% of false positives (Table S3). Partitioning the ecological distances before running the Mantel test (Fig. S22) or using Mantel tests with network permutations keeping the number of partners constant (Fig. S23) had even lower false positive rates (∼4% and ∼5% respectively; Table S3). However, partitioning the ecological distances led to an artefactual detection of negative phylogenetic signals in more than 30% of the simulated networks (Fig. S23).

### Effect of phylogenetic uncertainty, sampling asymmetry, and network heterogeneity on measures of phylogenetic signal in species interactions

As expected, phylogenetic uncertainty in the partner’s tree increased when the length of the simulated DNA sequence used for phylogenetic reconstruction decreased (Fig. S24), resulting in a decreased statistical power of Mantel tests using UniFrac distances (Fig. S25). However, even when the simulated DNA sequences were the shortest (75 base pairs), resulting in very noisy reconstructed partners’ trees (Fig. S26), the statistical power of the Mantel tests using UniFrac distances decreased by less than 5% (Fig. S25).

Our results on the statistical performance of tests of phylogenetic signal were similar when considering sampling asymmetry (Figs. S27-30): PBLM spuriously detected phylogenetic signals when it should not, and Mantel tests had decent statistical performances, especially when using weighted UniFrac distances. In addition, the correlations of the Mantel tests in guild A were generally higher when significant (Fig. S29).

Our clade-specific tests of phylogenetic signals using Mantel tests while correcting for multiple testing recovered a significant phylogenetic signal in 82% of the nodes where mutualism originated (Fig. S31), as well as in most of the ascending nodes. Conversely, we did not find spurious phylogenetic signals in nodes with only neutrally-evolving lineages (Fig. S31).

### General guidelines and illustration with application on the orchid-fungus mycorrhizal network from La Réunion

Figure 4 provides general guidelines based on our results and empirical considerations for accurate tests of phylogenetic signal in interaction networks. We applied these guidelines to the orchid-fungus mycorrhizal network from La Réunion (available in Martos et al. (2012)). First (step 1), simple Mantel tests of the phylogenetic signal in species interactions for fungi and orchids revealed a significant but low phylogenetic signal (R<0.10) on the orchid side using Jaccard distances; however, the significance disappeared with UniFrac distances (Table S4). Similarly, marginally not-significant and low phylogenetic signals were detected on the mycorrhizal fungi side (R<0.04; Table S4). Next (step 2), results were qualitatively similar when using Mantel tests with permutations keeping the number of partners constant, suggesting that the phylogenetic signal in species interaction did not result from a phylogenetic signal in the number of partners. Our investigation of clade-specific phylogenetic signals in species interactions in orchids (option 1) revealed a significant phylogenetic signal in Angraecineae, a sub-tribe composed of 34 epiphytic species (Mantel test: R=0.37; Bonferroni-corrected p-value=0.016; Fig. 5) interacting with 53 fungi, suggesting that closely related Angraecineae tend to interact with more similar mycorrhizal fungi. When we checked the robustness of the significant phylogenetic signal detected in Angraecineae (option 2) by subsampling the Angraecineae clade down to 10 species, we still recovered a significant signal in species interactions (Fig. S32). Similarly, we still recovered a significant signal when changing the arbitrary age of the polytomies corresponding to unresolved orchid genera (Fig. S33).

**Figure 4:**
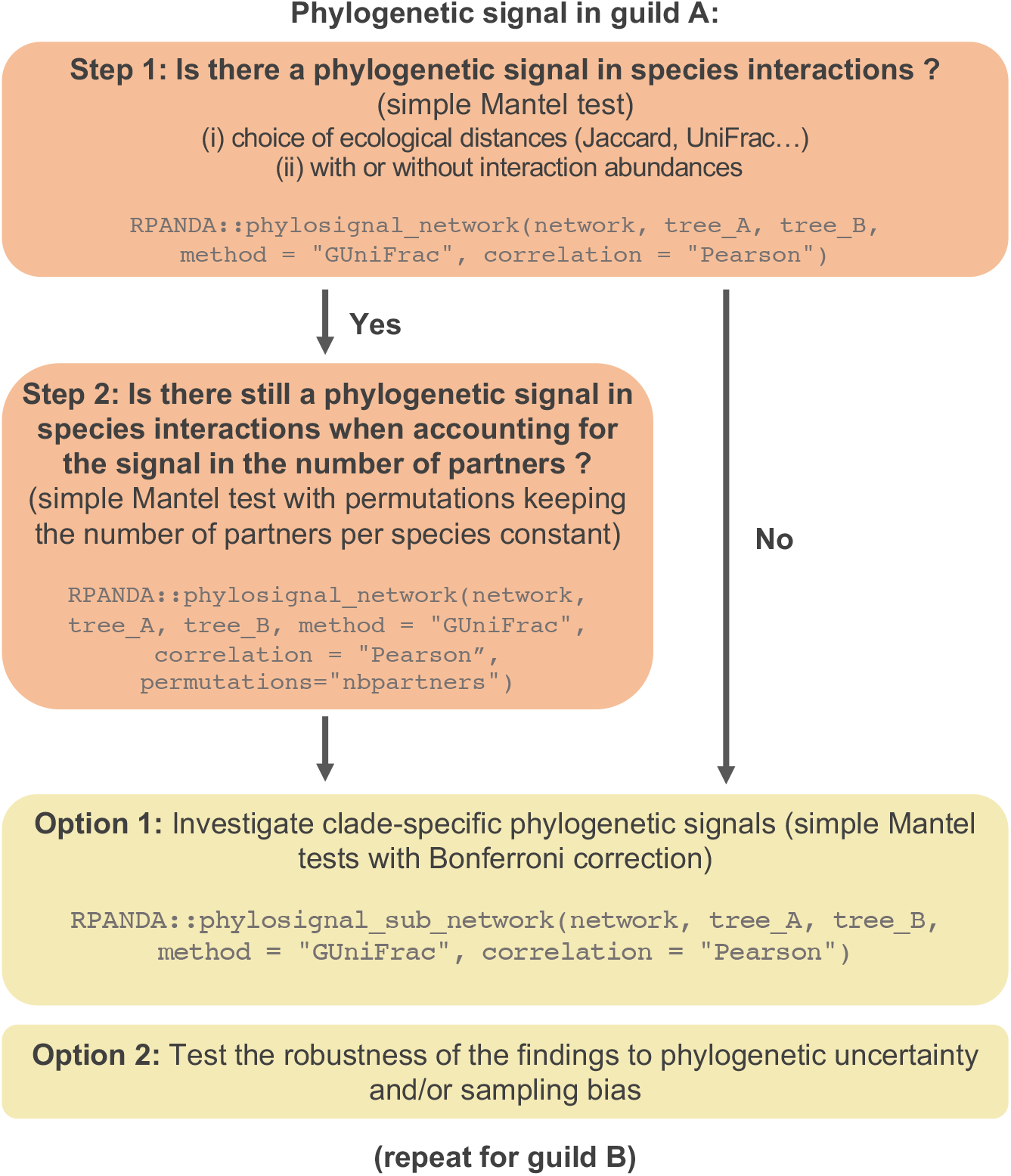
Recommended guidelines to measure phylogenetic signal in species interactions within bipartite ecological networks. This guideline is composed of two fixed steps followed by two optional ones and can be applied as soon as a bipartite interaction network (with or without abundances) and at least the phylogenetic tree of guild A are available. The phylogenetic tree does not need to be binary, rooted, or ultrametric. For each step, an example of the corresponding function available in the R-package RPANDA is indicated in grey. **Step 1:** The first step consists of testing for a phylogenetic signal in species interactions for guild A (*i*.*e*. whether closely related species from guild A tend to interact with similar partners from guild B) using a one-tailed simple Mantel test. This step requires picking an ecological distance (*e*.*g*. UniFrac or Jaccard distances) and a type of correlation (Pearson correlation by default). **Step 2:** Next, to assess whether a phylogenetic signal in species interactions really comes from the identity of species interactions (and not from a phylogenetic signal in the number of partners), the second step consists of testing whether the phylogenetic signal in guild A remains significant when the significance of the Mantel correlation is evaluated using network permutations keeping the number of partners constant. **Option 1:** Clade-specific phylogenetic signals in guild A can be tested using simple Mantel tests while correcting for multiple testing (*e*.*g*. Bonferroni correction). It can be used to test whether some clades present different intensities of phylogenetic signals (*e*.*g*. because of higher specificity). **Option 2:** The robustness of the findings can be tested by looking at how the conclusions might be affected by phylogenetic uncertainty (*e*.*g*. using a Bayesian posterior of tree) or sampling bias. The potential effect of sampling bias can be investigated by subsampling all clades to the same number of species. If a phylogenetic tree for guild B is available, all these steps can be replicated to test for the phylogenetic signal in species interaction in guild B.

**Figure 5:**
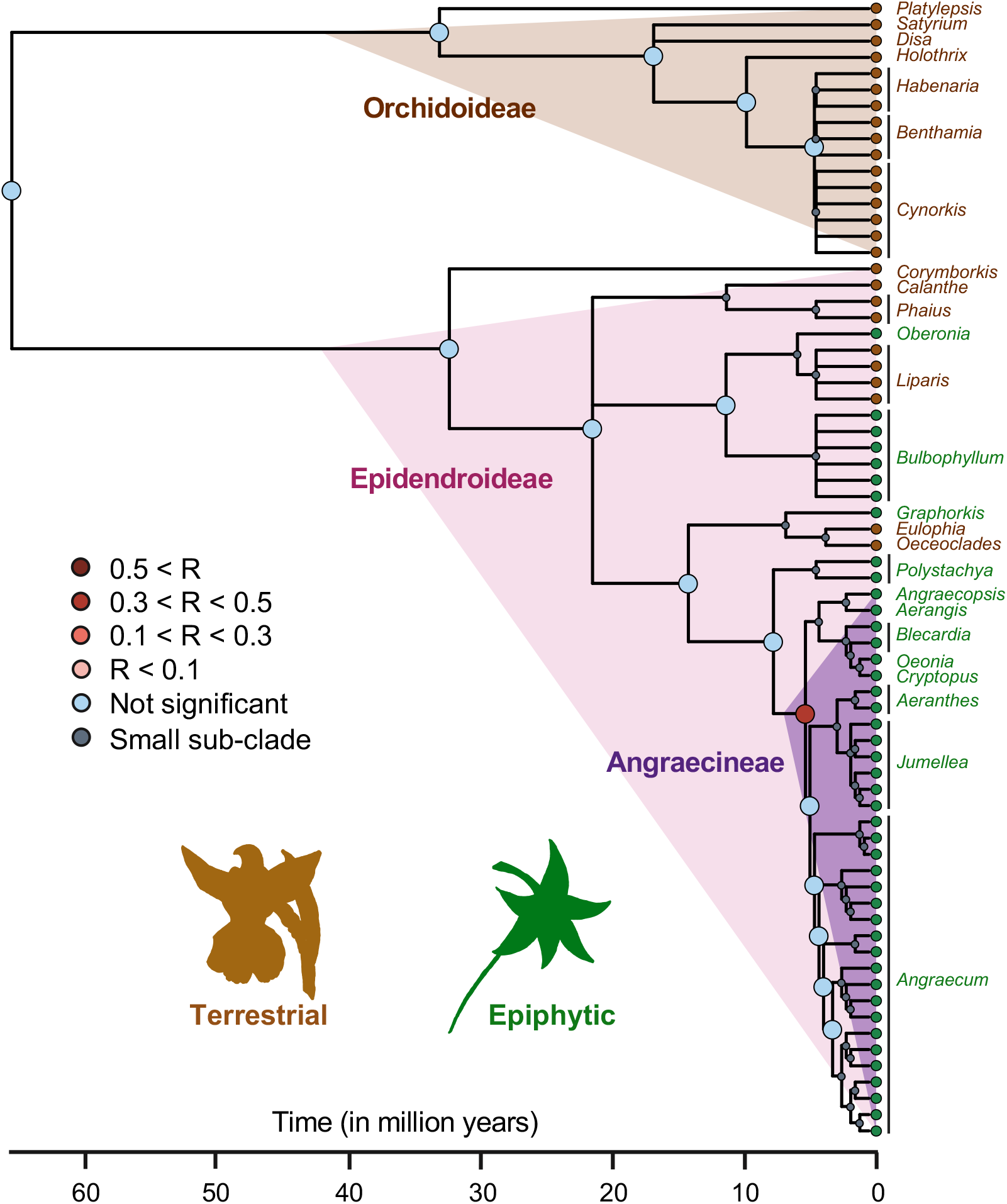
Empirical application on an orchid-fungus interaction network from La Réunion island (Martos *et al*., 2012): the clade-specific analyses of phylogenetic signals in species interactions revealed a significant phylogenetic signal in the epiphytic subtribe Angraecineae. The orchid phylogeny (Martos *et al*., 2012) is represented with its nodes colored according to the results of the Mantel test performed on the corresponding sub-network: in blue if non-significant, in grey when the node has less than 10 descendent species (the Mantel test was not performed), and in red when the phylogenetic signal is significant. Each one-tailed simple Mantel test was performed using the Pearson correlation, UniFrac distances, and 100,000 permutations, and its significance was evaluated while correcting for multiple testing (Bonferroni correction). For each species, its habitat (terrestrial or epiphytic) is indicated at the tips of the tree and the main orchid clades are highlighted in colors. Only the genera are indicated at the tips of the tree (see Supplementary Figure S32 for the species list).

## Discussion

We used simulations to perform a comparative analysis of the statistical performances of Mantel tests and the Phylogenetic bipartite linear model (PBLM; Ives & Godfray 2006) for testing for a phylogenetic signal in species interactions. Our results highlight the weaknesses of PBLM and partial Mantel tests and advocate for the use of regular Mantel tests and Mantel tests with network permutations keeping the number of partners per species constant.

The Phylogenetic bipartite linear model (PBLM) is widely used to test for phylogenetic signal in species interactions, however, we found that it has very frequent false positives (>30%). PBLM assumes that the interaction strength between two species is determined by the product of two unobserved traits evolving on the phylogenies of guilds A and B respectively, according to two independent Ornstein-Uhlenbeck processes with the selection strengths d_A_ and d_B_ (Supplementary Methods 3). PBLM tests the significance of d_A_ and d_B_, which measure the phylogenetic signal of the unobserved traits. A species with a high trait value will have high interaction strengths with many partner species (*i*.*e*. it is a generalist species), while a species with a low trait value will have low interaction strengths with most partner species, except with the few species with high trait values (*i*.*e* it is a specialist species). Therefore, we suspect that d_A_ and d_B_ measure phylogenetic signals in the number of partners rather than in species interactions. However, we also found significant d_A_ and d_B_ in the absence of phylogenetic signal in the number of partners, suggesting that PBLM is sensitive to model misspecification (it relies on strong hypotheses on how the number of partners evolves). In any case, our results suggest that PBLM should not be used as a routine for measuring phylogenetic signal in species interactions.

Other process-based approaches that extend PBLM (Hadfield et al., 2014; Li et al., 2020; Rafferty & Ives, 2013) allow inferring parameters thought to reflect the phylogenetic structure of interactions networks, while controlling for the phylogenetic signal in the number of partnerns as well as the heterogeneity in sampling effort (Hadfield *et al*., 2014). Our analyses using the PGLMM approach (Rafferty & Ives, 2013) on the smallest simulated networks suggested that it also has frequent false positives and intermediate statistical power when using weighted interactions. It would have been ideal to also test this approach on larger networks, but this was prohibited by their computational cost (Fig. S2). Indeed, fitting PGLMM can require >80 Gb of memory for some networks and our application of the Bayesian approach of Hadfield *et al*. (2014) ran several days (on an Intel 2.8 GHz, MacOSX laptop) without reaching convergence. Because of these high computational demands, these methods are typically not used to measure phylogenetic signal in species interactions in empirical studies, which is either done using Mantel tests or PBLM (see Fontaine and Thébault 2015; Xing et al. 2020; Corro et al. 2021 for recent examples). Future model developments of such approaches would thus benefit from faster inferences and our results highlight the need to thoroughly test these approaches with simulations before they are applied to empirical systems and biological conclusions are drawn.

We found that simple Mantel tests have a moderate statistical power and a reasonable false positive rate (<5%) when testing for phylogenetic signal in species interactions. Not surprisingly, these tests have a higher power for larger simulated networks. The fact that Mantel tests have a moderate power for measuring phylogenetic signal in species interactions corroborates the findings about Mantel tests in other contexts (Guillot & Rousset, 2013; Harmon & Glor, 2010). Hence, although simple Mantel tests might fail at detecting low phylogenetic signals, we can trust their results when they are significant. On the contrary, we found a high proportion of simulated networks (5-10%) presenting a significant negative phylogenetic signal in species interactions, suggesting that closely related species would tend to associate with dissimilar partners. Yet, we did not expect such an outcome in our simulations because we did not observe any negative phylogenetic signal in species traits. False positives are therefore frequent when testing for a negative phylogenetic signal using simple Mantel tests and detection of such signals in empirical networks should be interpreted with caution.

In addition, Pearson correlations performed better than Spearman and Kendall correlations, which is somewhat surprising, as correlations between phylogenetic and ecological distances are not particularly expected to be linear: Spearman and Kendall correlations have less stringent hypotheses, as they only assume monotonicity (Supplementary Methods 3), but they probably lose information. We also reported that using ecological distances that consider interaction abundances, such as weighted Jaccard or UniFrac distances, significantly improves the detection of phylogenetic signals. Using UniFrac distances, which rely on the phylogenetic relatedness of the partners, can be particularly relevant when species delineation is somewhat arbitrary, *e*.*g*. in microbial systems, as it is less sensitive to species delineation than Jaccard distances. In addition, results obtained with UniFrac distances were only moderately influenced by the phylogenetic uncertainty in the partner’s tree, which should thus not prevent the use of UniFrac distances. In the context of our *BipartiteEvol* simulations, which assume that species interactions are mediated by some phylogenetically-conserved traits on both sides of the network, we found that UniFrac distances outperform Jaccard distances.

We note however that a significant phylogenetic signal in UniFrac or Jaccard distances can reflect different evolutionary processes, such as one where the traits involved in the interaction are evolutionarily conserved on both sides of the networks in the case of UniFrac, and on only one side of the network in the case of Jaccard (Calatayud et al., 2016). Therefore, choosing between one or the other metric (or using both) can also be dictated by the question at stake. Also, if communities of interactors differ mainly in terms of recently diverged species, Jaccard distances may perform better, as UniFrac distances emphasize differences in long branches rather than recent splits (Sanders et al., 2014).

We also found that multiple simple Mantel tests combined with a Bonferroni correction perform rather well to investigate clade-specific phylogenetic signals. Such an approach can therefore be valuable for detecting the phylogenetic signals in particular sub-clades among large “meta-networks”, such as those describing host-microbiota phylosymbiosis (Song et al., 2020), which likely have heterogeneous phylogenetic signals across the network.

While simple Mantel tests have satisfactory statistical performances, these tests do not control for the potential confounding effect of the phylogenetic signal in the number of partners. Partial Mantel tests are frequently used for investigating a phylogenetic signal in species interactions while controlling for the signal in the number of partners; however, we found that they often detected significant signals in species interactions when we simulated signals in only the number of partners. Thus, partial Mantel tests fail at discerning whether evolutionary relatedness strictly affects the identity of partners, independently of the total number of partners associated with each species (Rezende et al., 2007). This corroborates the poor statistical performances of partial Mantel tests frequently observed in other contexts (Guillot & Rousset, 2013; Harmon & Glor, 2010). Among the alternative possibilities we tested, using sequential Mantel tests, *i*.*e*. testing first for the phylogenetic signal in species interactions, and if significant, testing for the phylogenetic signal in the number of partners, has both high statistical power and a low false positive rate. Yet, if both Mantel tests are significant, it does not say whether the signal is entirely due to the signal in the number of partners and therefore, sequential Mantel tests likely have very low power in this case. Alternatively, using methods that can explicitly partition ecological distances into parts due to dissimilarities in the number of partners *versus* the identity of the partners appears promising, although we detected a slight power decrease in our simulations and >30% of artefactual negative phylogenetic signals when partitioning unweighted Jaccard distances. Other partitioning approaches may give better results and should require further attention, as they offer a direct quantification of the contribution of the species identity *versus* the number of partners in the phylogenetic signal (Baselga, 2010; Calatayud et al., 2016; Leprieur et al., 2012). Finally, performing a Mantel test with network permutations designed to keep the number of partners associating with each species constant while shuffling their identity has infrequent false positives and does not decrease the statistical power. Therefore, if there is still a signal while constraining the number of partners, then we can safely conclude that evolutionary relatedness affects the identity of partners. We thus recommend using such network permutations to correct for the confounding effect of the phylogenetic signal in the number of partners (Figure 4).

By definition, phylogenetic signals in species interactions measure general patterns that are not informative of the processes at play (Losos, 2008). A better understanding of the ecological and evolutionary processes playing a role in the assembly of interaction networks (Harmon et al., 2019) will require developing integrative process-based approaches, for instance, an inference machinery for eco-evolutionary models such as *BipartiteEvol*. Classical inferences (generalized least-squares or likelihood-based approaches) might be challenging for such complex models (Hadfield et al., 2014), but strategies such as machine learning provide promising alternatives.

In the mycorrhizal network from La Réunion, we found non-significant or weak phylogenetic signals in species interactions at the level of the entire orchid-fungus network, suggesting these interactions are generally poorly conserved over long evolutionary timescales (Jacquemyn et al., 2011; Martos et al., 2012; Perez-Lamarque et al., 2022). Conversely, clade-specific Mantel tests detected a significant phylogenetic signal in the Angraecineae epiphytic clade that is experiencing a radiation on La Réunion island. This signal is likely produced by the different orchids genera in Angraecineae associated with specific fungal clades (Martos et al., 2012). Thus, our results corroborate a trend toward mycorrhizal specialization in epiphytic orchids compared with terrestrial species (Xing et al., 2019), as the epiphytic habitats might require particular adaptations and stronger dependences on specific mycorrhizal fungi.

Interaction networks are increasingly being analyzed to unravel the evolutionary processes shaping their structure and to predict their stability. Currently-used tools for measuring phylogenetic signals are clearly misleading. The approach we propose based on Mantel tests may have a limited statistical power, but it avoids false positives, and it is flexible as it allows using different ecological distances and/or permutation strategies. By emphasizing the limits of current tests of phylogenetic signal, we hope to stimulate new developments in the statistical adjustment to empirical data of process-based models for the evolution of interaction networks.

## Acknowledgements

The authors acknowledge M. Elias, E. Thébault, and D. de Vienne for helpful discussions. They also thank I. Overcast, S. Lambert, I. Quintero, C. Fruciano, J. Clavel, and A. Silva for comments on an early version of the manuscript. Preprint version 6 of this article has been peer-reviewed and recommended by Peer Community In Evolutionary Biology (https://doi.org/10.24072/pci.evolbiol.100150).

## Data, scripts and codes availability

The R functions used to measure phylogenetic signals in bipartite interaction networks, including (simple, partial, and clade-specific) Mantel tests and PBLM, are available in the R-package RPANDA (Morlon et al. 2016) (functions phylosignal_network and phylosignal_sub_network). A tutorial can be found at https://github.com/BPerezLamarque/Phylosignal_network. Amended functions of BipartiteEvol are also included in RPANDA. All generated data, scripts, and codes for simulating the networks and for measuring the phylogenetic signals in species interactions are publicly accessible through the Open Science Framework (osf) portal (https://doi.org/10.17605/OSF.IO/ZGM86).

## Supplementary material

Supplementary materials (including Supplementary Methods, Tables, and Figures) are available through the Open Science Framework (osf) portal (https://doi.org/10.17605/OSF.IO/ZGM86).

