## Supplementary material for "Do closely related species interact with similar partners? Testing for phylogenetic signal in bipartite interaction networks": Figure S

**Main manuscript:** Perez-Lamarque B, Maliet O, Pichon B, Selosse M-A, Martos F, Morlon H (2022) Do closely related species interact with similar partners? Testing for phylogenetic signal in bipartite interaction networks. bioRxiv, 2021.08.30.458192, ver. 6 peer-reviewed and recommended by Peer Community in Evolutionary Biology. https://doi.org/10.1101/2021.08.30.458192

**Supplementary Methods (1-3)**
**Supplementary Tables (1-6)**
**Supplementary Figures (1-33)**
**Supplementary References**

### Supplementary Methods:

#### Supplementary Methods 1: Expression of the fitness functions of interacting individuals from guilds A and B in the *BipartiteEvol* simulations

Each pair of interacting individuals from guilds A and B is characterized by six-dimensional traits ($x_A$ and $x_B$).

We considered that guilds A and B are both obligate partners, such that their fitness $W_A$ and $W_B$ can be written as a function of the parameters $\alpha_A$ and $\alpha_B$ for the mutualistic or neutral cases:

$$W_A(x_A, x_B) = e^{-\left\|x_A - x_B\right\|^2 \frac{\alpha_A{}^2}{2}} \qquad \text{(Eq. 1)}$$

$$W_B(x_A, x_B) = e^{-\left\|x_A - x_B\right\|^2 \frac{\alpha_B{}^2}{2}} \qquad \text{(Eq. 2)}$$

or for the antagonistic case:

$$W_A(x_A, x_B) = 1 - e^{-\left\|x_A - x_B\right\|^2 \frac{\alpha_A{}^2}{2}} \qquad \text{(Eq. 3)}$$

$$W_B(x_A, x_B) = e^{-\left\|x_A - x_B\right\|^2 \frac{\alpha_B{}^2}{2}} \qquad \text{(Eq. 4)}$$

See Figure 2 for an illustration of the mutualistic and neutral cases.

**Supplementary Methods 2: Comparing the structures of simulated and empirical networks:**

To compare the structures of the networks simulated using BipartiteEvol to the structures of empirical networks, we downloaded the Web of Life database (web-of-life.es (Fortuna *et al.*, 2014)) and the database of Michalska-Smith & Allesina (2019). These databases contain bipartite interaction networks at local or regional scales for a variety of antagonistic and mutualistic systems. We kept networks containing at least 10 species in each guild. We obtained a total of 334 empirical networks comprising bacteria-phage (antagonistic; n=17), seed dispersal (mutualistic; n=29), host-parasite (antagonistic; n=77), host-parasitoid (antagonistic; n=10), ant-plant (mutualistic; n=3), herbivory (antagonistic; n=23), anemone-fish (mutualistic; n=1), and pollination (mutualistic; n=174) networks.

We characterized the structures of each empirical or simulated network using three different metrics: the nestedness, the modularity, and the connectance (*i.e.* the ratio of realized interactions in the network). Each network was first converted into a binary matrix. We computed nestedness using the *NODFc* function correcting for network size (R-package maxnodf; Hoeppke & Simmons, 2021) and modularity using the function *computeModules* (R-package bipartite; Dormann *et al.*, 2008) with the "Beckett" method.

First, we separately compared these different metrics across empirical and simulated networks. Second, we jointly visualized them using principal component analyses (PCA). PCAs were performed with the functions *PCA* and *fviz_pca_ind* (FactoMineR and factoextra R-packages; Lê *et al.*, 2008).

**Supplementary Methods 3: Methods for measuring phylogenetic signal in species interactions:**

Mantel tests (Mantel, 1967) were introduced to examine the correlation between two dissimilarity matrices. They have been used to measure the phylogenetic signal in species interactions by computing the correlation between the matrix of phylogenetic distances (for species pairs from guild A for example, Figure 1) and the matrix of 'ecological dissimilarities' between the sets of interacting partners (*i.e.* species from guild B interacting with species pairs from guild A). The correlation (R) with $-1<R<1$ is often evaluated using Pearson correlation, that is the mean of the products of the corresponding elements in the two standardized dissimilarity matrices. Alternatively, R can be evaluated using Spearman correlation (computed from dissimilarities transformed into ranks), or Kendall correlation (computed by counting the pairs of observations that have the same rank). The parametric Pearson correlation is statistically more powerful, but makes stronger hypotheses (it assumes a linear relationship) than the non-parametric Spearman and Kendall correlations (which assume only a monotonic relationship). A positive (resp. negative) correlation indicates a positive phylogenetic signal (resp. negative phylogenetic signal) in species interactions. Its significance is evaluated using randomizations by repeatedly permuting one of the original dissimilarity matrices: one-tailed p-values are obtained by comparing the rank of the original correlation R to the randomized correlations. Ecological dissimilarities of interacting partners between two species from guild A can be measured with various indices. Two classical indices are the Jaccard distance, defined as the number of their unshared partners from guild B divided by their total number of partners, and the UniFrac distance, which incorporates phylogenetic relatedness between partners, computed as the fraction of unshared branch length in the phylogenetic tree of their partners from guild B. Both indices also have a weighted version that accounts for interaction strength (e.g. weighted Jaccard and generalized/weighted UniFrac; Chen *et al.* 2012).

Partial Mantel tests examine the correlation between two dissimilarities matrices while accounting for a third dissimilarity matrix (Smouse *et al.*, 1986). When testing for phylogenetic signal in bipartite interaction networks, these tests are useful for controlling the phylogenetic signal in the number of partners; indeed similarity in the number of partners can decrease the value of ecological dissimilarity metrics independently of the identity of the partners, such that a phylogenetic signal in the number of partners can generate a phylogenetic signal in species interactions that is not linked to an evolutionary conservatism of interacting partners. Partial Mantel tests therefore investigate the correlation between phylogenetic and ecological dissimilarities while controlling for the pairwise difference in the number of partners between species pairs (Rezende *et al.*, 2007).

The Phylogenetic bipartite linear model (PBLM; Ives & Godfray 2006) assumes that interaction strengths between species from guilds A and B are determined by (unobserved) traits that evolve on the two phylogenies each following a simplified Ornstein-Uhlenbeck process (Blomberg *et al.*, 2003). The strength of interaction between two species is assumed to be given by the product of their two traits. Under these assumptions, $d_A$ and $d_B$ can be estimated from the two phylogenies and the matrix of interaction strengths using generalized least squares (Ives & Godfray 2006). $d_A$ and $d_B$ are then interpreted as a measure of phylogenetic signal in species interactions. If d=1, the traits evolve as Brownian processes; if d=0, there is no effect of the phylogeny (similar as evolution on a star phylogeny, *i.e.* no phylogenetic signal); 0<d<1 generates stabilizing selection (*i.e.* phylogenetic signal) and d>1 disruptive selection (*i.e.* negative phylogenetic signal). Ives & Godfray (2006) proposed two approaches to assess the significance of the signal. The simplest consists of comparing the mean square errors (MSE) of the generalized least squares regression to the same MSE obtained using star phylogenies ($MSE_{star}$). $MSE<MSE_{star}$ is interpreted as a significant phylogenetic signal in species interactions. The second approach uses a bootstrapping strategy to build 95% confidence intervals around the estimated $d_A$ and $d_B$ values: the null hypothesis (absence of phylogenetic signal in guild A, resp. B) is rejected if the confidence interval around $d_A$ (resp. $d_B$) does not include 0. While designed primarily for applications to bipartite networks characterized by matrices of interaction strengths (*e.g.* net attack rate of a parasitoid on its hosts), PBLM has been applied to weighted networks characterized by matrices of interaction abundance (*i.e.* the number of times the interaction has been observed) and unweighted (binary) networks, using 1 for the interaction strength when species interact and 0 otherwise (Ives & Godfray, 2006; Vázquez *et al.*, 2009; Jacquemyn *et al.*, 2011; Martos *et al.*, 2012; Xing *et al.*, 2020).

The phylogenetic generalized linear mixed model (PGLMM; Rafferty and Ives 2013) uses linear mixed models to infer phylogenetic signals in both the number of partners and species interactions. In its general form, the model assumes that the abundance $Y_i$ of the pairwise interaction $i$ between one species from guild A and one species from guild B is given by:

$$Y_i = \mathrm{a} + \alpha_{A[i]} + b_{A[i]} + c_i + d_{B[i]} + f_{B[i]} + g_i + h_i + e_i$$

where a is the mean abundance of the interaction, $e_i$ the error term of the model, and the other terms correspond to random effects. $\alpha_{A[i]}$ (resp. $d_{B[i]}$ ) models the variation of the abundances between species from guild A (resp. B). $b_{A[i]}$ (resp. $f_{B[i]}$) models the variation of the total abundances of interactions between species from guild A (resp. B) that can be explained by the phylogenetic history of guild A (resp. B); it thus corresponds to a measure of the phylogenetic signal in the total number of partners (considering the abundances of the interactions) in guild A (resp. B). $c_i$ (resp. $g_i$) models whether closely related species from guild A (resp. B) tend to interact with similar species from guild B; it thus corresponds to a measure of the phylogenetic signal in species interactions in guild A (resp. b). Finally, $h_i$ models whether closely related species from guild A tend to interact with closely related species from guild B (i.e. a "cophylogenetic signal"). If the abundances do not follow a Gaussian distribution, generalized linear mixed models can be used to fit the model with a Poisson distribution (Li *et al.*, 2020). Similarly, if only presence/absence data are available (*i.e.* unweighted interaction networks), generalized linear mixed models can be used with a binomial distribution (Li *et al.*, 2020).

Following Lajoie & Kembel (2021), to evaluate the significance of the different parameters of the models, one can fit models of increasing complexity in a stepwise framework and evaluate the support of the models using likelihood ratio tests.

### Supplementary Tables:

**Supplementary Table 1: False positive rate of simple Mantel tests measuring phylogenetic signal in species interactions evaluated using *BipartiteEvol* simulations (Maliet *et al.*, 2020). False positive rates vary according to the ecological distance, the correlation, and the network sizes.**

Each grid indicates the percentage of simulated networks that present a significant phylogenetic signal in guilds A or B (*i.e.* p-value < 0.05 for the test of phylogenetic signal) while no phylogenetic signal is expected (false positive). These networks correspond to networks simulated under neutrality ($\alpha_A$=0; $\alpha_B$=0) or to mutualistic or antagonistic networks that present no phylogenetic signal in species traits (see Supplementary Figure 5). Expectations are based on Mantels tests for measuring the phylogenetic signals in species traits (but results are qualitatively similar when using Pagel's λ instead; Supplementary Figure 11).

One-tailed Mantel tests were performed using 10,000 permutations. See Table S5 for the number of simulations per category of network sizes.

Bolded values are false positive rates larger than 5%.

| Ecological distance | Correlation | Network size | False positive rate in guild A | False positive rate in guild B |
|---|---|---|---|---|
| Weighted Jaccard | Pearson | <150 | **0.081** | **0.062** |
| | | 150-250 | 0.038 | 0.020 |
| | | >250 | 0.038 | 0.018 |
| | Spearman | <150 | 0.022 | 0.023 |
| | | 150-250 | 0.013 | 0.007 |
| | | >250 | 0.027 | 0.011 |
| | Kendall | <150 | 0.018 | 0.012 |
| | | 150-250 | 0.009 | 0.010 |
| | | >250 | 0.017 | 0.011 |
| Unweighted Jaccard | Pearson | <150 | 0.044 | 0.035 |
| | | 150-250 | 0.031 | 0.013 |
| | | >250 | 0.031 | 0.011 |
| | Spearman | <150 | 0.018 | 0.023 |
| | | 150-250 | 0.016 | 0.010 |
| | | >250 | 0.020 | 0.011 |
| | Kendall | <150 | 0.018 | 0.023 |
| | | 150-250 | 0.016 | 0.010 |
| | | >250 | 0.024 | 0.014 |
| Weighted UniFrac | Pearson | <150 | 0.029 | 0.046 |
| | | 150-250 | 0.031 | 0.030 |
| | | >250 | 0.041 | 0.025 |
| | Spearman | <150 | 0.026 | 0.019 |
| | | 150-250 | 0.031 | 0.020 |
| | | >250 | 0.041 | 0.028 |
| | Kendall | <150 | 0.018 | 0.019 |
| | | 150-250 | 0.025 | 0.017 |
| | | >250 | 0.034 | 0.025 |
| Unweighted UniFrac | Pearson | <150 | 0.037 | 0.054 |
| | | 150-250 | 0.044 | 0.023 |
| | | >250 | 0.034 | 0.021 |
| | Spearman | <150 | 0.026 | 0.031 |
| | | 150-250 | 0.041 | 0.027 |
| | | >250 | 0.038 | 0.025 |

| | | | | |
|---|---|---|---|---|
| | Kendall | <150 | 0.022 | 0.039 |
| | | 150-250 | 0.025 | 0.017 |
| | | >250 | 0.031 | 0.021 |

**Supplementary Table 2: Statistical power of simple, partial, and sequential Mantel tests, measured as the percentage of *BipartiteEvol* networks with a detected phylogenetic signal when expected (Maliet *et al.*, 2020).**

As a reminder, in simulations using *BipartiteEvol*, we expect a significant phylogenetic signal in species interactions in networks simulated with an effect of the traits on individual fitness (antagonistic and mutualistic simulations) and when traits are phylogenetically conserved (see Supplementary Figure 5). Expectations are based on Mantels tests for measuring the phylogenetic signals in species traits (but results are qualitatively similar when using Pagel's λ instead; Supplementary Figure 11).

Each grid indicates the percentage of networks for which we expect phylogenetic signal that actually present a significant Pearson correlation between ecological distances (measuring using Jaccard or UniFrac distances and using weighed or unweighted interactions) and phylogenetic distances in guilds A and B, using different one-tailed Mantel tests: (i) a simple Mantel test between the two distance matrices, (ii) a partial Mantel test controlling for the differences in numbers of partners (pairwise differences in the number of partners), and (iii) two sequential simple Mantel tests – a first one tests the correlation between ecological distances and phylogenetic distances, and a second one tests the correlation between differences in numbers of partners and phylogenetic distances: thus, they control that if there is a phylogenetic signal in species interactions, it is not explained by a phylogenetic signal in the number of partners.

| | (i) statistical power of the simple Mantel test | | (ii) statistical power of the partial Mantel test | | (iii) statistical power of the sequential Mantel tests | |
|---|---|---|---|---|---|---|
| **Ecological distances** | **guild A** | **guild B** | **guild A** | **guild B** | **guild A** | **guild B** |
| Weighted Jaccard | 39.3% | 34.6% | 37.9% | 33.8% | 37.2% | 32.9% |
| Unweighted Jaccard | 31.2% | 27.9% | 30.8% | 27.2% | 29.6% | 26.8% |
| Weighted UniFrac | 43.4% | 43.1% | 43.2% | 42.6% | 41.1% | 41.7% |
| Unweighted UniFrac | 35.6% | 35.1% | 34.9% | 34.1% | 34.0% | 33.7% |

| | (iv) statistical power of the Mantel test with beta diversity partitioning | | (v) statistical power of the Mantel test with permutations keeping constant the number of partners | |
|---|---|---|---|---|
| **Ecological distances** | **guild A** | **guild B** | **guild A** | **guild B** |
| Weighted Jaccard | NA | NA | 45.7% | 44.6% |
| Unweighted Jaccard | 25.6% | 23.8% | 46.3% | 45.5% |
| Weighted UniFrac | NA | NA | 44.4% | 43.4% |
| Unweighted UniFrac | NA | NA | 40.8% | 38.9% |

**Supplementary Table 3: Percentage of networks simulated with evolutionary conservatism in the number but not the identity of partners for which simple, partial, and sequential Mantel tests detect a significant phylogenetic signal in species interactions. These percentages reflect the false positive rate of the different tests.**

Each grid indicates the percentage of networks simulated with phylogenetic signal in the number of partners of guild A (according to an Ornstein-Uhlenbeck process ($a_A$>0) or a Brownian motion ($a_A$=0)) that present a significant correlation between ecological distances (measured using unweighted Jaccard or UniFrac) and phylogenetic distances in guilds A and B, using different one-tailed Mantel tests: (i) a simple Mantel test between the two distance matrices, (ii) a partial Mantel test controlling for differences in numbers of partners (pairwise differences in the number of partners), and (iii) two sequential simple Mantel tests – a first one tests the correlation between ecological distances and phylogenetic distances, and a second one tests the correlation between differences in numbers of partners and phylogenetic distances: thus, they control that if there is a phylogenetic signal in species interactions, it is not explained by a phylogenetic signal in the number of partners.

**(a) for all the simulated networks (with $a_A$>0 or $a_A$=0)**

| Ecological distances | Unweighted Jaccard | Unweighted UniFrac |
|---|---|---|
| (i) false positive rate of the simple Mantel test | 33.0% | 37.4% |
| (ii) false positive rate of the partial Mantel test | 29.2% | 25.6% |
| (iii) false positive rate of the sequential Mantel tests | 7.0% | 7.6% |
| (iv) false positive rate of the Mantel test with beta diversity partitioning | 3.8% | NA |
| (v) false positive rate of the Mantel test with permutations keeping constant the number of partners | 3.6% | 6.6% |

**(b) for only the simulated networks presenting a significant signal in the number of partners**

| Ecological distances | Unweighted Jaccard | Unweighted UniFrac |
|---|---|---|
| (i) false positive rate of the simple Mantel test | 41.5% | 47.6% |
| (ii) false positive rate of the partial Mantel test | 35.4% | 29.4% |
| (iii) false positive rate of the sequential Mantel tests | 0% | 0% |
| (iv) false positive rate of the Mantel test with beta diversity partitioning | 1.0% | NA |
| (v) false positive rate of the Mantel test with permutations keeping constant the number of partners | 3.5% | 7.7% |

**Supplementary Table 4: Phylogenetic signal in the entire orchid-fungus mycorrhizal network from La Réunion evaluated using simple Mantel tests with different ecological distances:**
One-tailed Mantel tests were performed using the Pearson correlation and 10,000 permutations. Significant positive or negative correlations (*i.e.* significant phylogenetic signals) are highlighted in bold.

Given that Mantel tests using weighted UniFrac distances have the lowest false positive rate and the highest power according to our simulations, there is likely no phylogenetic signal in this network. This discrepancy can be due to the fact that UniFrac distances tend to mainly weight the differences in long internal branches, whereas Jaccard distances can be more sensitive to recent "fungal speciations" (Sanders *et al.*, 2014). Thus, the detection of a phylogenetic signal using Jaccard distances could be the result of an over-splitting of fungal species.

| Ecological distances | Orchids | | | Fungi | | |
|---|---|---|---|---|---|---|
| | Pearson correlation | P-value for phylogenetic signal | P-value for negative phylogenetic signal | Pearson correlation | P-value for phylogenetic signal | P-value for negative phylogenetic signal |
| Weighted Jaccard | **0.096** | 0.0006 | 0.999 | **0.029** | 0.032 | 0.97 |
| Unweighted Jaccard | **0.094** | 0.0007 | 0.999 | **0.032** | 0.028 | 0.97 |
| Weighted UniFrac | -0.032 | 0.65 | 0.35 | 0.013 | 0.22 | 0.78 |
| Unweighted UniFrac | -0.046 | 0.74 | 0.26 | 0.014 | 0.20 | 0.80 |

**Supplementary Table 5:** Number of networks simulated using *BipartiteEvol* (Maliet *et al.,* 2020) in each size category (total number of species in guilds A and B) according to the different types of simulations (antagonism, mutualism, or neutral – with different sets of parameters). The discrete categories (<150 species; 150-250 species; and >250 species) have been chosen to have the most even number of simulations per category.

| Types | <150 species | 150-250 species | >250 species |
|---|---|---|---|
| Antagonism (i) | 31 | 41 | 48 |
| Antagonism (ii) | 32 | 41 | 47 |
| Antagonism (iii) | 35 | 41 | 44 |
| Antagonism (iv) | 38 | 41 | 41 |
| Antagonism (v) | 32 | 37 | 51 |
| Antagonism (vi) | 34 | 43 | 43 |
| Antagonism (vii) | 34 | 40 | 47 |
| Antagonism (viii) | 39 | 41 | 40 |
| Antagonism (ix) | 31 | 45 | 44 |
| Mutualism (i) | 69 | 49 | 2 |
| Mutualism (ii) | 37 | 42 | 41 |
| Mutualism (iii) | 32 | 41 | 47 |
| Mutualism (iv) | 40 | 69 | 11 |
| Mutualism (v) | 38 | 47 | 35 |
| Mutualism (vi) | 35 | 43 | 42 |
| Neutral | 158 | 201 | 241 |

**Supplementary Table 6:** Number of networks simulated using *BipartiteEvol* (Maliet *et al.*, 2020) in each size category (total number of species in guilds A and B) according to the different types of simulations (antagonism, mutualism, or neutral – with different sets of parameters) **when testing for the network asymmetry** (down to 10% of the species from guild A only were subsampled). The discrete categories (<80 species; 80-140 species; and >140 species) have been chosen to have the most even number of simulations per category.

| Types | <80 species | 80-140 species | >140 species |
|---|---|---|---|
| Antagonism (i) | 47 | 65 | 8 |
| Antagonism (ii) | 37 | 56 | 27 |
| Antagonism (iii) | 42 | 44 | 34 |
| Antagonism (iv) | 41 | 58 | 21 |
| Antagonism (v) | 32 | 52 | 36 |
| Antagonism (vi) | 51 | 64 | 5 |
| Antagonism (vii) | 41 | 42 | 37 |
| Antagonism (viii) | 55 | 60 | 5 |
| Antagonism (ix) | 39 | 54 | 27 |
| Mutualism (i) | 57 | 63 | 0 |
| Mutualism (ii) | 31 | 60 | 29 |
| Mutualism (iii) | 36 | 46 | 38 |
| Mutualism (iv) | 31 | 59 | 30 |
| Mutualism (v) | 34 | 49 | 37 |
| Mutualism (vi) | 34 | 47 | 39 |
| Neutral | 177 | 225 | 198 |

### Supplementary Figures:

### Simulating phylogenetic signal in species interactions using *BipartiteEvol* (Maliet *et al.,* 2020):

**Supplementary Figure 1: Measuring the variability in the species trees simulated using *BipartiteEvol* (Maliet *et al.*, 2020) due to non-monophyletic species delineations.**

Because we amended the species delineation of the original *BipartiteEvol* model (Maliet *et al.*, 2020) and considered that each combination of traits forms a new species, species do not always result in a monophyletic group of individuals. Thus, we investigated here that the non-monophyletic species delineation gave coherent species trees by extracting two species trees per genealogy by randomly selecting one individual per species and verified the correlation between the two phylogenetic distance matrices using a one-tailed Mantel test (Pearson correlation with 10,000 permutations).

Here, we represent the Pearson correlations (R) for each of the 2,400 simulated networks. Panel (a) (resp. (b)) represents the Pearson correlation between pairs of simulated phylogenetic trees of the guild A (resp. B). The panels detailed the results per set of parameters ($\alpha_A$ and $\alpha_B$) and according to the size of the phylogenetic trees (<70, 70-100, or >100 species).

The large majority of the phylogenetic trees were identical (R=1) or very similar (R very close to one in few case) suggesting that most species were actually monophyletic and confirmed that our simulations generated consistent species trees. Only a very few trees (mostly corresponding to small mutualistic networks) have a correlation R<0.9, suggesting that some species are indeed not monophyletic, but the correlations remains larger than 0.65, suggesting that it does not deeply affect the tree topologies (and might therefore only marginally affect our results).

In addition, this change in the species delineation did not affect the structural properties in the simulated networks, that present significant nestedness and/or modularity according to the sets of simulated parameters (results not shown).

Boxplots present the median surrounded by the first and third quartiles, and whiskers extend to the extreme values but no further than 1.5 of the inter-quartile range.

(a)

(b)

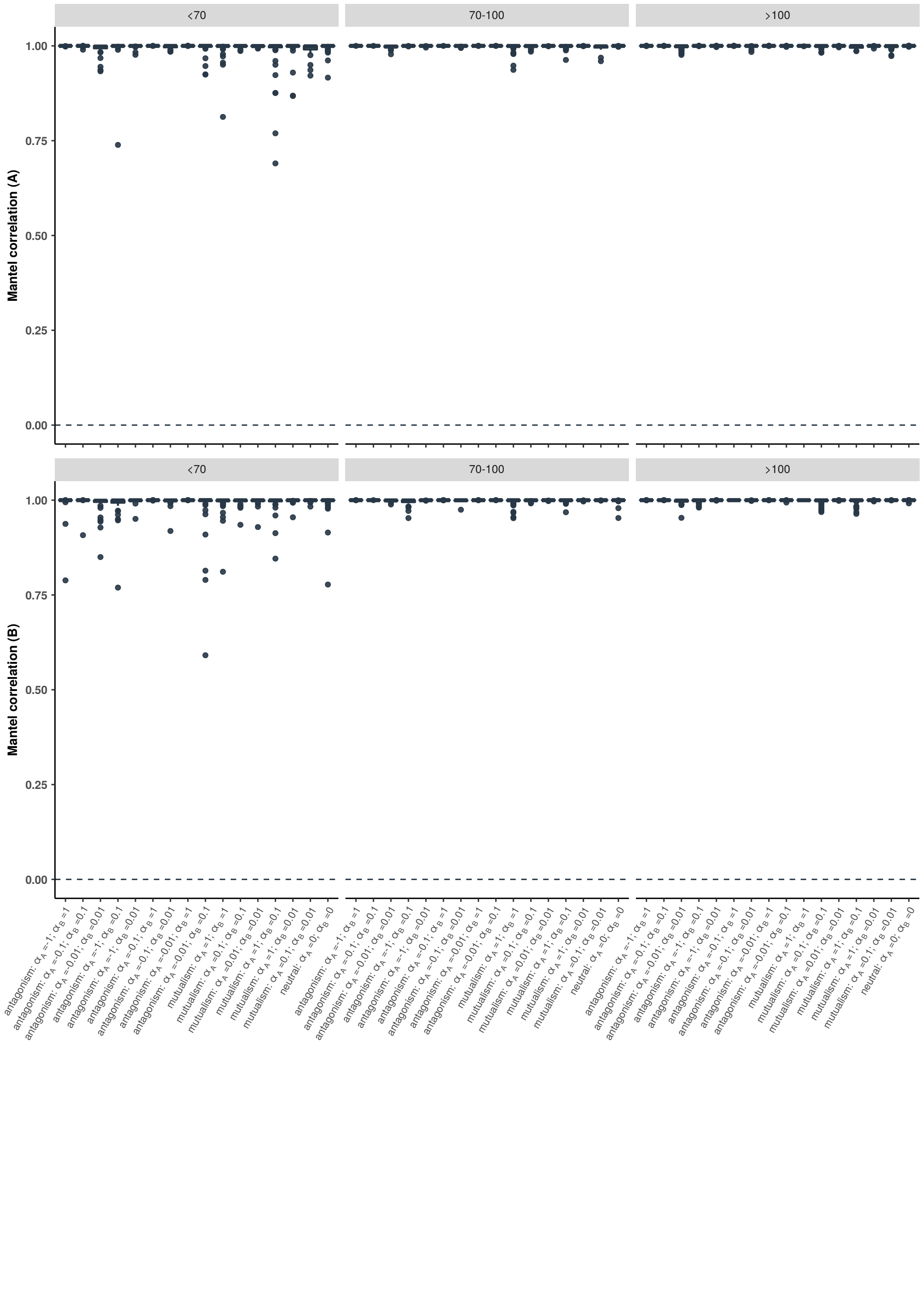

<70
70-100
>100
Mantel correlation (A)
Mantel correlation (B)
1.00
0.75
0.50
0.25
0.00

**Supplementary Figure 2: Computation time of the different approaches for evaluating phylogenetic signals in species interactions:**

For 10 interaction networks simulated using *BipartiteEvol* (Maliet *et al.*, 2020) randomly sampled among the small networks simulated with 500 pairs of individuals, we reported here the computation time (in second) on an Intel 2.8 GHz MacOSX laptop of different approaches for measuring phylogenetic signal in species interactions: (i) simple Mantel tests with weighted Jaccard or UniFrac distances, (ii) simple Mantel tests with weighted Jaccard or UniFrac distances and with permutations designed to keep constant the number of partners, (iii) PBLM, and (iv) PGLMM. Simple Mantel tests were run with Pearson correlations and 10,000 permutations, while Mantel tests keeping constant the number of partners were run with 1,000 permutations only.

Note that the computation time of PBLM and PGLMM tend to increase exponentially with the network size. PBLM and PGLMM also require more memory than the Mantel tests (not shown). Boxplots present the median surrounded by the first and third quartiles, and whiskers extend to the extreme values but no further than 1.5 of the inter-quartile range. The "y-axis" is log-scaled.

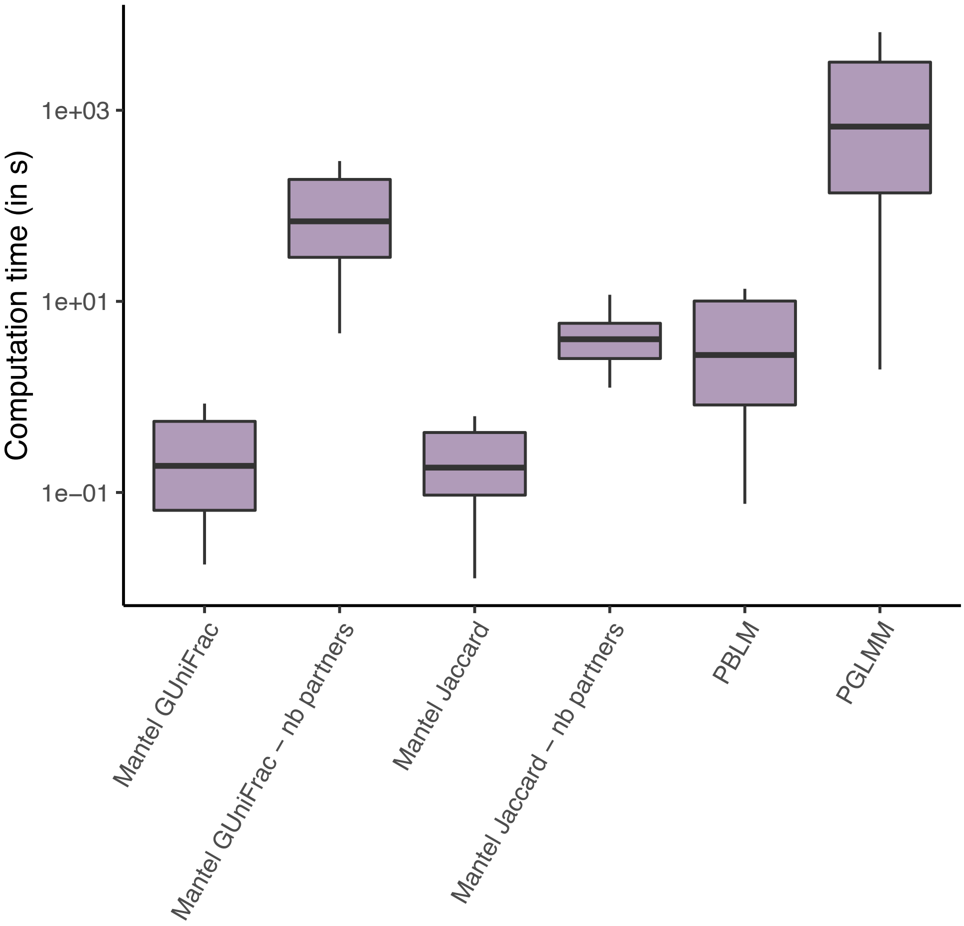

**Supplementary Figure 3: Networks simulated using *BipartiteEvol* (Maliet *et al.*, 2020) covered a large range of sizes.**

Numbers of species in guilds A (a) and B (b) as a function of the set of parameters ($\alpha_A$ and $\alpha_B$), and colored according to the total number of pairs of individuals for each guild (from 500 to 5,000).

Boxplots present the median surrounded by the first and third quartiles, and whiskers extend to the extreme values but no further than 1.5 of the inter-quartile range.

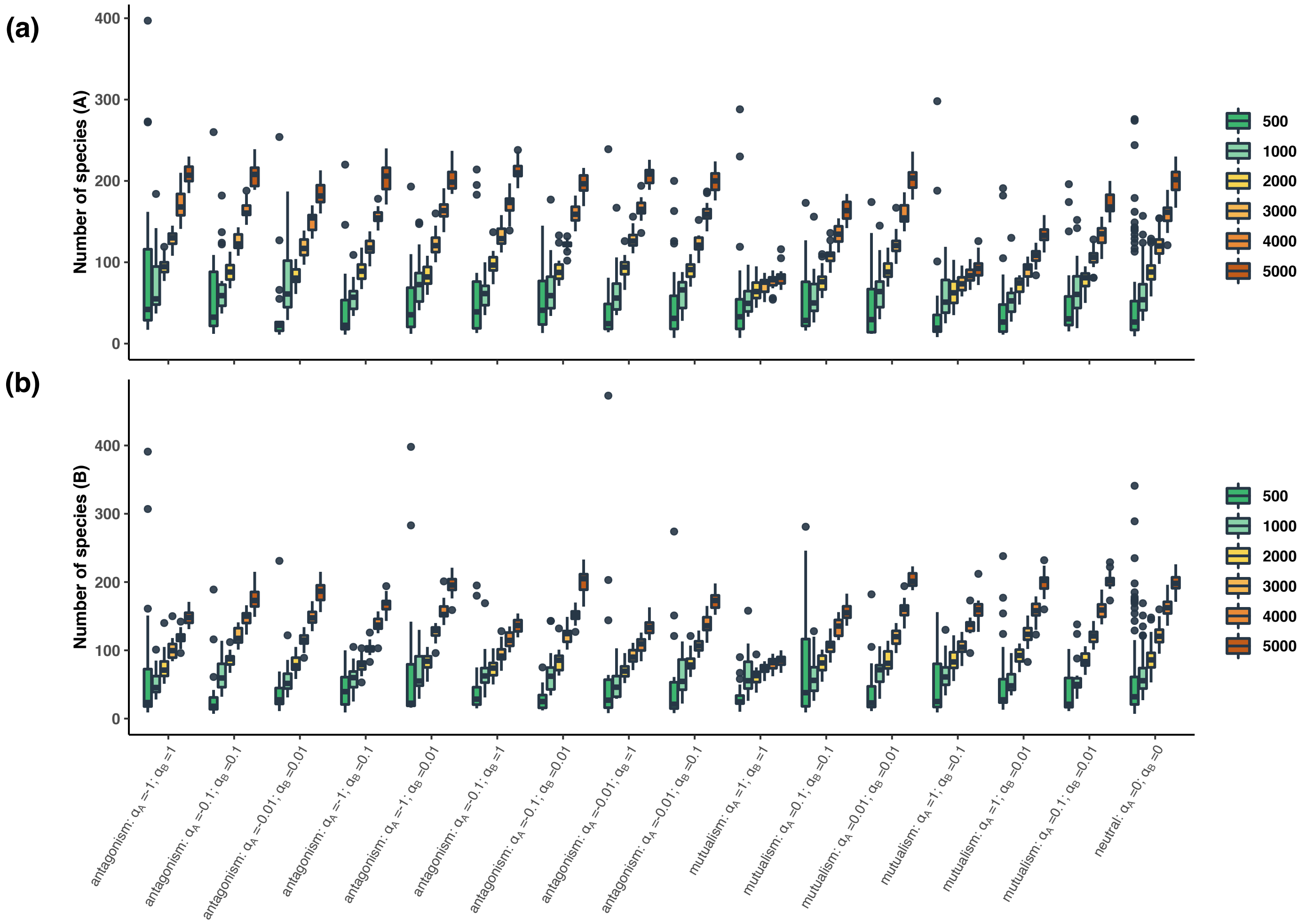

**Supplementary Figure 4: Networks simulated using *BipartiteEvol* (Maliet *et al.*, 2020) covered a realistic and large range of structures compared to empirical networks.**

For each BipartiteEvol simulated network (with neutral, mutualistic, or antagonistic sets of parameters) and each empirical (mutualistic or antagonistic) network, boxplots indicate the network connectance **(A)**, nestedness **(B)**, and modularity **(C)**. They present the median surrounded by the first and third quartiles, and whiskers extend to the extreme values but no further than 1.5 of the inter-quartile range. The connectances of the BipartiteEvol simulated networks tend to be lower than the connectances of the empirical networks, which is explained by the fact that empirical networks often miss rare species as an effect on undersampling. When we only sampled the 10% most abundant species (which represented a very large fraction of the total interactions), we obtained networks with connectances similar to those of empirical networks (Supplementary Figure 28). Our tests were performed on both full and subsampled networks.

**(D)** Projection of the empirical or simulated networks on the two principal components (PC1 and PC2) obtained using principal coordinate analysis (PCA) on the connectance, nestedness, and modularity of each network. For facilitating the visualization of the figure, we randomly sampled a subset of the BipartiteEvol simulated networks (we sampled 150 neutral, mutualistic, or antagonistic networks).
Simulated and empirical networks belong to the same cluster, although simulated networks are more clustered than empirical ones, suggesting that BipartiteEvol simulations have structures comparable to empirical ones although they do not fully replicate their full heterogeneity.

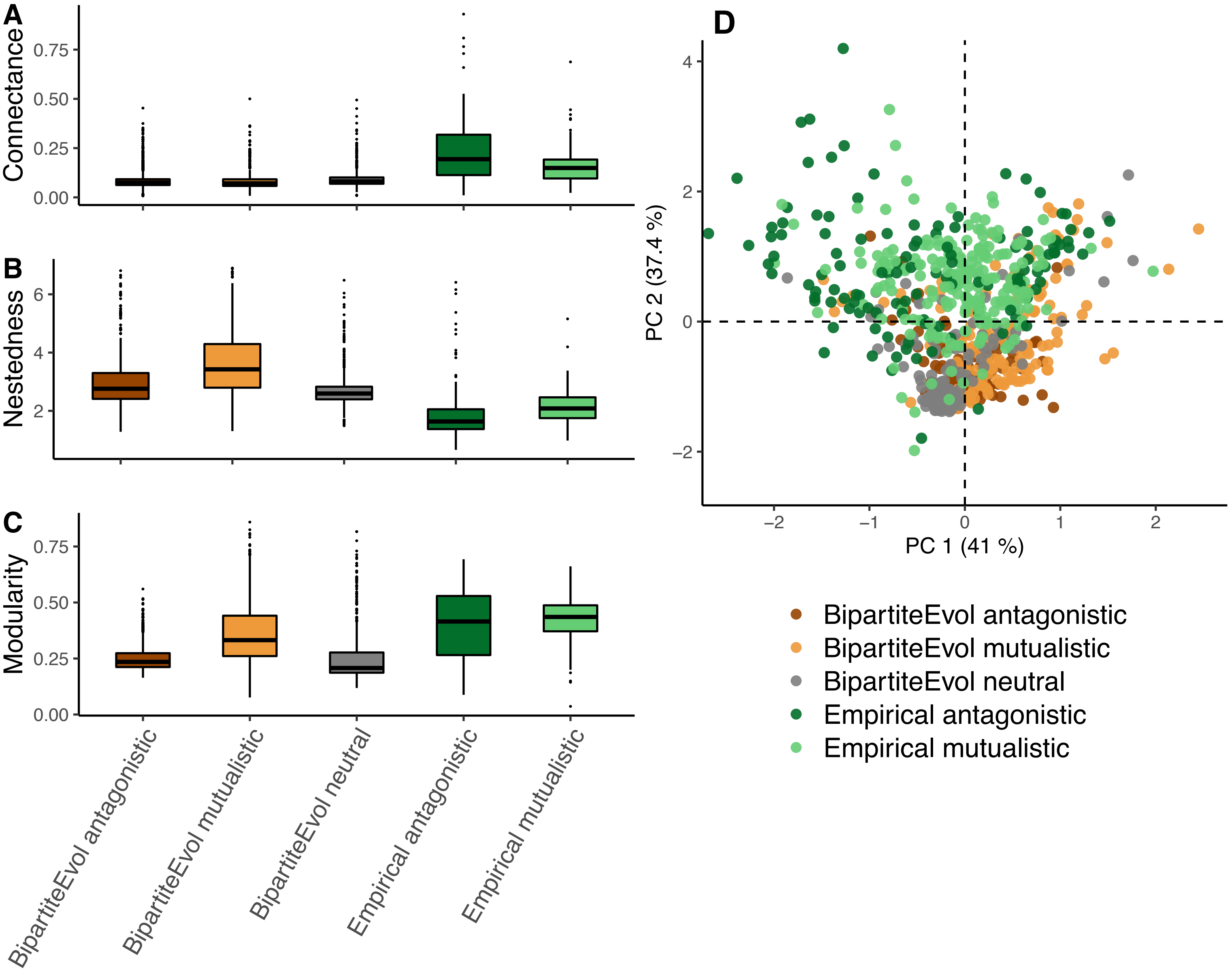

A
Connectance
0.00
0.25
0.50
0.75
B
Nestedness
2
4
6
C
Modularity
0.00
0.25
0.50
0.75
BipartiteEvol antagonistic
BipartiteEvol mutualistic
BipartiteEvol neutral
Empirical antagonistic
Empirical mutualistic
D
PC 2 (37.4 %)
PC 1 (41 %)
−2
−1
0
1
2
4
BipartiteEvol antagonistic
BipartiteEvol mutualistic
BipartiteEvol neutral
Empirical antagonistic
Empirical mutualistic

**Supplementary Figure 5: Closely-related species tend to have more similar traits with most sets of parameters in the *BipartiteEvol* simulations (Maliet *et al.*, 2020) based on Mantel tests (A) and Pagel's λ (B):**

**(A)** We first tested the phylogenetic signal in species' traits using Mantel tests. For each simulated network, panel (a) (resp. (b)) represents the results of the Mantel test between the phylogenetic distance matrix and the trait distance matrix for the guild A (resp. B). Each element of the trait distance matrix was computed as the pairwise Euclidian distance between the trait values of two species ($i$ and $j$) $|x_i - x_j|$. For each combination of parameters ($\alpha_A$ and $\alpha_B$), the bar indicates the percentage of simulated networks that present a significant positive correlation (in green; p-value<0.05 for the test of phylogenetic signal), a significant negative correlation (in red; p-value<0.05 for the test of negative phylogenetic signal), or no significant correlation (in yellow; both p-values>0.05). One-tailed Mantel tests were performed using the Pearson correlation (R) and 10,000 permutations.

Note that 21 networks (resp. 12 networks) present a negative phylogenetic signal in the species traits of for guild A (resp. guild B). Because these networks are marginal and are likely false positives of the Mantel tests, we discarded them for testing the statistical performances (statistical power and false positive rate) of the different approaches measuring phylogenetic signal in species interactions.

**(B)** We then tested the phylogenetic signal in species' traits using Pagel's λ. For each simulated network, panel (a) (resp. (b)) represents the results of a likelihood ratio test (LRT) comparing the maximum-likelihood Pagel's λ with a model with λ=0. For each combination of parameters ($\alpha_A$ and $\alpha_B$), the bar indicates the percentage of simulated networks that present a significant phylogenetic signal (in green; p-value<0.05) or no significant phylogenetic signal (in yellow; p-value>0.05).

### (A) Testing the phylogenetic signal in species traits using Mantel tests:

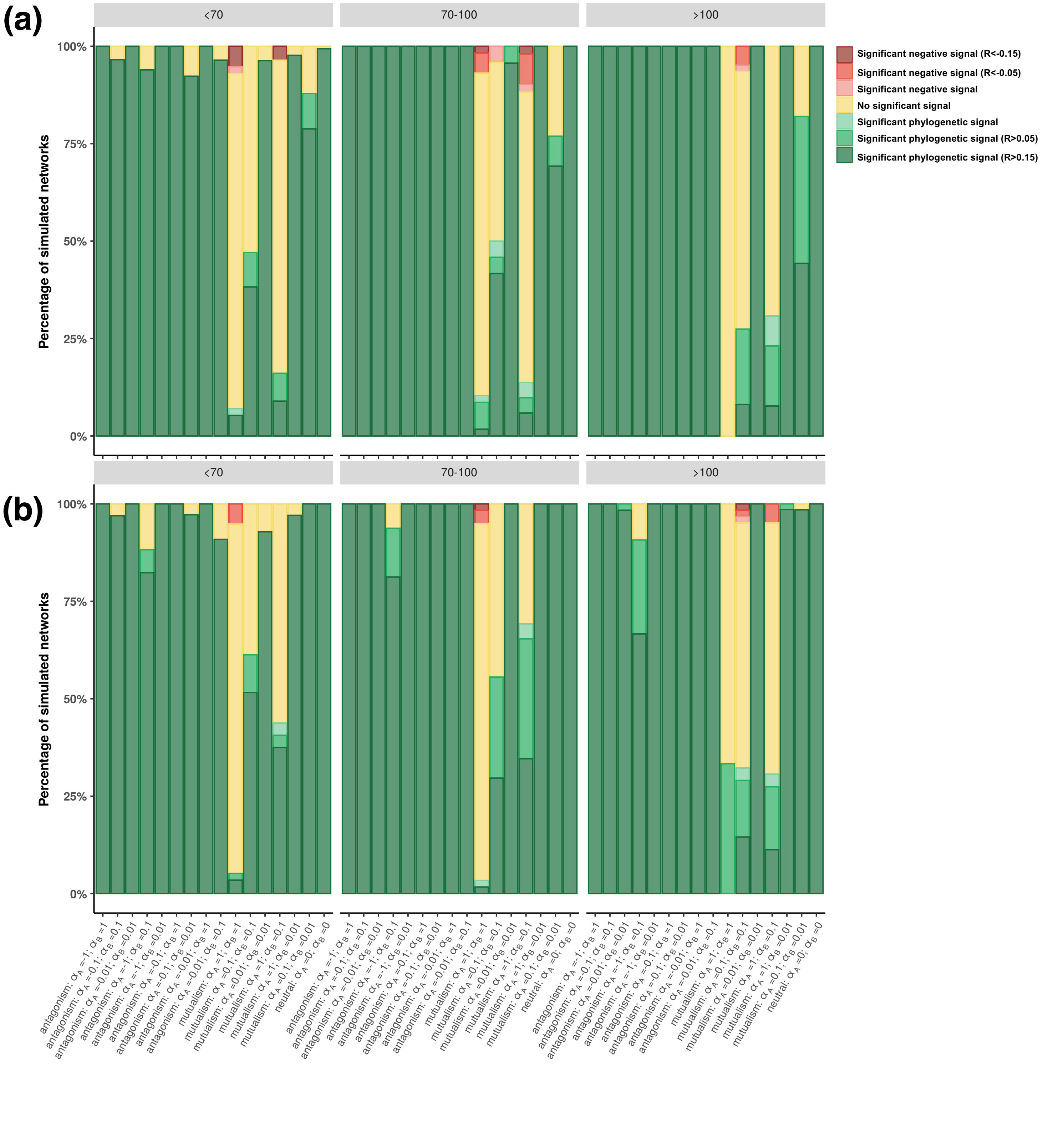

### (B) Testing the phylogenetic signal in species traits using Pagel's λ:

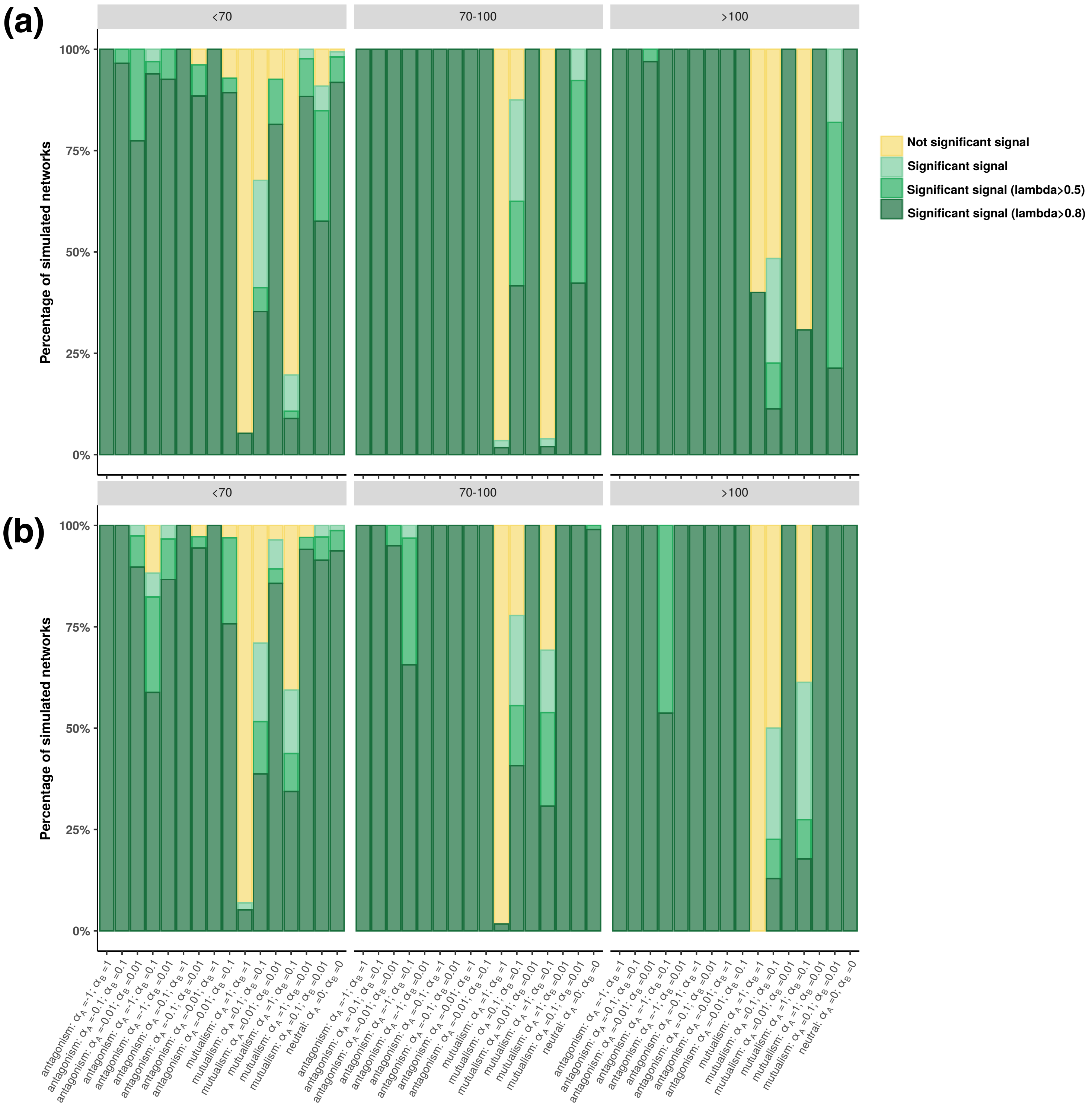

**Supplementary Figure 6: The phylogenetic signals in species interactions estimated using simple Mantel tests in guilds A (left) and B (right) in the *BipartiteEvol* simulations (Maliet *et al.*, 2020). Such phylogenetic signals vary according to the set of parameters ($\alpha_A$ and $\alpha_B$), the different measures of the ecological distances, and the correlations (R) used for the Mantel tests: Pearson (A), Spearman (B), or Kendall (C).**

For each combination of parameters ($\alpha_A$ and $\alpha_B$), the bar indicates the percentage of simulated networks that present a significant positive correlation (in green; p-value<0.05 for the test of phylogenetic signal), a significant negative correlation (in red; p-value<0.05 for the test of negative phylogenetic signal), or no significant correlation (in yellow; both p-values>0.05).

The different panels in rows correspond to the 4 tested ecological distances (Jaccard or UniFrac distances computed with weighted or unweighted networks) and for each panel, the subpanels detail the results according to the network size (total number of species from guilds A and B) separated in three categories (<150, 150-250, or >250 species; see Table S5 for the number of simulated networks per category).

One-tailed Mantel tests were performed using 10,000 permutations.

### (A) Phylogenetic signals in species interactions estimated using simple Mantel tests with Pearson correlations:

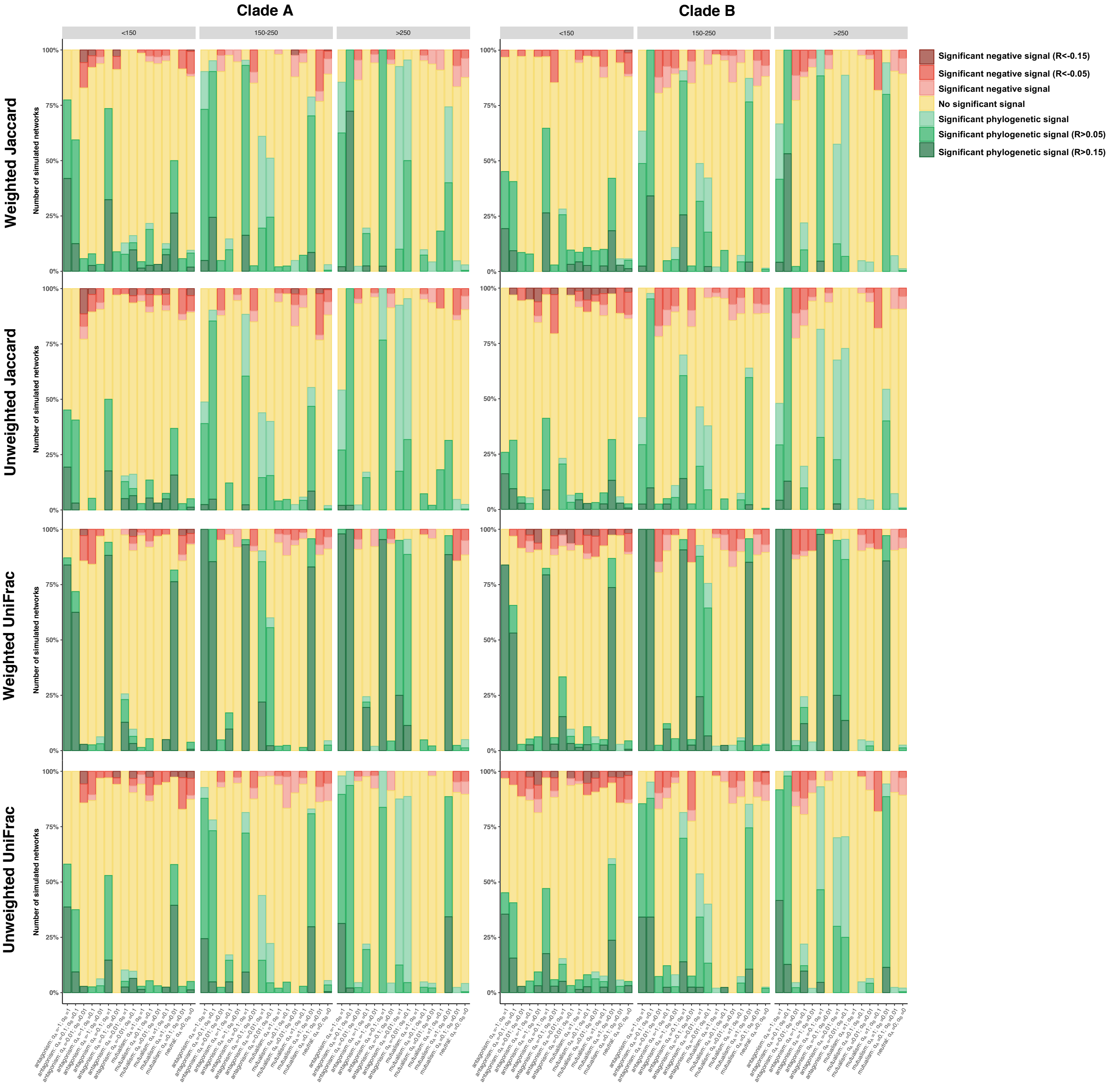

**(B) Phylogenetic signals in species interactions estimated using simple Mantel tests with Spearman correlations:**

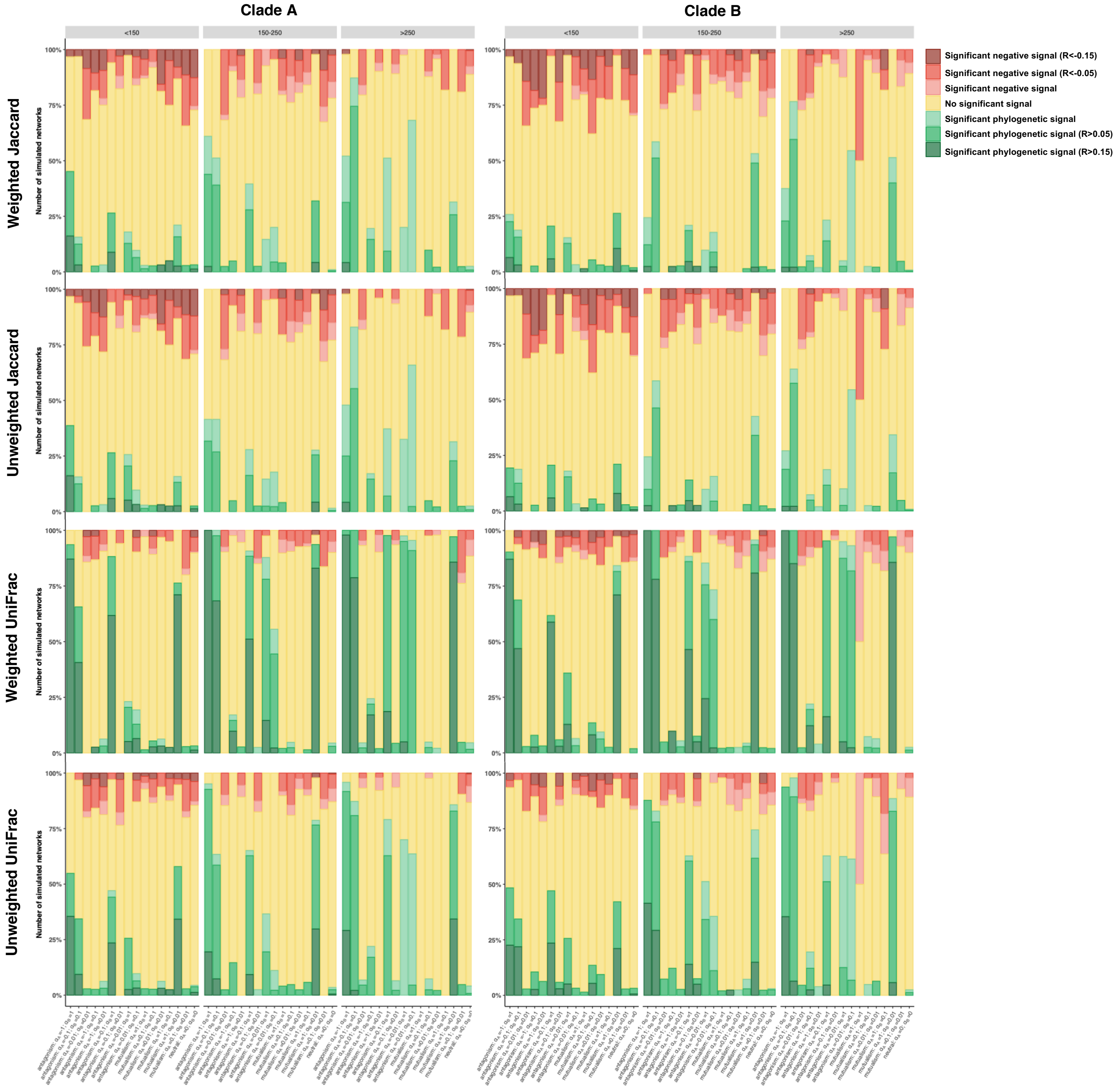

### (C) Phylogenetic signals in species interactions estimated using simple Mantel tests with Kendall correlations:

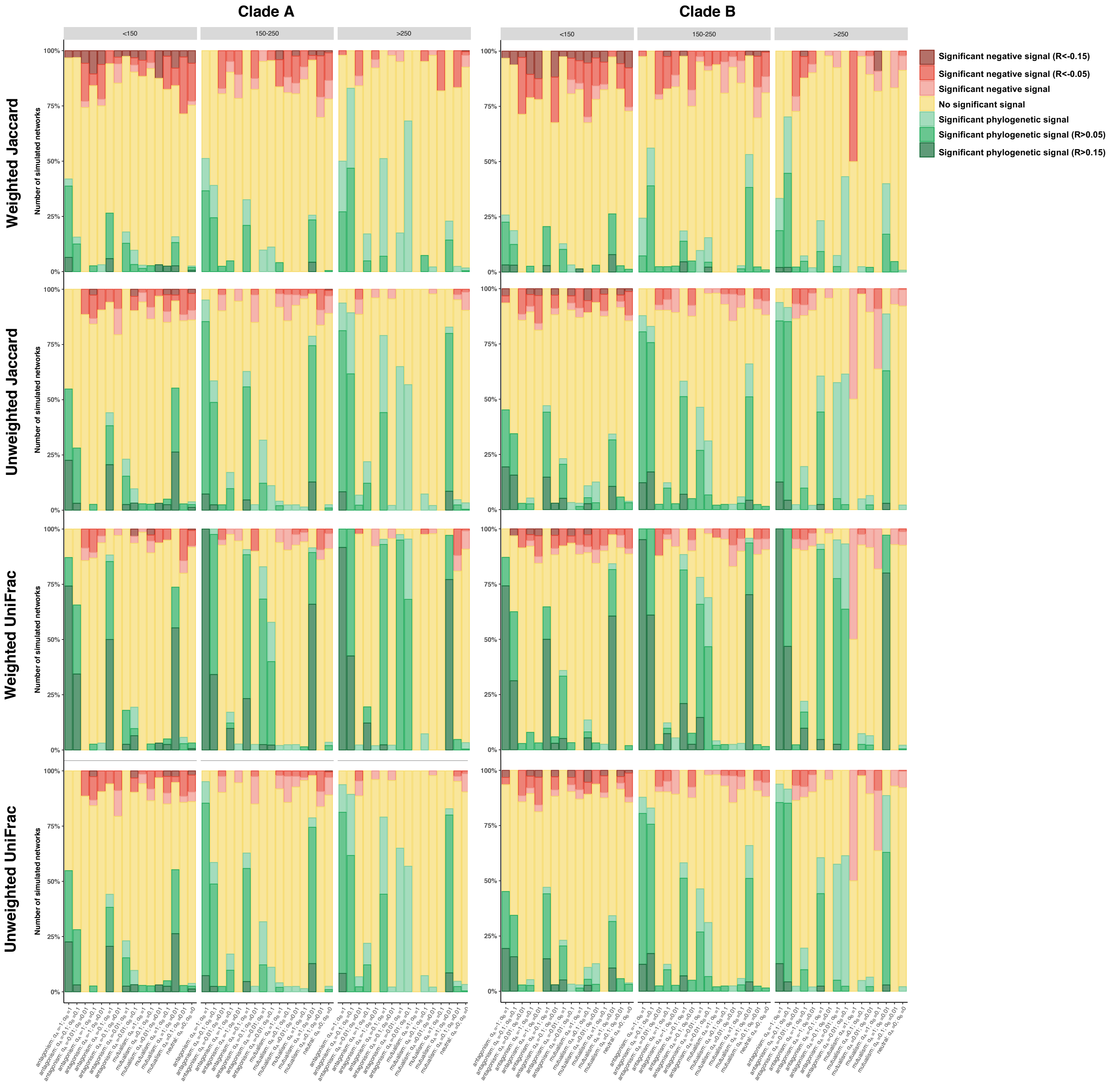

**Supplementary Figure 7: Relationship between the strength of phylogenetic signal in species traits (measured using Mantel correlation (R) (a-c) or Pagel's λ (d-f)) and the strength of phylogenetic signal in species interactions (measured using Mantel correlation (a, b, d, e) and the parameter *d* of PBLM (c, f)).**

Only the results for clade A are represented here (clade B gave similar results).

Each dot corresponds to one simulated network. The shape of the dots indicates whether or not we expected phylogenetic signal in species interactions and the color indicates whether or not we found a significant phylogenetic signal in species interactions.

Phylogenetic signal in species interactions was first tested using Mantel tests with either weighted Jaccard (a, d) or weighted UniFrac (b, e) distances. One-tailed Mantel tests between phylogenetic distances and ecological distances were performed using 10,000 permutations. We reported significant positive correlations (in green; p-value<0.05 for the test of phylogenetic signal), significant negative correlations (in red; p-value<0.05 for the test of negative phylogenetic signal), or no significant correlation (in yellow; both p-values>0.05). Second, phylogenetic signal was estimated using PBLM (c, f). We reported non-significant (in yellow; $MSE \geq MSE_{star}$) or significant (green; $MSE < MSE_{star}$) phylogenetic signals.

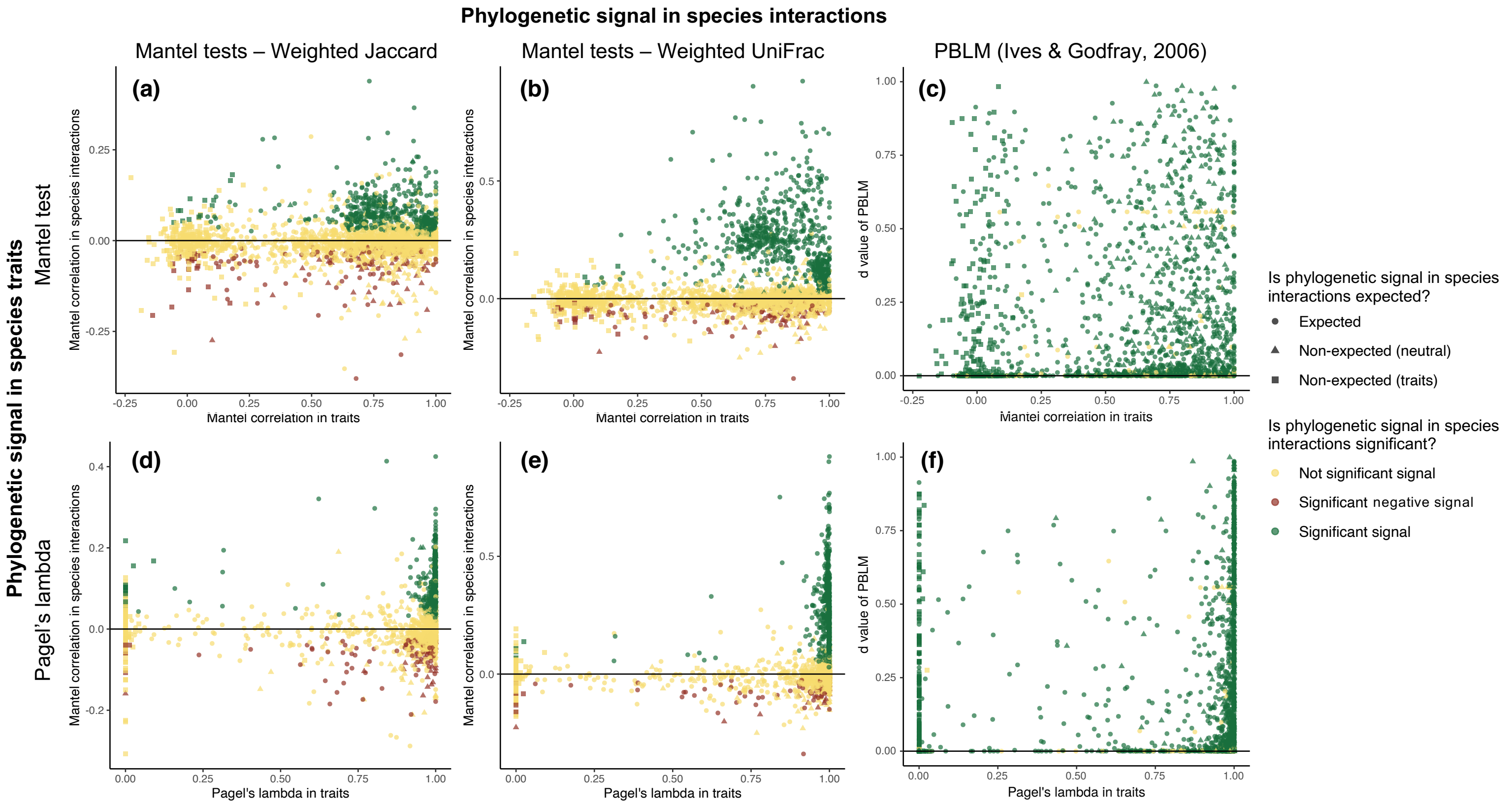
Phylogenetic signal in species interactions
Mantel tests – Weighted Jaccard
Mantel tests – Weighted UniFrac
PBLM (Ives & Godfray, 2006)
Phylogenetic signal in species traits
Mantel test
Pagel's lambda
(a)
(b)
(c)
(d)
(e)
(f)
Mantel correlation in species interactions
d value of PBLM
Mantel correlation in traits
Pagel's lambda in traits
Is phylogenetic signal in species interactions expected?
Expected
Non-expected (neutral)
Non-expected (traits)
Is phylogenetic signal in species interactions significant?
Not significant signal
Significant negative signal
Significant signal

**Supplementary Figure 8: Phylogenetic signals estimated using the Phylogenetic bipartite linear model (PBLM; Ives & Godfray, 2006) in the networks simulated using *BipartiteEvol* (Maliet *et al.*, 2020) according to the different sets of parameters ($\alpha_A$ and $\alpha_B$): When evaluating its significance using mean square errors (MSE), PBLM present a very high false positive rate.**

**(a-b)** For each combination of parameters ($\alpha_A$ and $\alpha_B$), the bar indicates the percentage of simulated networks that present no significant (in yellow; $MSE \geq MSE_{star}$) or a significant (in green; $MSE < MSE_{star}$) phylogenetic signals. Phylogenetic signals are shaded from light green to dark green according to the strength of the signal (e.g. in dark green if $d_A > 0.15$ or $d_B > 0.15$). PBLM was run on the weighted networks (a) or on the unweighted networks (b).

**(c-d) PBLM presents a very high false positive rate even when more stringent cutoffs were used:** For each combination of parameters ($\alpha_A$ and $\alpha_B$), the bar indicates the percentage of simulated networks that present no significant (in yellow; $(MSE_{star} - MSE)/MSE_{star} \leq 5\%$) or a significant (in green; $(MSE_{star} - MSE)/MSE_{star} > 5\%$)) phylogenetic signals. Phylogenetic signals are shaded from light green to dark green according to the strength of the signal (e.g. in dark green if $d_A > 0.15$ or $d_B > 0.15$). PBLM was run on the weighted networks (c) or on the unweighted networks (d).

For each panel, the subpanels detail the results according to the network size (total number of species from guilds A and B) separated in three categories (<150, 150-250, or >250 species). See Table S5 for the number of simulated networks per category (in particular the 100% of strong significant signal in the simulations mutualism (i) and with sizes >250 is only supported by 2 simulated networks).

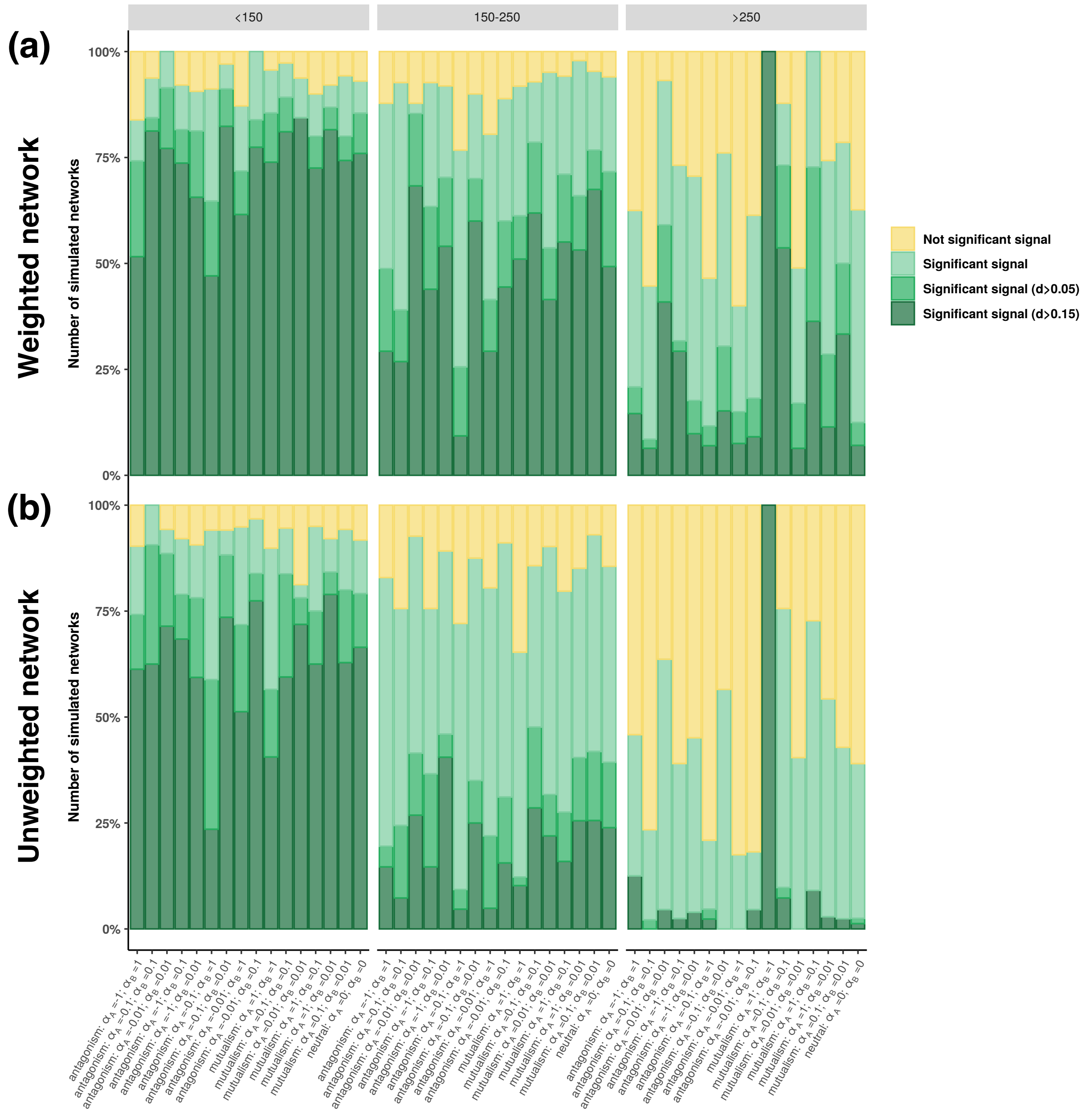

(a)
(b)
Weighted network
Unweighted network
Number of simulated networks
<150
150-250
>250
100%
75%
50%
25%
0%
Not significant signal
Significant signal
Significant signal (d>0.05)
Significant signal (d>0.15)

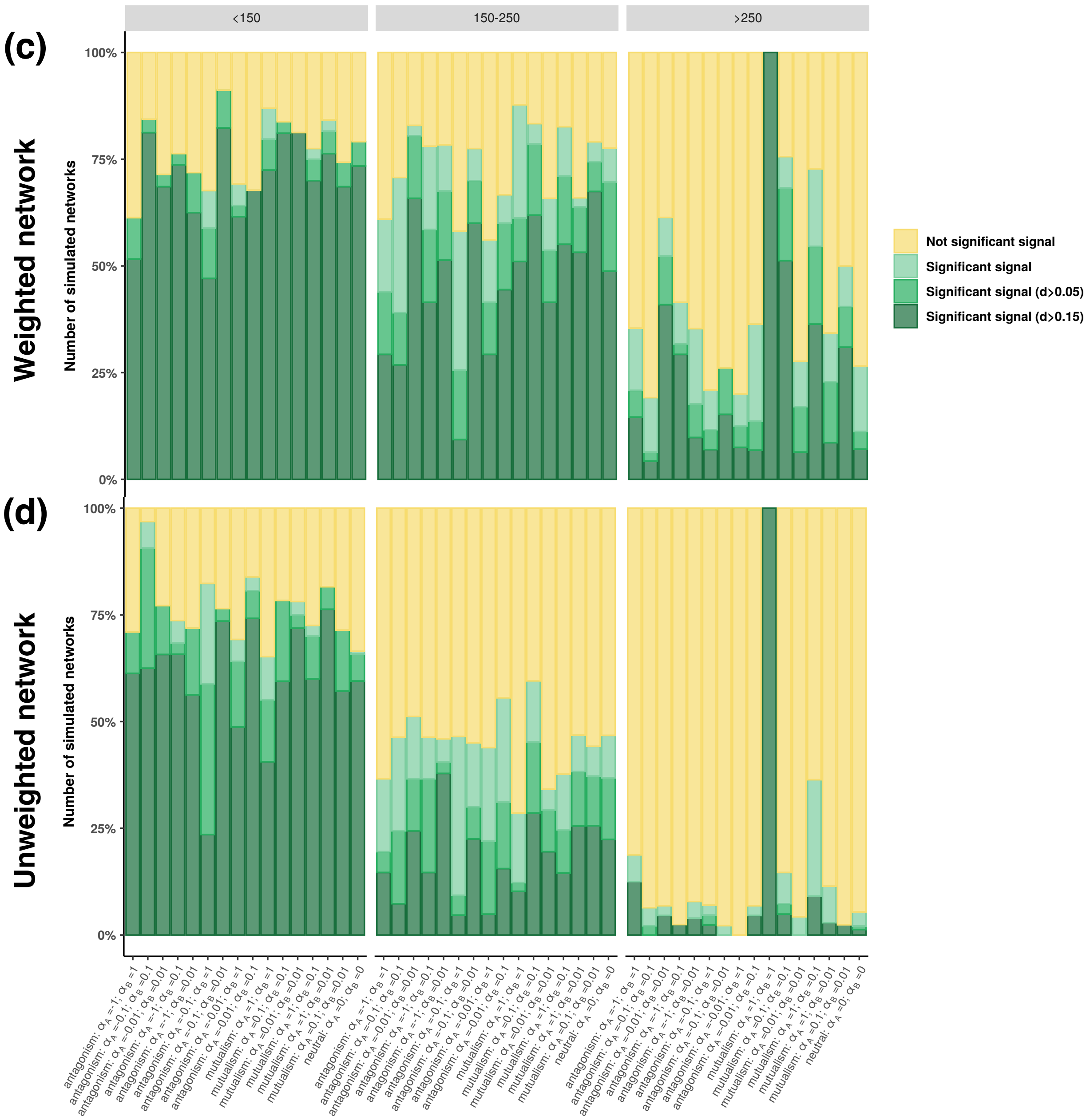

(c)
Weighted network
(d)
Unweighted network
<150
150-250
>250
Number of simulated networks
100%
75%
50%
25%
0%
Not significant signal
Significant signal
Significant signal (d>0.05)
Significant signal (d>0.15)
antagonism: $\alpha_A$ =-1; $\alpha_B$ =1
antagonism: $\alpha_A$ =-0.1; $\alpha_B$ =0.1
antagonism: $\alpha_A$ =-0.01; $\alpha_B$ =0.01
antagonism: $\alpha_A$ =-1; $\alpha_B$ =0.1
antagonism: $\alpha_A$ =-1; $\alpha_B$ =0.01
antagonism: $\alpha_A$ =-0.1; $\alpha_B$ =1
antagonism: $\alpha_A$ =-0.1; $\alpha_B$ =0.01
antagonism: $\alpha_A$ =-0.01; $\alpha_B$ =1
antagonism: $\alpha_A$ =-0.01; $\alpha_B$ =0.1
mutualism: $\alpha_A$ =1; $\alpha_B$ =1
mutualism: $\alpha_A$ =0.1; $\alpha_B$ =0.1
mutualism: $\alpha_A$ =0.01; $\alpha_B$ =0.01
mutualism: $\alpha_A$ =1; $\alpha_B$ =0.1
mutualism: $\alpha_A$ =1; $\alpha_B$ =0.01
mutualism: $\alpha_A$ =0.1; $\alpha_B$ =0.01
neutral: $\alpha_A$ =0; $\alpha_B$ =0

**Supplementary Figure 9: Phylogenetic signal estimated using the Phylogenetic bipartite linear model (PBLM; Ives & Godfray, 2006) in the networks simulated using *BipartiteEvol* (Maliet *et al.*, 2020) according to the different sets of parameters ($\alpha_A$ and $\alpha_B$): When evaluating its significance using bootstrapping, PBLM presents a very high false positive rate even when more stringent cutoffs were used.**

For each PBLM run, its significance was evaluated by performing 100 bootstraps (following Ives & Godfray, 2006). We therefore obtained 95% confidence intervals around both $d_A$ and $d_B$. We considered that we cannot exclude the absence of phylogenetic signal in guild A (resp. B) if values lower than 0.05 belong to the confidence interval of $d_A$ (resp. $d_B$).

For each combination of parameters ($\alpha_A$ and $\alpha_B$), the bar indicates the percentage of simulated networks that present no significant (in yellow; the confidence intervals around $d_A$ and $d_B$ both contain values lower than 0.05) or significant (in green; at least one confidence interval around $d_A$ or $d_B$ does not contain values lower than 0.05) phylogenetic signals: Light green indicates that one confidence interval around $d_A$ or $d_B$ does not contain values lower than 0.05 (*i.e.* there is a significant phylogenetic signal in one guild only), whereas dark green indicates that both confidence intervals around $d_A$ and $d_B$ does not contain values lower than 0.05 (*i.e.* there are significant phylogenetic signals in both guilds).

PBLM was run on the weighted networks (a) or on the unweighted networks (b). Because of computational limitations, bootstrapping was only run for *BipartiteEvol* networks simulated with 500 pairs of individuals only (*i.e.* only 400 simulated networks). In addition, the results of only 387 networks are present here: indeed, the computation for 13 networks were too long and computationally intensive (more than 100 Go of memory during several days on a computer cluster) to be computed.

Note that this bootstrapping strategy is likely incorrect when interaction networks have low connectances (Ives & Godfray, 2006).

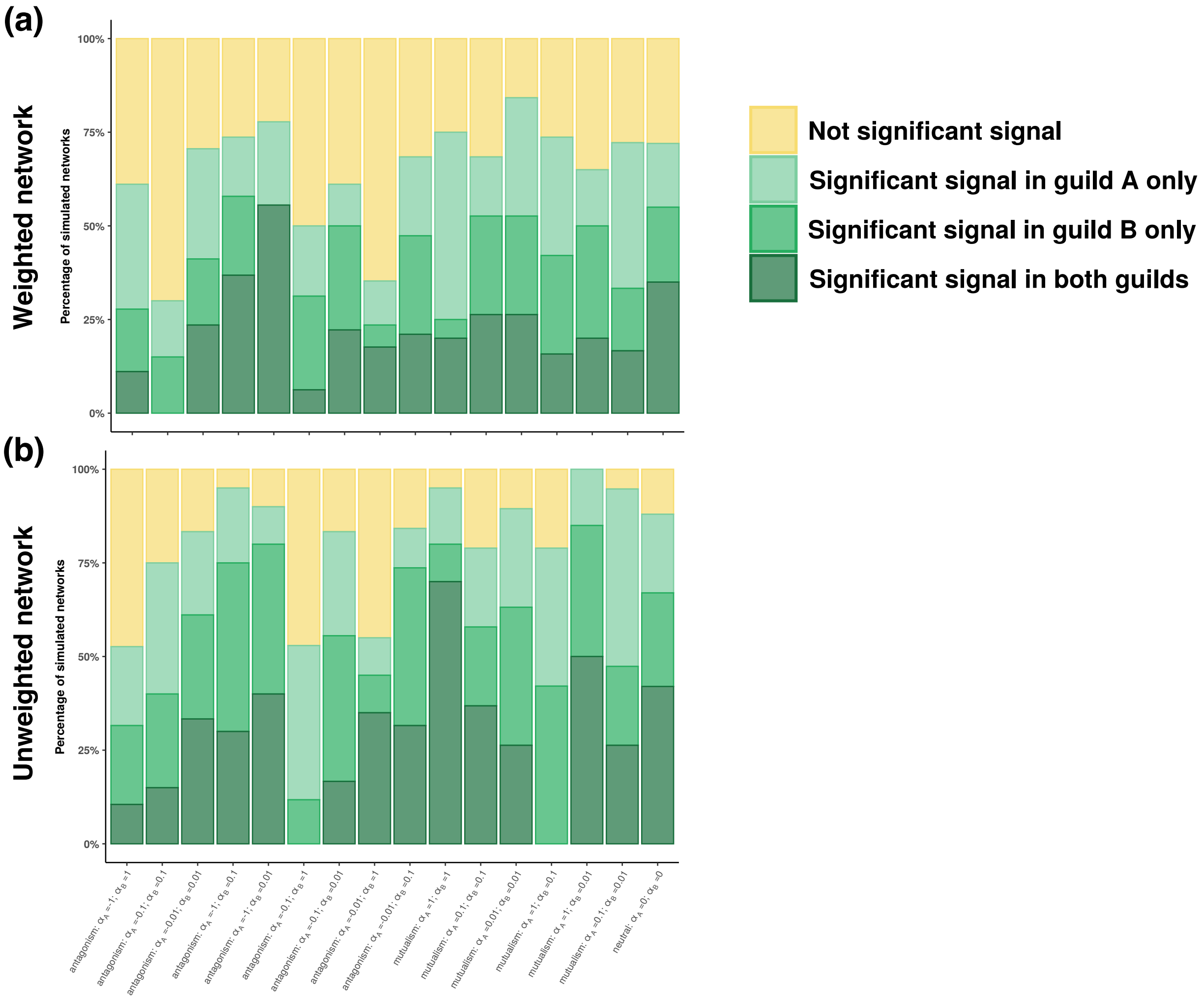
(a)
Weighted network
Percentage of simulated networks
100%
75%
50%
25%
0%
(b)
Unweighted network
Percentage of simulated networks
100%
75%
50%
25%
0%
antagonism: $\alpha_A$ =-1; $\alpha_B$ =1
antagonism: $\alpha_A$ =-0.1; $\alpha_B$ =0.1
antagonism: $\alpha_A$ =-0.01; $\alpha_B$ =0.01
antagonism: $\alpha_A$ =-1; $\alpha_B$ =0.1
antagonism: $\alpha_A$ =-1; $\alpha_B$ =0.01
antagonism: $\alpha_A$ =-0.1; $\alpha_B$ =1
antagonism: $\alpha_A$ =-0.1; $\alpha_B$ =0.01
antagonism: $\alpha_A$ =-0.01; $\alpha_B$ =1
antagonism: $\alpha_A$ =-0.01; $\alpha_B$ =0.1
mutualism: $\alpha_A$ =1; $\alpha_B$ =1
mutualism: $\alpha_A$ =0.1; $\alpha_B$ =0.1
mutualism: $\alpha_A$ =0.01; $\alpha_B$ =0.01
mutualism: $\alpha_A$ =1; $\alpha_B$ =0.1
mutualism: $\alpha_A$ =1; $\alpha_B$ =0.01
mutualism: $\alpha_A$ =0.1; $\alpha_B$ =0.01
neutral: $\alpha_A$ =0; $\alpha_B$ =0
Not significant signal
Significant signal in guild A only
Significant signal in guild B only
Significant signal in both guilds

**Supplementary Figure 10: The phylogenetic generalized linear mixed model (PGLMM; Rafferty and Ives 2013) applied to the weighted networks simulated using *BipartiteEvol* (Maliet *et al.*, 2020) present a high false positive rate and a low statistical power.**

For each panel, the simulations are divided between networks where phylogenetic signal in species interactions is expected (*i.e.* networks (i) simulated with an effect of the traits on individual fitness - antagonistic and mutualistic simulations - and (ii) presenting traits that are phylogenetically conserved according to a Mantel test – see Supplementary Figure 5A) and networks where phylogenetic signal in species interactions is not expected (*i.e.* neutral simulations ($\alpha = 0$) or simulated networks where we observed no phylogenetic signal in the traits). Thus, in each panel, the first bar indicates the statistical power of the test, whereas the second and third bar indicate the false positive rate of the test. PGLMM were run on weighted networks with a Gaussian distribution **(a-b)**, on weighted networks with a Poisson distribution (**c-d**), or on unweighted networks with a binomial distribution (**e-f**).

**a, c, & e:** Phylogenetic signals in the total number of partners estimated using PGLMM. The bar indicates the percentage of simulated networks that present no significant (in yellow; LRT: p-value>0.05) or significant phylogenetic signals in clade(s) A and/or B (green; LRT: p-value<0.05). The significances of the corresponding parameters of PGLMM ($b_{A[i]}$ and $f_{B[i]}$) were evaluated using stepwise model selection based on likelihood ratio tests (Supplementary Methods 3).

**b, d, & f:** Phylogenetic signals in species interactions estimated using PGLMM. For a given combination of parameters, the bar indicates the percentage of simulated networks that present no significant (in yellow; LRT: p-value>0.05) or significant phylogenetic signals in clades A and/or B (green; LRT: p-value<0.05). The significances of the corresponding parameters of PGLMM ($c_i$ and $g_i$) were evaluated using stepwise model selection based on likelihood ratio tests (Supplementary Methods 3).

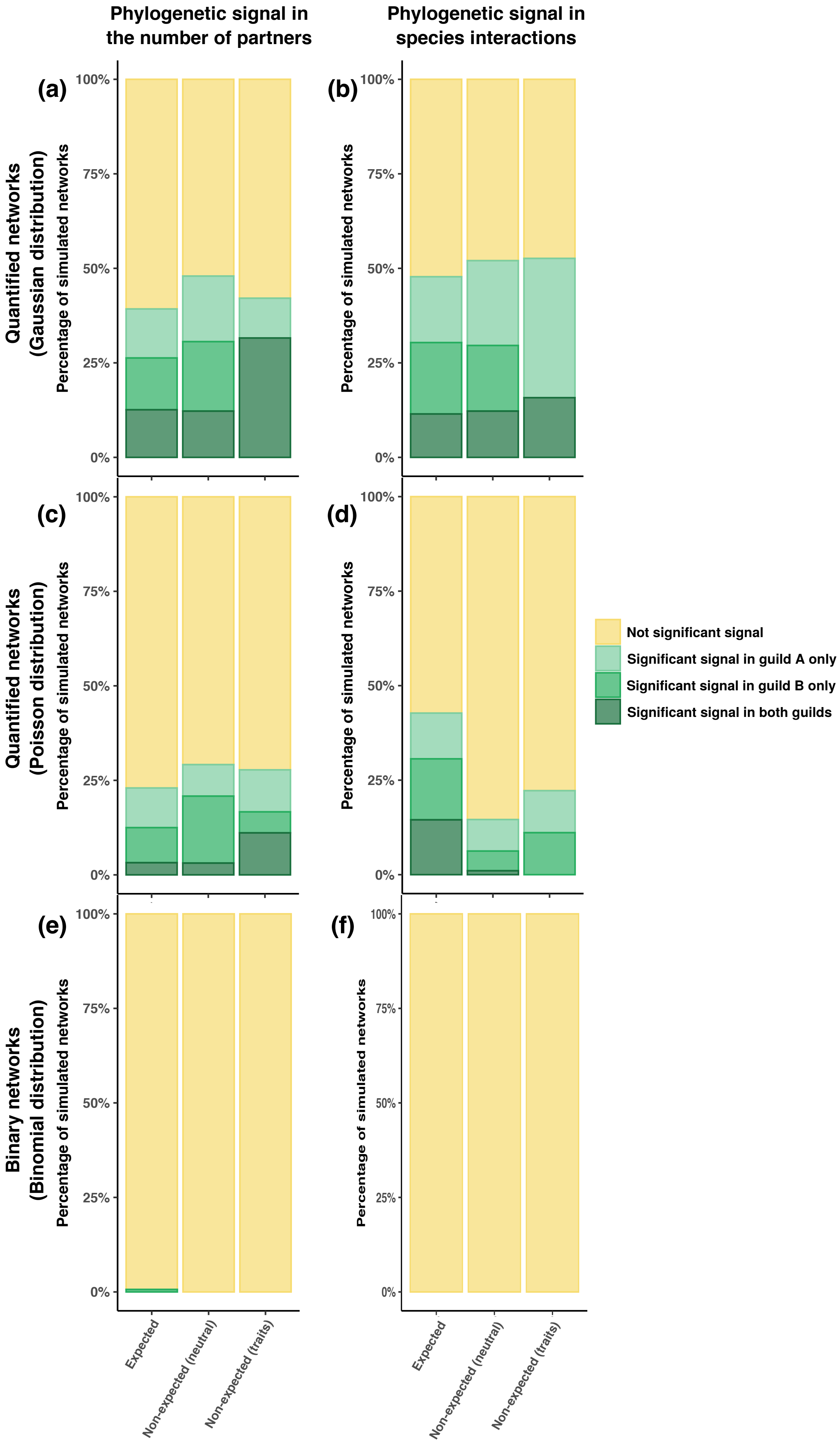

Phylogenetic signal in the number of partners
Phylogenetic signal in species interactions
(a)
(b)
(c)
(d)
(e)
(f)
Quantified networks (Gaussian distribution)
Quantified networks (Poisson distribution)
Binary networks (Binomial distribution)
Percentage of simulated networks
100%
75%
50%
25%
0%
Not significant signal
Significant signal in guild A only
Significant signal in guild B only
Significant signal in both guilds
Expected
Non-expected (neutral)
Non-expected (traits)

**Supplementary Figure 11: The inferred statistical performances of the simple Mantel tests and the Phylogenetic bipartite linear model (PBLM; Ives & Godfray, 2006) are similar when expectations are based on Pagel's λ test of phylogenetic signal in traits**:

For each panel, the simulations are divided between networks where phylogenetic signal in species interactions is expected (*i.e.* networks (i) simulated with an effect of the traits on individual fitness - antagonistic and mutualistic simulations - and (ii) with traits that are phylogenetically conserved according to Pagel's λ test – see Supplementary Figure 5B) and networks where phylogenetic signal in species interactions is not expected (*i.e.* neutral simulations ($\alpha = 0$) or simulations where we observed no phylogenetic signal in the traits).

**a-d:** Phylogenetic signal in species interactions estimated using simple Mantel tests with Pearson correlation (R) in the guilds A (a, c) and B (b, d). The different panels in rows correspond to the 2 tested ecological distances: weighted Jaccard (a, b) or weighted UniFrac (c, d) distances. One-tailed Mantel tests between phylogenetic distances and ecological distances were performed using 10,000 permutations. In each panel, the bars indicate the percentage of simulated networks that present a significant positive correlation (in green; p-value<0.05 for the test of phylogenetic signal), a significant negative correlation (in red; p-value<0.05 for the test of negative phylogenetic signal), or no significant correlation (in yellow; both p-values>0.05). Significant phylogenetic signals (resp. significant negative phylogenetic signals) are shaded from light green (resp. red) to dark green (resp. red) according to the strength of the signal: we arbitrarily considered a "low signal" when R<0.05 (resp. R>-0.05), an "intermediate signal" when 0.05<R<0.15 (resp. -0.05>R>-0.15), and a "strong signal" when R>0.15 (resp. R<-0.15).

**e:** Phylogenetic signals estimated using PBLM. For a given combination of parameters, the bar indicates the percentage of simulated networks that present no significant (in yellow; $MSE \geq MSE_{star}$) or a significant (green; $MSE < MSE_{star}$) phylogenetic signal. Phylogenetic signals are shaded from light green to dark green according to the strength of the signal: we arbitrarily considered a "low signal" when $d_A<0.05$ and $d_B<0.05$, an "intermediate signal" when $d_A>0.05$ or $d_B>0.05$, and a "strong signal" when $d_A>0.15$ or $d_B>0.15$. PBLM was run on the weighted networks.

In each panel, the first bar indicates the statistical power of the test, whereas the second and third bars indicate the false positive rate of the test. Note that the strength of the phylogenetic signals (based on the R and d values) are not directly comparable.

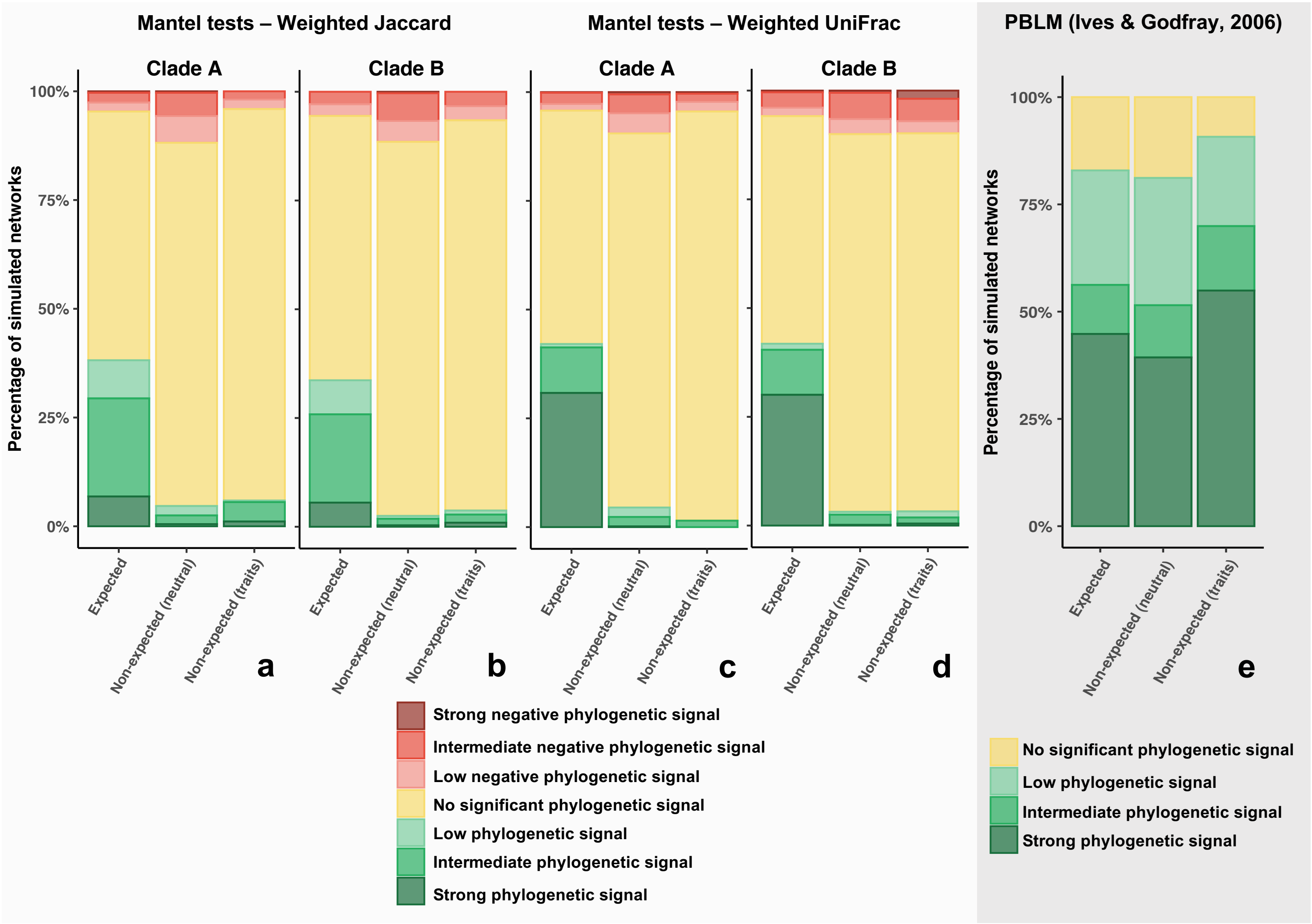

Mantel tests – Weighted Jaccard
Mantel tests – Weighted UniFrac
PBLM (Ives & Godfray, 2006)
Clade A
Clade B
Clade A
Clade B
Percentage of simulated networks
100%
75%
50%
25%
0%
Expected
Non-expected (neutral)
Non-expected (traits)
a
b
c
d
e
Strong negative phylogenetic signal
Intermediate negative phylogenetic signal
Low negative phylogenetic signal
No significant phylogenetic signal
Low phylogenetic signal
Intermediate phylogenetic signal
Strong phylogenetic signal

### Testing the confounding effect of differences in the number of partners on the phylogenetic signal in species interactions:

**Supplementary Figure 12: Phylogenetic signal in the number of partners: correlation between the phylogenetic distances and the differences in numbers of partners evaluated using simple Mantel tests in guilds A (a) and B (b) simulated using *BipartiteEvol* (Maliet *et al.,* 2020).**

For each combination of parameters ($\alpha_A$ and $\alpha_B$), the bar indicates the percentage of simulated networks that present a significant positive correlation (in green; p-value<0.05 for the test of phylogenetic signal), a significant negative correlation (in red; p-value<0.05 for the test of negative phylogenetic signal), or no significant correlation (in yellow; both p-values>0.05).

The subpanels detail the results according to the network size (total number of species from guilds A and B) separated in three categories <150, 150-250, or >250 species; see Table S5).

One-tailed Mantel tests based on the Pearson correlation (R) were performed using 10,000 permutations.

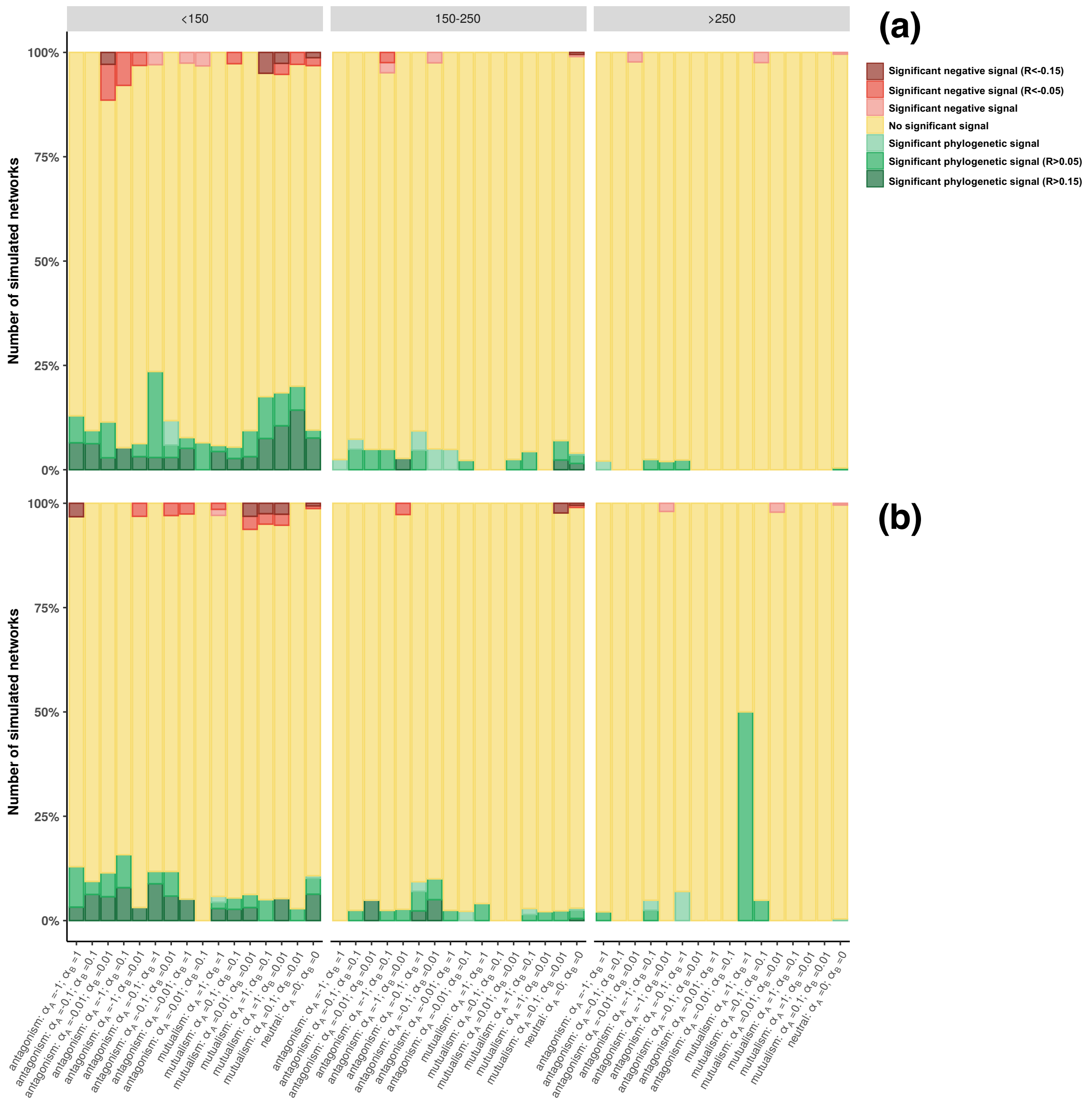

<150
150-250
>250
(a)
(b)
Significant negative signal (R<-0.15)
Significant negative signal (R<-0.05)
Significant negative signal
No significant signal
Significant phylogenetic signal
Significant phylogenetic signal (R>0.05)
Significant phylogenetic signal (R>0.15)
Number of simulated networks
100%
75%
50%
25%
0%
antagonism: αA =-1; αB =1
antagonism: αA =-0.1; αB =0.1
antagonism: αA =-0.01; αB =0.01
antagonism: αA =-1; αB =0.1
antagonism: αA =-1; αB =0.01
antagonism: αA =-0.1; αB =1
antagonism: αA =-0.1; αB =0.01
antagonism: αA =-0.01; αB =1
antagonism: αA =-0.01; αB =0.1
mutualism: αA =1; αB =1
mutualism: αA =0.1; αB =0.1
mutualism: αA =0.01; αB =0.01
mutualism: αA =1; αB =0.1
mutualism: αA =1; αB =0.01
mutualism: αA =0.1; αB =0.01
neutral: αA =0; αB =0

**Supplementary Figure 13: Testing the correlation between ecological distances and differences in numbers of partners in the *BipartiteEvol* simulations (Maliet *et al.*, 2020).**

Panels on the left (resp. on the right) represent the Pearson correlation between the phylogenetic distances and the differences in numbers of partners for guild A (resp. B) evaluated using simple Mantel tests with 10,000 permutations. Each difference in numbers of partners was computed as the pairwise difference in the number of partners (the degree) of two species (*i* and *j*).

The different panels in rows correspond to the 4 tested ecological distances (Jaccard or UniFrac distances computed with weighted or unweighted networks). Each panel details the results per set of parameters ($\alpha_A$ and $\alpha_B$) and colors indicates the total number of pairs of individuals for each guild (from 500 to 5,000).

Boxplots present the median surrounded by the first and third quartiles, and whiskers extend to the extreme values but no further than 1.5 of the inter-quartile range.

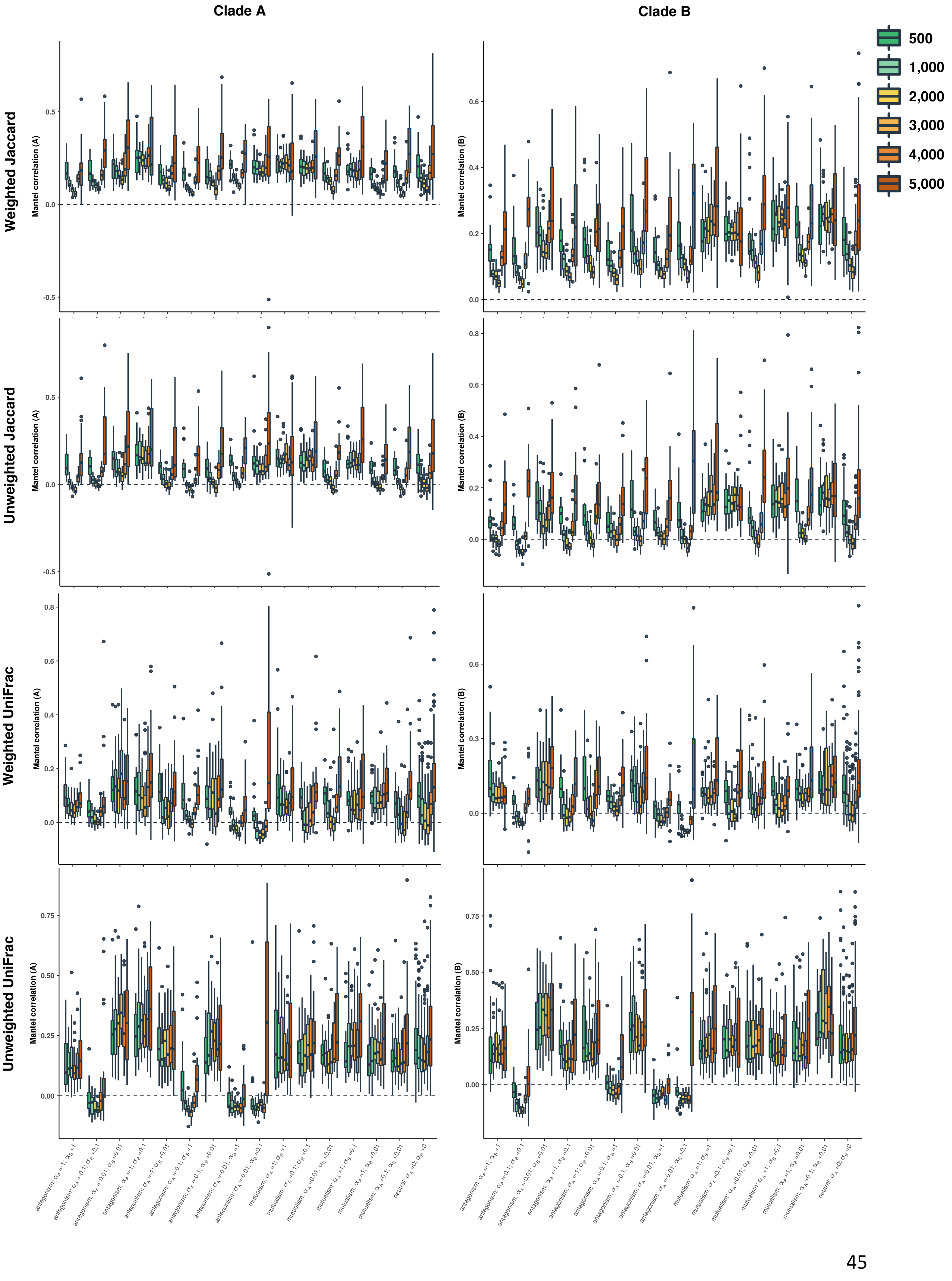

Clade A
Clade B
500
1,000
2,000
3,000
4,000
5,000
Weighted Jaccard
Unweighted Jaccard
Weighted UniFrac
Unweighted UniFrac
Mantel correlation (A)
Mantel correlation (B)
antagonism: $\alpha_A$ =-1; $\alpha_B$ =1
antagonism: $\alpha_A$ =-0.1; $\alpha_B$ =0.1
antagonism: $\alpha_A$ =-0.01; $\alpha_B$ =0.01
antagonism: $\alpha_A$ =-1; $\alpha_B$ =0.1
antagonism: $\alpha_A$ =-1; $\alpha_B$ =0.01
antagonism: $\alpha_A$ =-0.1; $\alpha_B$ =1
antagonism: $\alpha_A$ =-0.1; $\alpha_B$ =0.01
antagonism: $\alpha_A$ =-0.01; $\alpha_B$ =1
antagonism: $\alpha_A$ =-0.01; $\alpha_B$ =0.1
mutualism: $\alpha_A$ =1; $\alpha_B$ =1
mutualism: $\alpha_A$ =0.1; $\alpha_B$ =0.1
mutualism: $\alpha_A$ =0.01; $\alpha_B$ =0.01
mutualism: $\alpha_A$ =1; $\alpha_B$ =0.1
mutualism: $\alpha_A$ =1; $\alpha_B$ =0.01
mutualism: $\alpha_A$ =0.1; $\alpha_B$ =0.01
neutral: $\alpha_A$ =0; $\alpha_B$ =0

**Supplementary Figure 14: Phylogenetic signals in species interactions estimated using partial Mantel tests in guilds A (left) and B (right) in networks simulated using *BipartiteEvol* (Maliet *et al.,* 2020): partial Mantel tests measured the correlation between the phylogenetic distances and the ecological distances, while controlling for differences in numbers of partners.**

For each combination of parameters ($\alpha_A$ and $\alpha_B$), the bar indicates the percentage of simulated networks that present a significant positive correlation (in green; p-value<0.05 for the test of phylogenetic signal), a significant negative correlation (in red; p-value<0.05 for the test of negative phylogenetic signal), or no significant correlation (in yellow; both p-values>0.05).

The different panels in rows correspond to the 4 tested ecological distances (Jaccard or UniFrac distances computed with weighted or unweighted networks) and for each panel, the subpanels detail the results according to the network size (total number of species from guilds A and B) separated in three categories <150, 150-250, or >250 species; see Table S5).

One-tailed Mantel tests based on the Pearson correlation (R) were performed using 10,000 permutations.

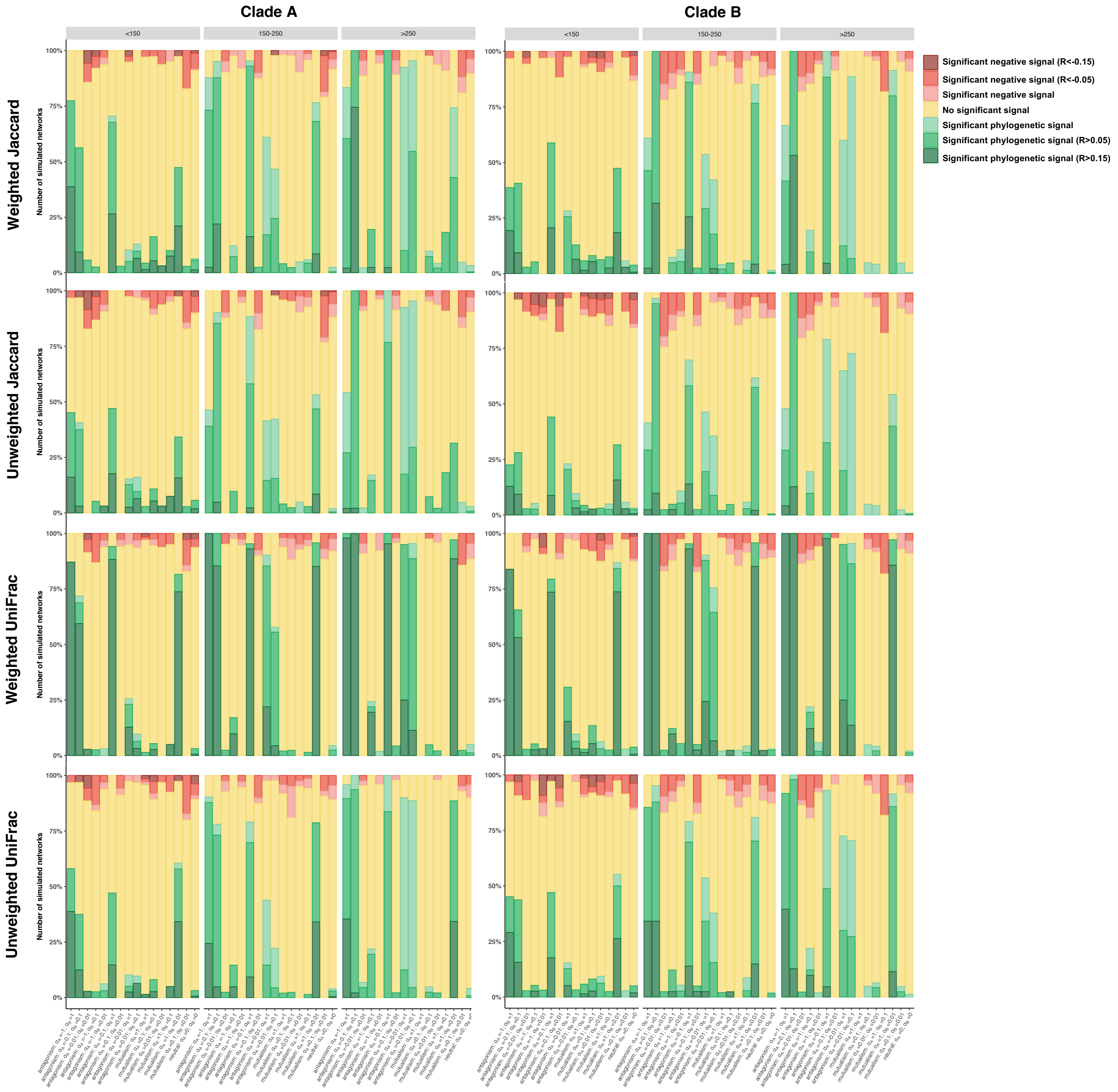

Clade A
Clade B
<150
150-250
>250
Weighted Jaccard
Unweighted Jaccard
Weighted UniFrac
Unweighted UniFrac
Number of simulated networks
100%
75%
50%
25%
0%
Significant negative signal (R<-0.15)
Significant negative signal (R<-0.05)
Significant negative signal
No significant signal
Significant phylogenetic signal
Significant phylogenetic signal (R>0.05)
Significant phylogenetic signal (R>0.15)

**Supplementary Figure 15: Phylogenetic signals in species interactions estimated using simple Mantel tests with beta diversity partitioning in guilds A (left) and B (right) in networks simulated using *BipartiteEvol* (Maliet *et al.*, 2020): beta diversity partitioning decomposes the part of the unweighted Jaccard distances that is due to partner turnover (*i.e.* partner species identity), irrespectively of the number of partners (Baselga, 2010).**

For each combination of parameters ($\alpha_A$ and $\alpha_B$), the bar indicates the percentage of simulated networks that present a significant positive correlation (in green; p-value<0.05 for the test of phylogenetic signal), a significant negative correlation (in red; p-value<0.05 for the test of negative phylogenetic signal), or no significant correlation (in yellow; both p-values>0.05).

For each panel, the subpanels detail the results according to the network size (total number of species from guilds A and B) separated in three categories <150, 150-250, or >250 species; see Table S5).

One-tailed Mantel tests based on the Pearson correlation (R) were performed using 10,000 permutations.

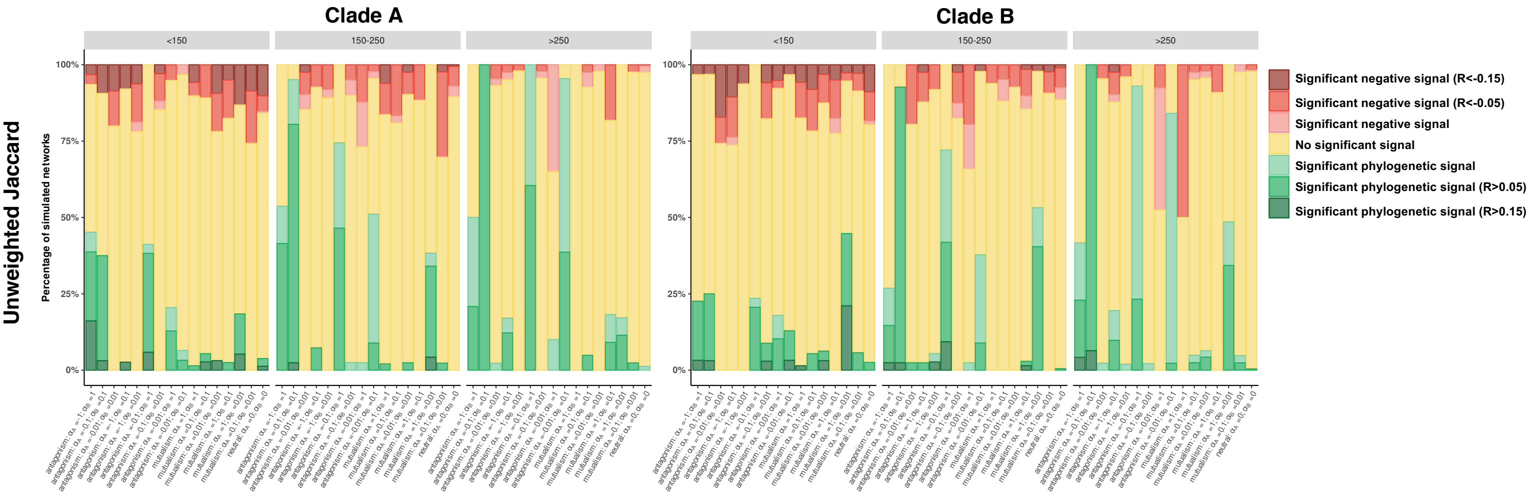

**Supplementary Figure 16: Phylogenetic signals in species interactions estimated <u>using Mantel tests using permutations keeping constant the number of partners per species</u> in guilds A (left) and B (right) in networks simulated using *BipartiteEvol* (Maliet *et al.*, 2020):**

For each combination of parameters ($\alpha_A$ and $\alpha_B$), the bar indicates the percentage of simulated networks that present a significant positive correlation (in green; p-value<0.05 for the test of phylogenetic signal), a significant negative correlation (in red; p-value<0.05 for the test of negative phylogenetic signal), or no significant correlation (in yellow; both p-values>0.05).

The different panels in rows correspond to the 4 tested ecological distances (Jaccard or UniFrac distances computed with weighted or unweighted networks) and for each panel, the subpanels detail the results according to the network size (total number of species from guilds A and B) separated in three categories <150, 150-250, or >250 species; see Table S5).

One-tailed Mantel tests based on the Pearson correlation (R) were performed using 1,000 permutations keeping constant the number of partners per species in guilds A.

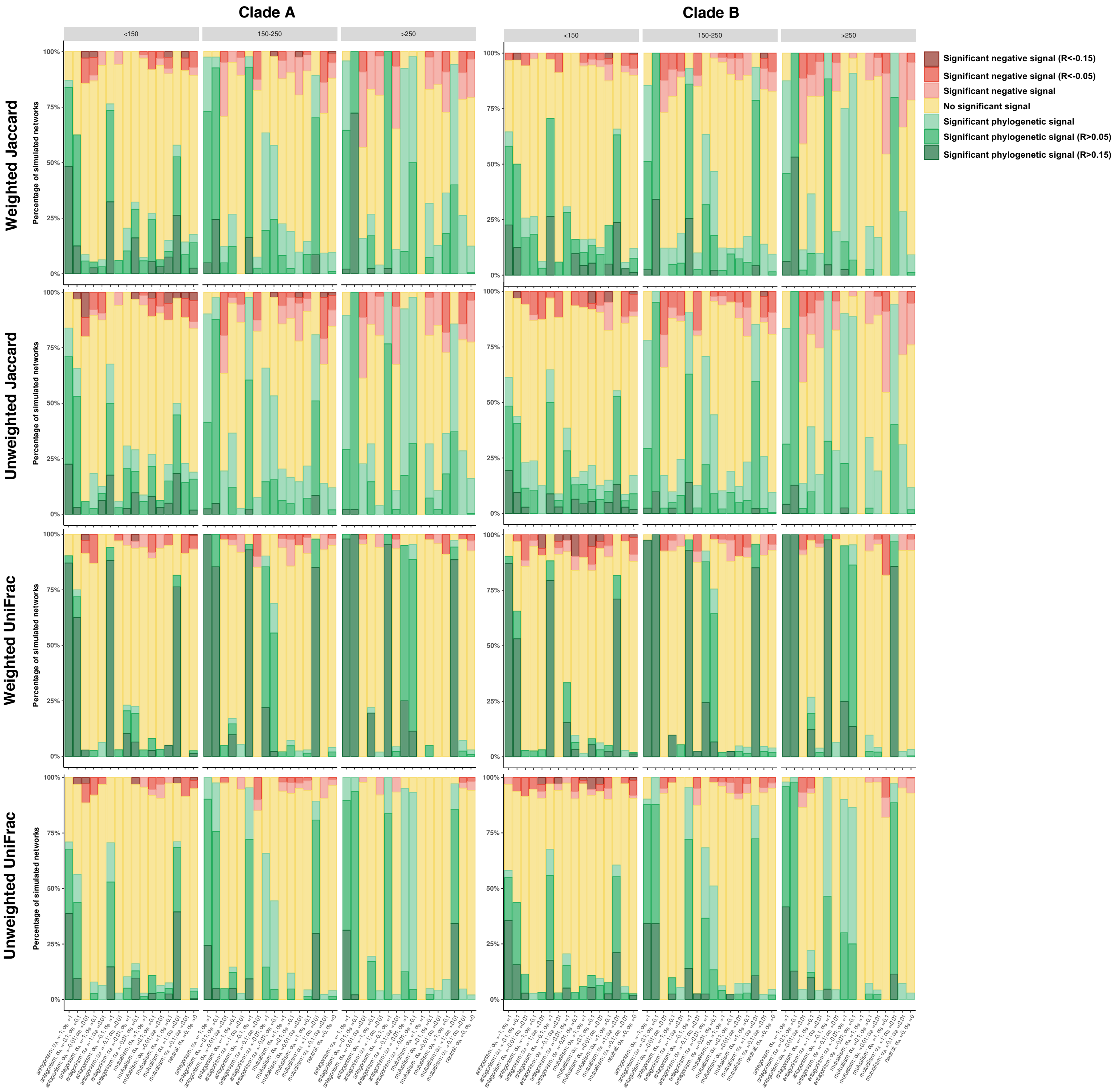
Clade A
Clade B
<150
150-250
>250
Weighted Jaccard
Unweighted Jaccard
Weighted UniFrac
Unweighted UniFrac
Percentage of simulated networks
100%
75%
50%
25%
0%
Significant negative signal (R<-0.15)
Significant negative signal (R<-0.05)
Significant negative signal
No significant signal
Significant phylogenetic signal
Significant phylogenetic signal (R>0.05)
Significant phylogenetic signal (R>0.15)

**Supplementary Figure 17: Realistic size ranges and connectances (*i.e.* ratios of realized interactions) of the networks simulated with phylogenetic conservatism in the number, but not the identity, of partners.**

Numbers of species in guilds A (a) and B (b) and connectances (c) are represented as a function of the $a_A$ values.

$a_A$>0 corresponds to simulations where the number of partners of the species in guild A has evolved following an Ornstein-Uhlenbeck process.

$a_A$=0 corresponds to simulations where the number of partners of the species in guild A has evolved following a Brownian motion.

Boxplots present the median surrounded by the first and third quartiles, and whiskers extend to the extreme values but no further than 1.5 of the inter-quartile range.

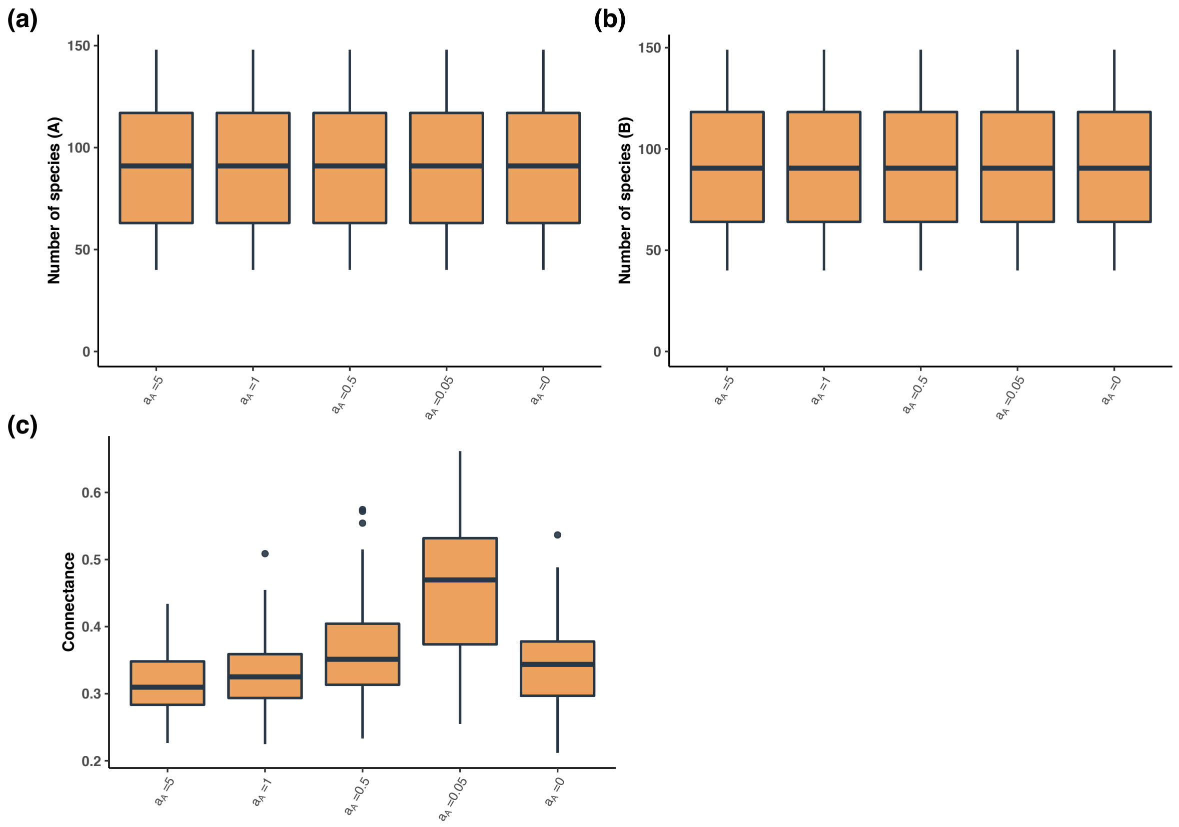

**Supplementary Figure 18: Phylogenetic signal in the number of partners: Correlation between the phylogenetic distances and the differences in numbers of partners evaluated using simple Mantel tests in guild A in networks simulated with phylogenetic conservatism in the number, but not the identity, of partners.**

For each $\alpha_A$ value, the bar indicates the percentage of simulated networks that present a significant positive correlation (in green; p-value<0.05 for the test of phylogenetic signal), a significant negative correlation (in red; p-value<0.05 for the test of negative phylogenetic signal), or no significant correlation (in yellow; both p-values>0.05).

$\alpha_A$>0 corresponds to simulations where the number of partners of the species in guild A has evolved following an Ornstein-Uhlenbeck process.

$\alpha_A$=0 corresponds to simulations where the number of partners of the species in guild A has evolved following a Brownian motion.

The subpanels detail the results according to the network size (total number of species from guilds A and B) separated in three categories (<150, 150-200, or >200 species).

One-tailed Mantel tests based on the Pearson correlation (R) were performed using 10,000 permutations.

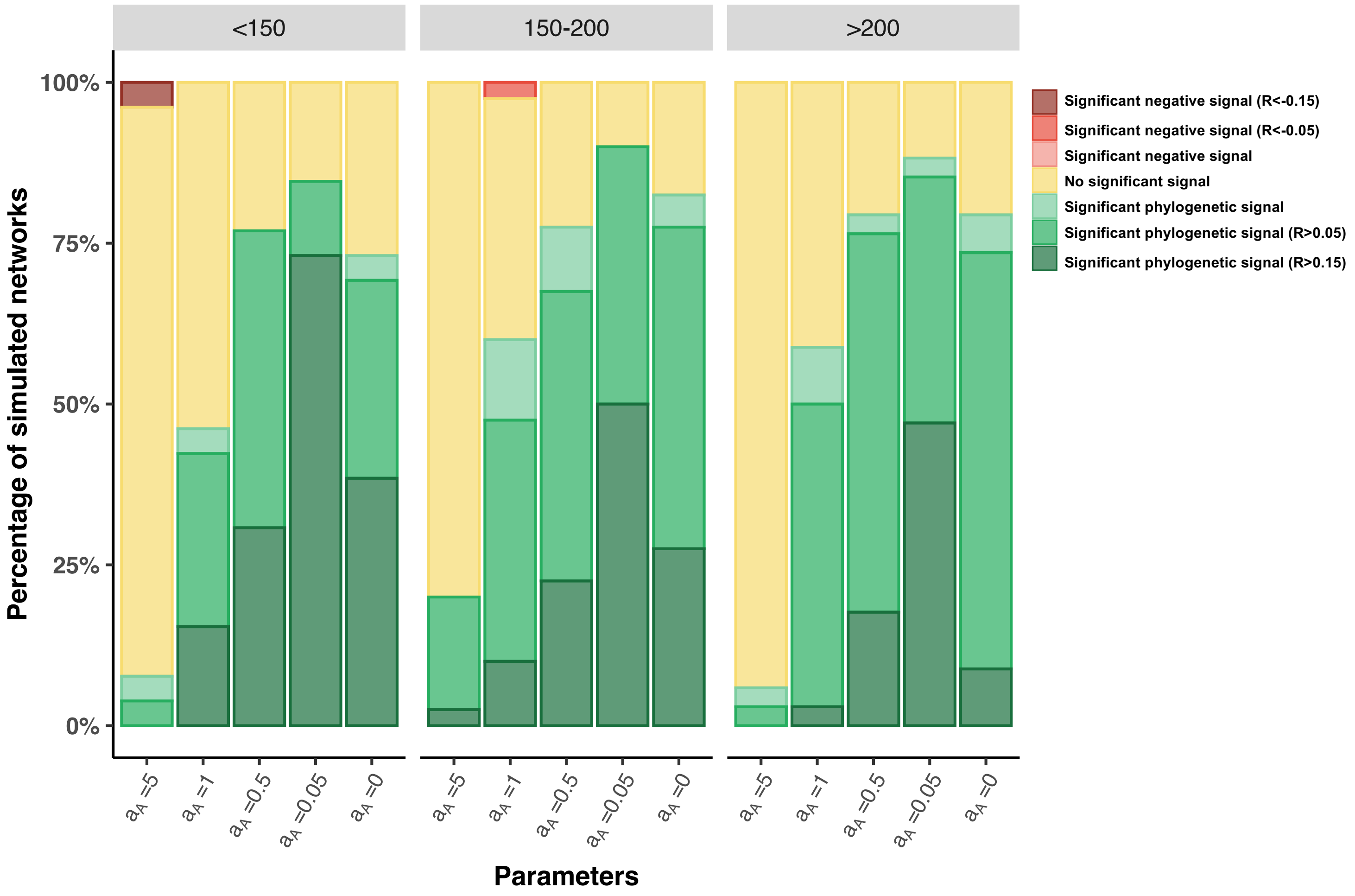

<150
150-200
>200
100%
75%
50%
25%
0%
Percentage of simulated networks
$a_A$ =5
$a_A$ =1
$a_A$ =0.5
$a_A$ =0.05
$a_A$ =0
Parameters
Significant negative signal (R<-0.15)
Significant negative signal (R<-0.05)
Significant negative signal
No significant signal
Significant phylogenetic signal
Significant phylogenetic signal (R>0.05)
Significant phylogenetic signal (R>0.15)

**Supplementary Figure 19: Testing the correlation between ecological distances and differences in numbers of partners in networks simulated with phylogenetic conservatism in the number, but not the identity, of partners.**

Panels represent the Pearson correlation between the phylogenetic distances and the differences in numbers of partners for guild A evaluated using simple Mantel tests with 10,000 permutations. Each difference in numbers of partners was computed as the pairwise difference in the number of partners (the degree) of two species (*i* and *j*). The different panels in (a and b) correspond to the 2 tested ecological distances (unweighted Jaccard or UniFrac distances respectively). Each panel details the results per parameter value ($a_A$) and colors indicates the number of species from guild A. Boxplots present the median surrounded by the first and third quartiles, and whiskers extend to the extreme values but no further than 1.5 of the inter-quartile range.

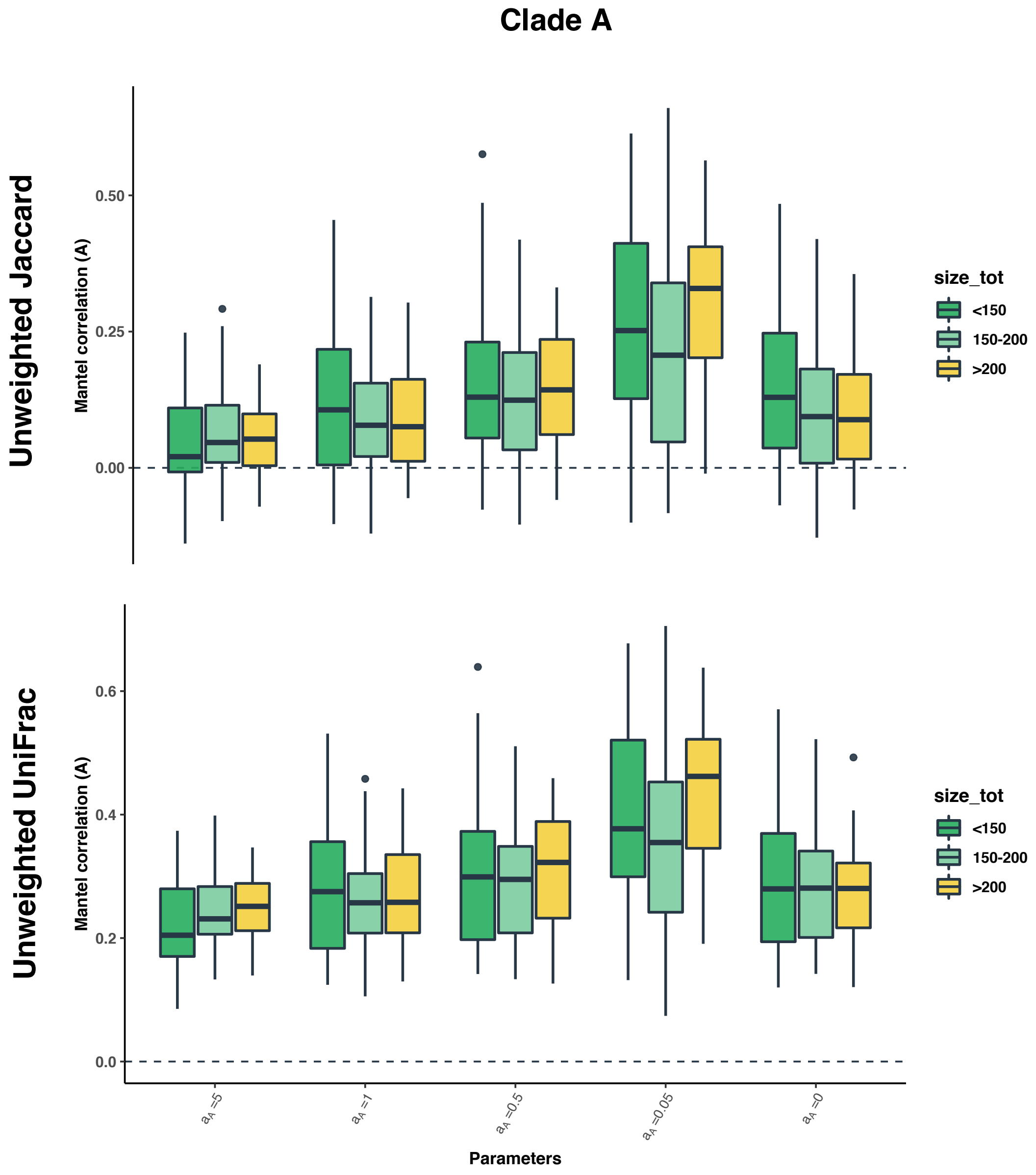

**Supplementary Figure 20: Phylogenetic signals in species interactions estimated using simple Mantel tests in guild A in networks simulated with phylogenetic conservatism in the number, but not the identity, of partners.**

For each $a_A$ value, the bar indicates the percentage of simulated networks that present a significant positive correlation (in green; p-value<0.05 for the test of phylogenetic signal), a significant negative correlation (in red; p-value<0.05 for the test of negative phylogenetic signal), or no significant correlation (in yellow; both p-values>0.05).

$a_A$>0 corresponds to simulations where the number of partners of the species in guild A has evolved following an Ornstein-Uhlenbeck process.

$a_A$=0 corresponds to simulations where the number of partners of the species in guild A has evolved following a Brownian motion.

The different panels in rows correspond to the 2 tested ecological distances (Jaccard or UniFrac distances computed on unweighted networks) and for each panel, the subpanels detail the results according to the network size (total number of species from guilds A and B) separated in three categories (<150, 150-200, or >200 species).

One-tailed Mantel tests based on the Pearson correlation (R) were performed using 10,000 permutations.

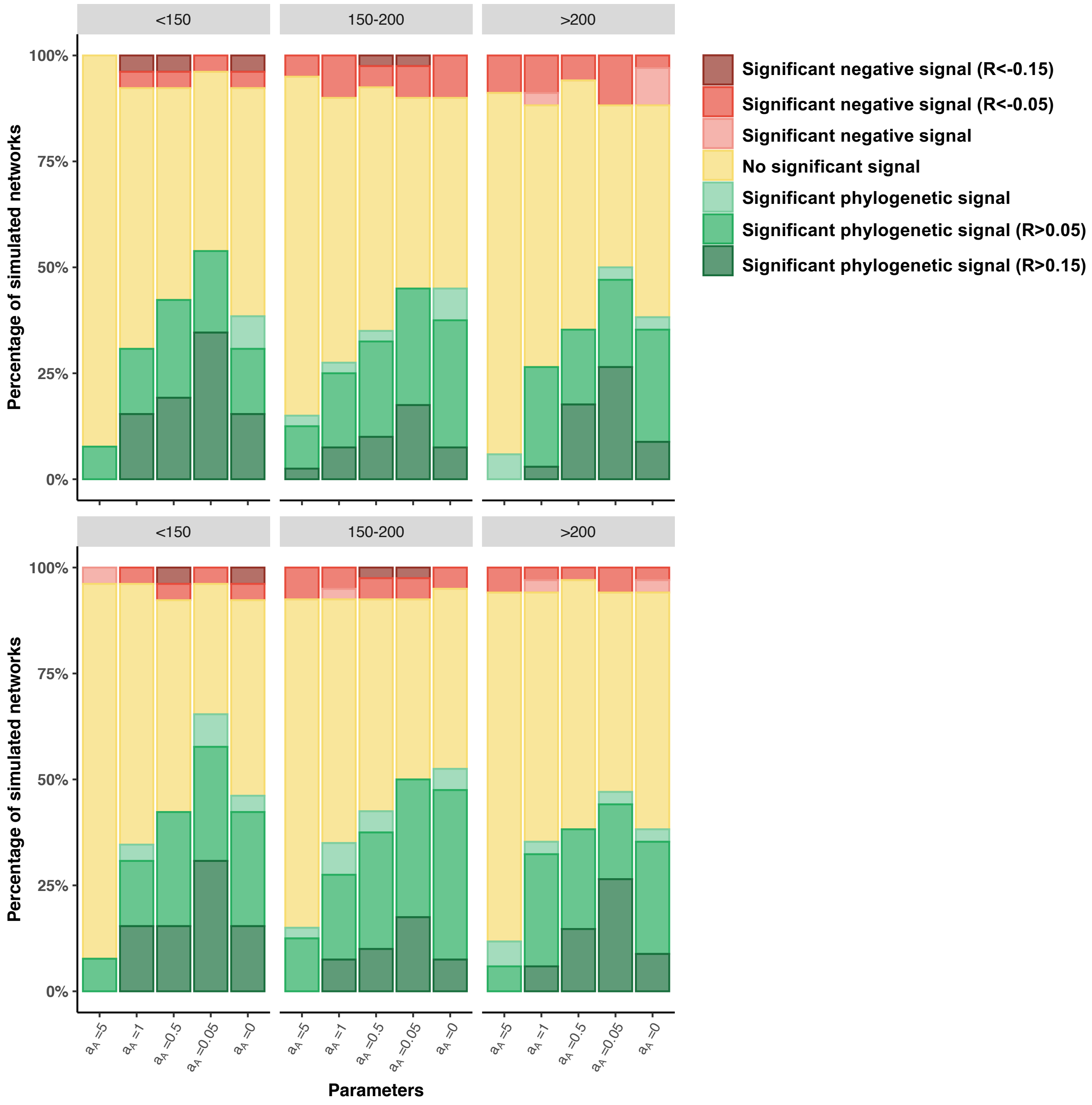

<150
150-200
>200
100%
75%
50%
25%
0%
Percentage of simulated networks
Significant negative signal (R<-0.15)
Significant negative signal (R<-0.05)
Significant negative signal
No significant signal
Significant phylogenetic signal
Significant phylogenetic signal (R>0.05)
Significant phylogenetic signal (R>0.15)
Parameters

**Supplementary Figure 21: Phylogenetic signals in species interactions estimated using <u>partial Mantel tests</u> in guild A in networks simulated with phylogenetic conservatism in the number, but not the identity, of partners: Partial Mantel tests measured the correlation between the phylogenetic distances and the ecological distances, while controlling for the differences in numbers of partners.**

For each $a_A$ value, the bar indicates the percentage of simulated networks that present a significant positive correlation (in green; p-value<0.05 for the test of phylogenetic signal), a significant negative correlation (in red; p-value<0.05 for the test of negative phylogenetic signal), or no significant correlation (in yellow; both p-values>0.05).

$a_A$>0 corresponds to simulations where the number of partners of the species in guild A has evolved following an Ornstein-Uhlenbeck process.

$a_A$=0 corresponds to simulations where the number of partners of the species in guild A has evolved following a Brownian motion.

The different panels in rows correspond to the 2 tested ecological distances (Jaccard or UniFrac distances computed on unweighted networks) and for each panel, the subpanels detail the results according to the network size (total number of species from guilds A and B) separated in three categories (<150, 150-200, or >200 species).

One-tailed Mantel tests based on the Pearson correlation (R) were performed using 10,000 permutations.

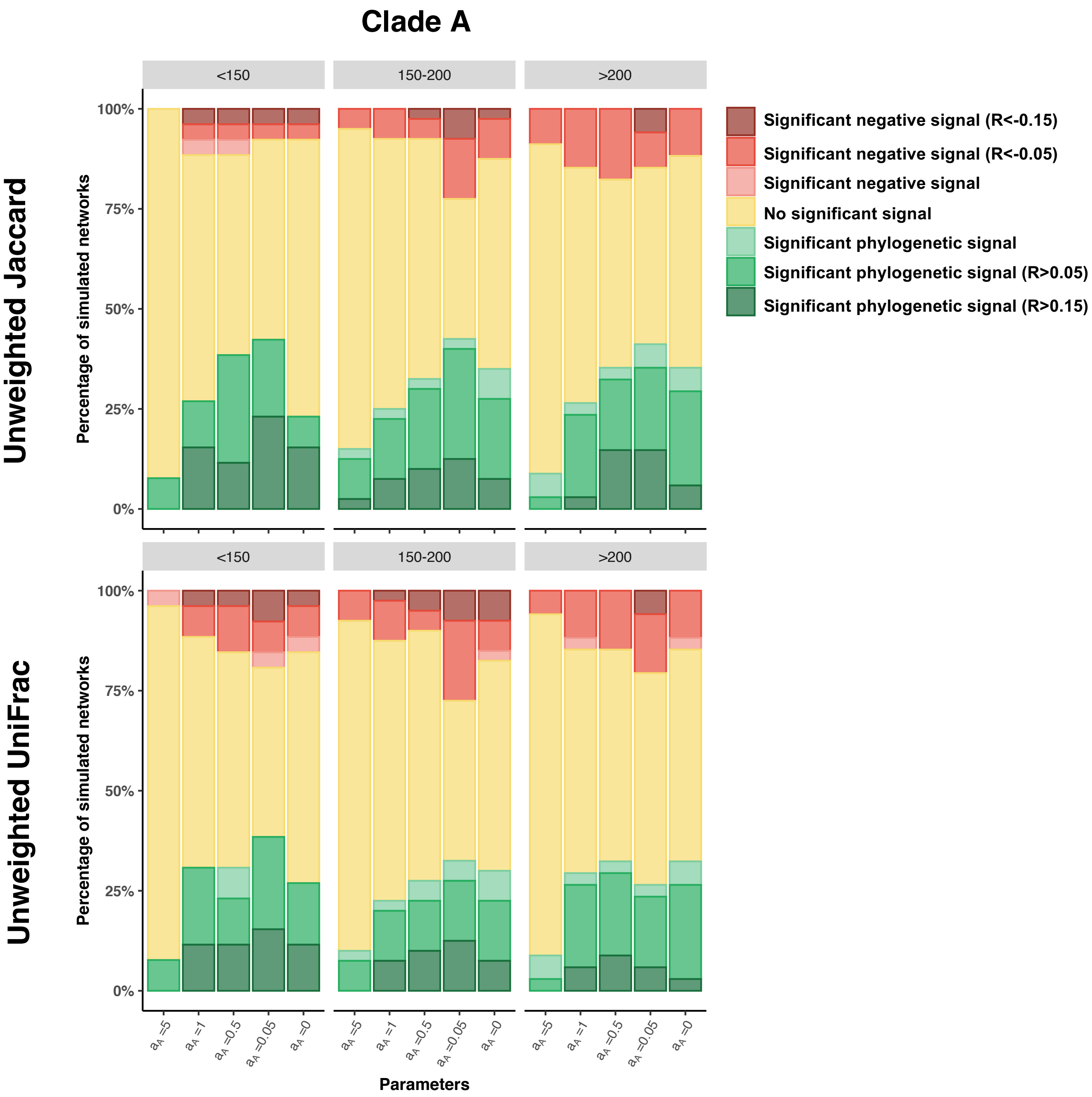
Clade A
Unweighted Jaccard
Unweighted UniFrac
Percentage of simulated networks
<150
150-200
>200
100%
75%
50%
25%
0%
Significant negative signal (R<-0.15)
Significant negative signal (R<-0.05)
Significant negative signal
No significant signal
Significant phylogenetic signal
Significant phylogenetic signal (R>0.05)
Significant phylogenetic signal (R>0.15)
$a_A$ =5
$a_A$ =1
$a_A$ =0.5
$a_A$ =0.05
$a_A$ =0
Parameters

**Supplementary Figure 22: Phylogenetic signals in species interactions estimated using simple Mantel tests with beta diversity partitioning in guilds A in networks simulated with phylogenetic conservatism in the number, but not the identity, of partners: beta diversity partitioning decomposes the part of the unweighted Jaccard distances that is due to partner turnover (*i.e.* partner species identity), irrespectively of the number of partners (Baselga, 2010).**

For each $a_A$ value, the bar indicates the percentage of simulated networks that present a significant positive correlation (in green; p-value<0.05 for the test of phylogenetic signal), a significant negative correlation (in red; p-value<0.05 for the test of negative phylogenetic signal), or no significant correlation (in yellow; both p-values>0.05).

$a_A>0$ corresponds to simulations where the number of partners of the species in guild A has evolved following an Ornstein-Uhlenbeck process.

$a_A=0$ corresponds to simulations where the number of partners of the species in guild A has evolved following a Brownian motion.

The subpanels detail the results according to the network size (total number of species from guilds A and B) separated in three categories (<150, 150-200, or >200 species).

One-tailed Mantel tests based on the Pearson correlation (R) were performed using 10,000 permutations.

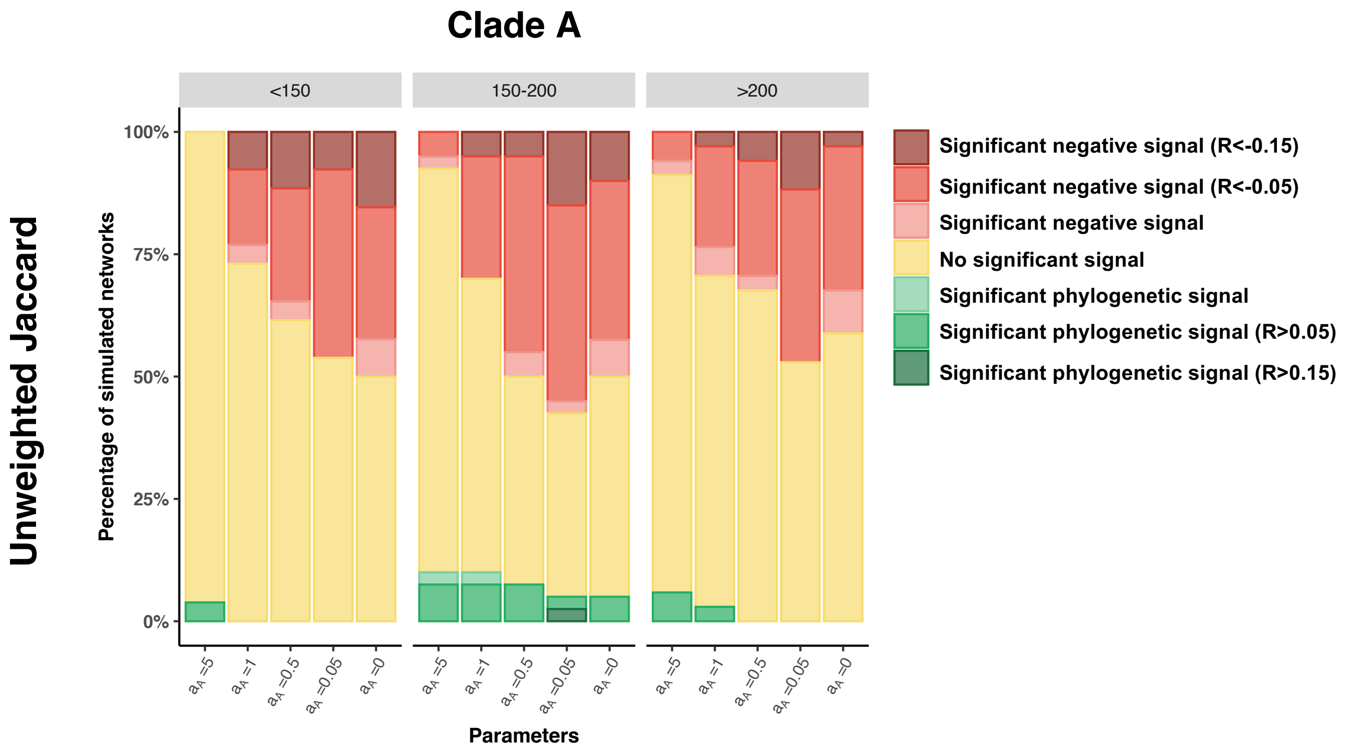

**Supplementary Figure 23: Phylogenetic signals in species interactions estimated using Mantel tests using permutations keeping constant the number of partners per species in guilds A in networks simulated with phylogenetic conservatism in the number, but not the identity, of partners:**

For each $a_A$ value, the bar indicates the percentage of simulated networks that present a significant positive correlation (in green; p-value<0.05 for the test of phylogenetic signal), a significant negative correlation (in red; p-value<0.05 for the test of negative phylogenetic signal), or no significant correlation (in yellow; both p-values>0.05).

$a_A$>0 corresponds to simulations where the number of partners of the species in guild A has evolved following an Ornstein-Uhlenbeck process.

$a_A$=0 corresponds to simulations where the number of partners of the species in guild A has evolved following a Brownian motion.

The different panels in rows correspond to the 2 tested ecological distances (Jaccard or UniFrac distances computed on unweighted networks) and for each panel, the subpanels detail the results according to the network size (total number of species from guilds A and B) separated in three categories (<150, 150-200, or >200 species).

One-tailed Mantel tests based on the Pearson correlation (R) were performed using 1,000 permutations keeping constant the number of partners per species in guilds A.

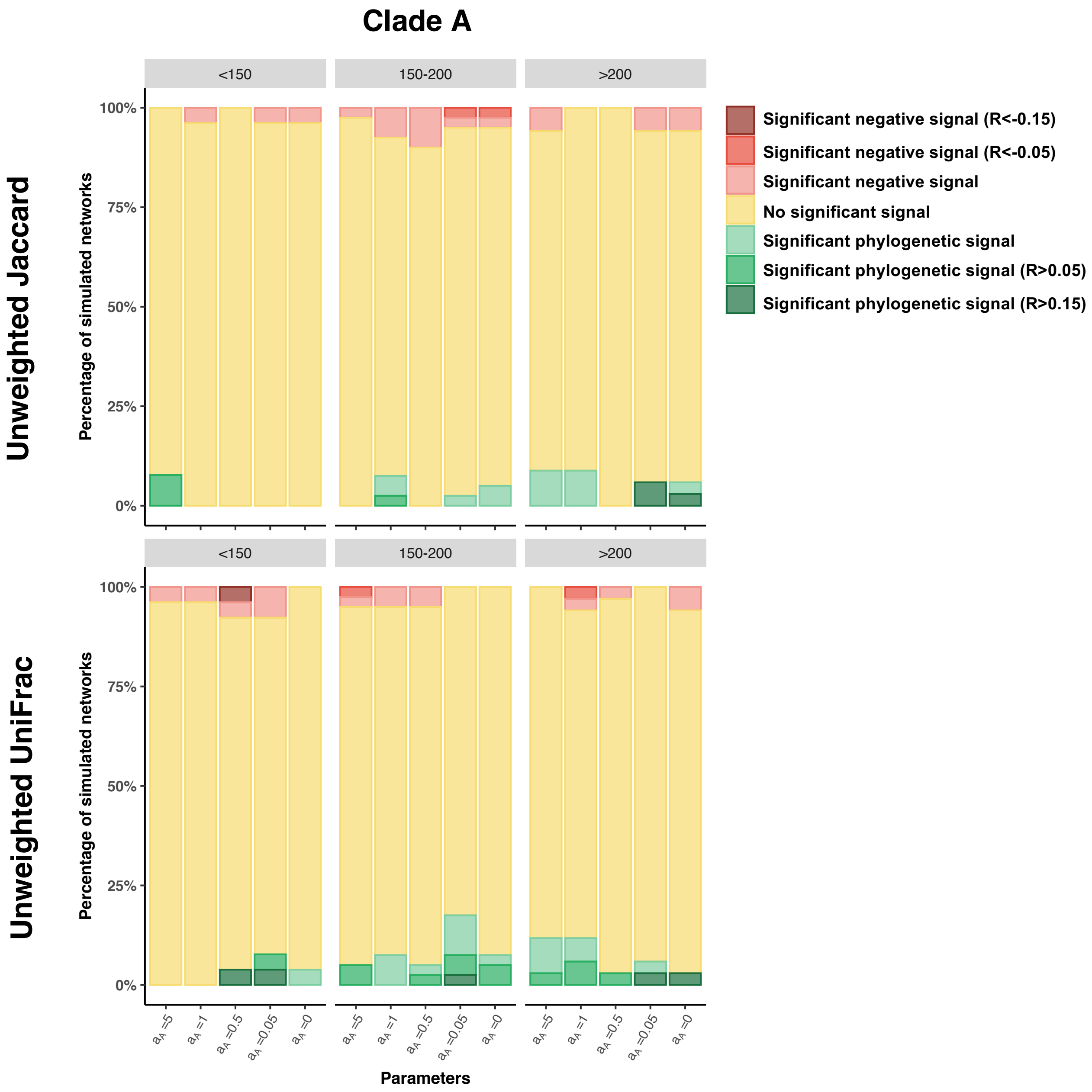

Clade A
Unweighted Jaccard
Unweighted UniFrac
Percentage of simulated networks
<150
150-200
>200
100%
75%
50%
25%
0%
Significant negative signal (R<-0.15)
Significant negative signal (R<-0.05)
Significant negative signal
No significant signal
Significant phylogenetic signal
Significant phylogenetic signal (R>0.05)
Significant phylogenetic signal (R>0.15)
$a_A$ =5
$a_A$ =1
$a_A$ =0.5
$a_A$ =0.05
$a_A$ =0
Parameters

### Testing phylogenetic uncertainty:

**Supplementary Figure 24: Phylogenetic uncertainty in the partners' trees increase when using short DNA sequences to reconstruct it:**

For each combination of parameters ($\alpha_A$ and $\alpha_B$), we indicate the correlation Pearson correlation between the original phylogenetic distances and the phylogenetic distances estimating after having reconstruing the tree from N-nucleotide-long DNA sequences simulated on the original tree. We tried N values from 1,200 (low phylogenetic uncertainty) to N=75 (high phylogenetic uncertainty).

The boxplots detail the results according to the network size (total number of species from guilds A and B) separated in three categories (<150, 150-250, or >250 species; see Table S5 for the number of simulated networks per category). Boxplots present the median surrounded by the first and third quartiles, and whiskers extend to the extreme values but no further than 1.5 of the inter-quartile range.

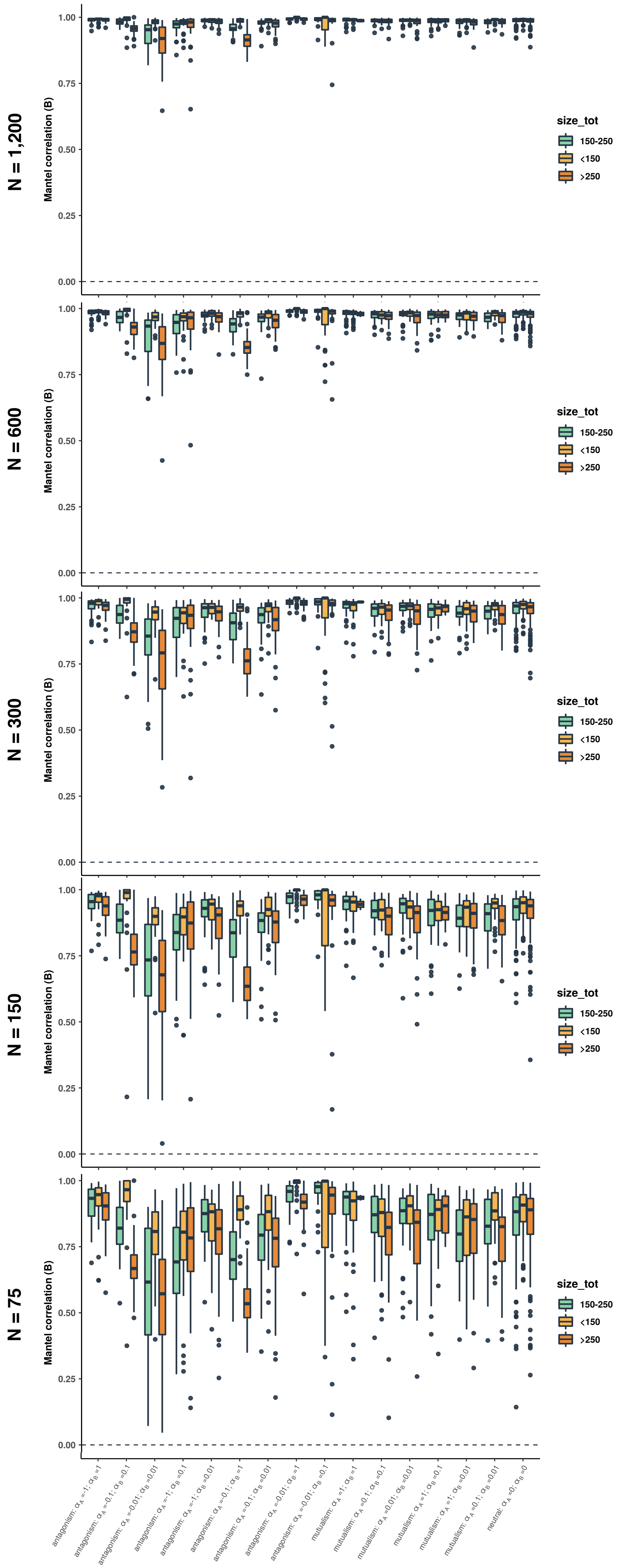
N = 1,200
N = 600
N = 300
N = 150
N = 75
Mantel correlation (B)
1.00
0.75
0.50
0.25
0.00
size_tot
150-250
<150
>250
antagonism: $\alpha_A$ =-1; $\alpha_B$ =1
antagonism: $\alpha_A$ =-0.1; $\alpha_B$ =0.1
antagonism: $\alpha_A$ =-0.01; $\alpha_B$ =0.01
antagonism: $\alpha_A$ =-1; $\alpha_B$ =0.1
antagonism: $\alpha_A$ =-1; $\alpha_B$ =0.01
antagonism: $\alpha_A$ =-0.1; $\alpha_B$ =1
antagonism: $\alpha_A$ =-0.1; $\alpha_B$ =0.01
antagonism: $\alpha_A$ =-0.01; $\alpha_B$ =1
antagonism: $\alpha_A$ =-0.01; $\alpha_B$ =0.1
mutualism: $\alpha_A$ =1; $\alpha_B$ =1
mutualism: $\alpha_A$ =0.1; $\alpha_B$ =0.1
mutualism: $\alpha_A$ =0.01; $\alpha_B$ =0.01
mutualism: $\alpha_A$ =1; $\alpha_B$ =0.1
mutualism: $\alpha_A$ =1; $\alpha_B$ =0.01
mutualism: $\alpha_A$ =0.1; $\alpha_B$ =0.01
neutral: $\alpha_A$ =0; $\alpha_B$ =0

**Supplementary Figure 25: The phylogenetic signals in species interactions estimated using simple Mantel tests in guilds A (left) and B (right) in *BipartiteEvol* simulations (Maliet *et al.*, 2020) vary according to the different levels of uncertainty in the partners' trees, but UniFrac distances still outperform Jaccard distances when there is some uncertainty in the phylogenetic tree of the partners:**

For each combination of parameters ($\alpha_A$ and $\alpha_B$), the bar indicates the percentage of simulated networks that present a significant positive correlation (in green; p-value<0.05 for the test of phylogenetic signal), a significant negative correlation (in red; p-value<0.05 for the test of negative phylogenetic signal), or no significant correlation (in yellow; both p-values>0.05).

The different panels in rows correspond to the different ecological distances from the original weighted (or generalized) UniFrac distances (on the top – computed on the original partners' phylogenetic trees), to the weighted UniFrac distances computed on the "noisy" partners' phylogenetic trees with a range level uncertainty (N= 1,200 … 75 base pairs), to the weighted Jaccard distances (on the bottom).

For each panel, the subpanels detail the results according to the network size (total number of species from guilds A and B) separated in three categories (<150, 150-250, or >250 species; see Table S5 for the number of simulated networks per category).

One-tailed Mantel tests based on the Pearson correlation (R) were performed using 10,000 permutations.

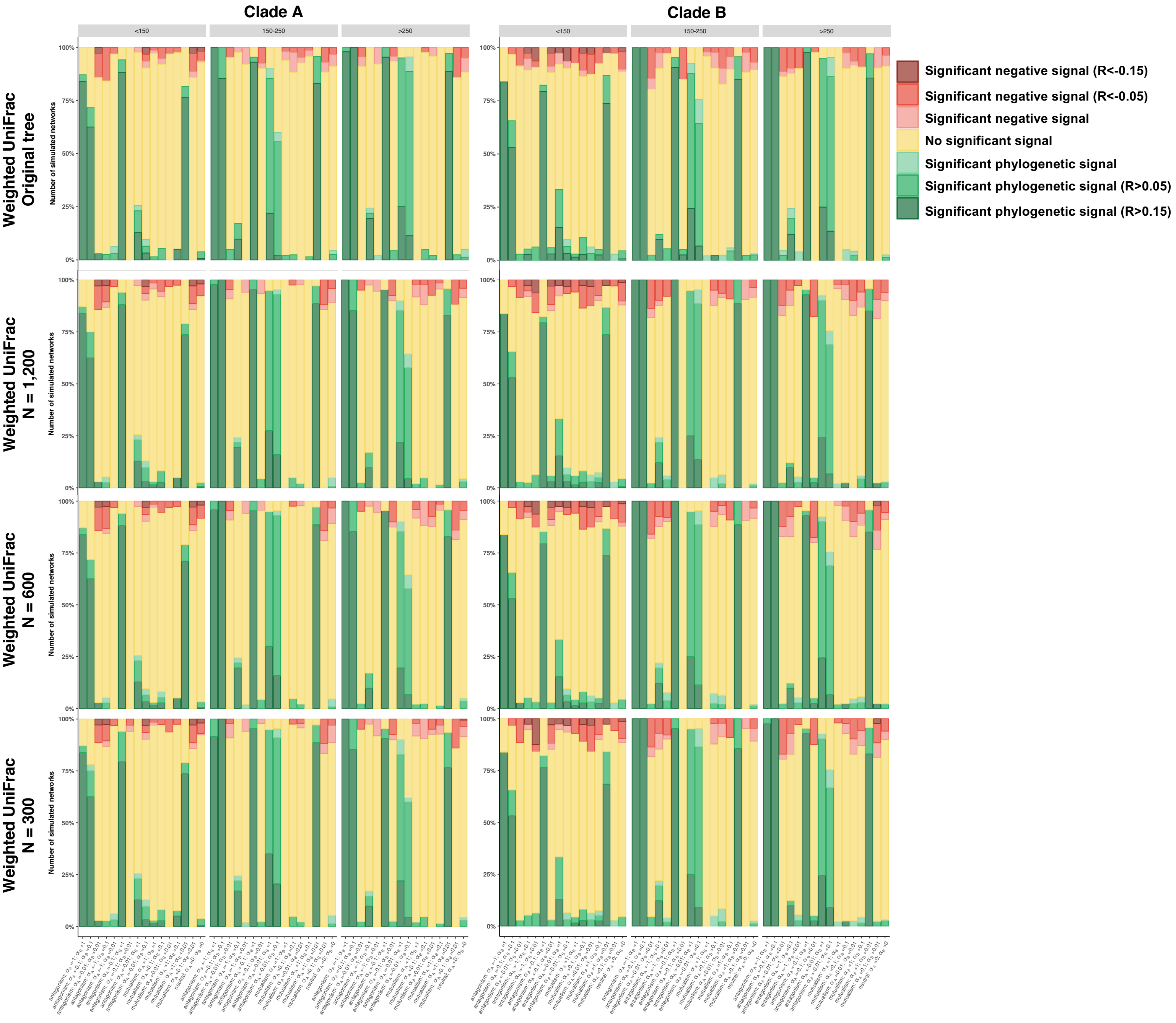
Clade A
Clade B
<150
150-250
>250
Weighted UniFrac Original tree
Weighted UniFrac N = 1,200
Weighted UniFrac N = 600
Weighted UniFrac N = 300
Number of simulated networks
100%
75%
50%
25%
0%
Significant negative signal (R<-0.15)
Significant negative signal (R<-0.05)
Significant negative signal
No significant signal
Significant phylogenetic signal
Significant phylogenetic signal (R>0.05)
Significant phylogenetic signal (R>0.15)

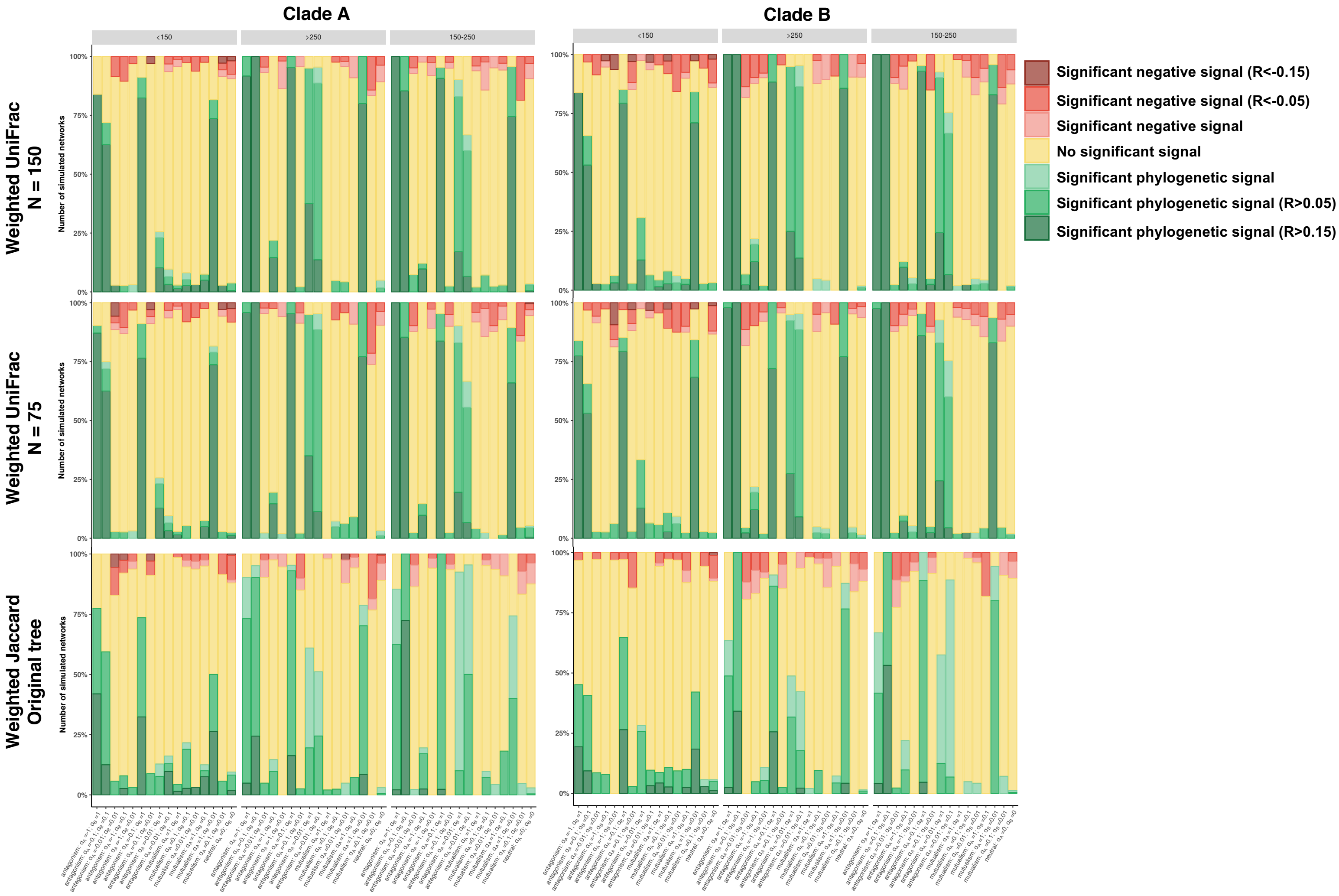
Clade A
Clade B
<150
>250
150-250
Weighted UniFrac N = 150
Weighted UniFrac N = 75
Weighted Jaccard Original tree
Number of simulated networks
100%
75%
50%
25%
0%
Significant negative signal (R<-0.15)
Significant negative signal (R<-0.05)
Significant negative signal
No significant signal
Significant phylogenetic signal
Significant phylogenetic signal (R>0.05)
Significant phylogenetic signal (R>0.15)

**Supplementary Figure 26: Example of the incorporation of uncertainty for one phylogenetic tree.**

The original phylogenetic distance matrix (a) - obtained from the *BipartiteEvol* simulations - is compared to the phylogenetic distance matrix of "noisy" reconstructed trees (b-f) obtained after simulating a nucleotidic alignment (with a number of nucleotide sites (N) between 1,200 base pairs (bp) and 75 bp) evolving on the original tree. Light yellow indicates that the phylogenetic distances are close to 0, whereas dark red represents high phylogenetic distances. While the external nodes tend to present important mismatch when N is low compared with the original matrix (the shorter the DNA sequences, the more uncertain are the phylogenetic reconstructions), the most internal nodes (and in particular the split between two major clades here) are still well visible in the phylogenetic distance matrices.

Yellow cells indicate low values whereas red cells correspond to high values.

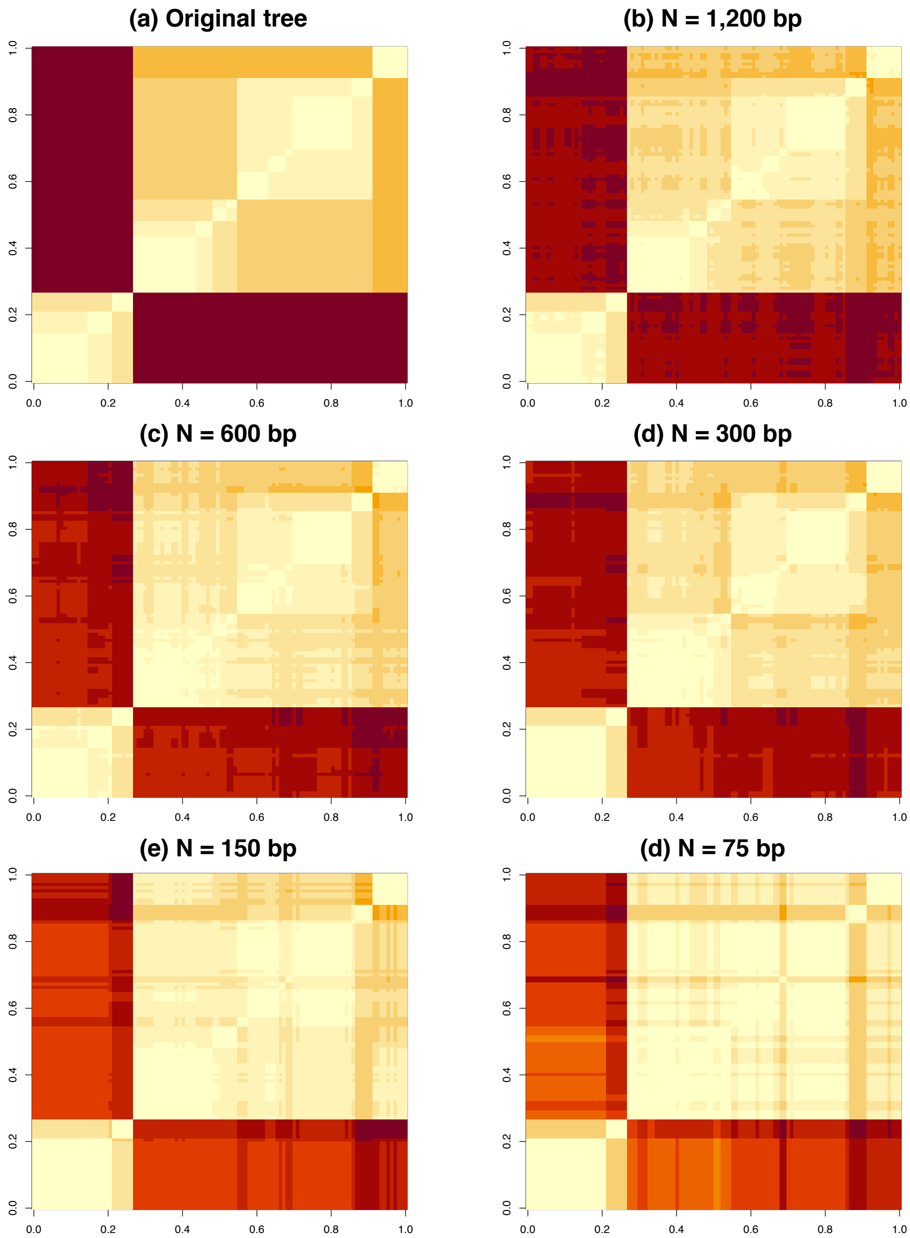

(a) Original tree
(b) N = 1,200 bp
(c) N = 600 bp
(d) N = 300 bp
(e) N = 150 bp
(d) N = 75 bp
0.0
0.2
0.4
0.6
0.8
1.0

### Sampling asymmetry in simulated interaction networks:

**Supplementary Figure 27: Size range of the networks simulated using *BipartiteEvol* (Maliet *et al.*, 2020) with sampling asymmetry.**

Numbers of species in guilds A (a) and B (b) as a function of the set of parameters ($\alpha_A$ and $\alpha_B$), and colored according to the total number of pairs of individuals for each guild (from 500 to 5,000).

Boxplots present the median surrounded by the first and third quartiles, and whiskers extend to the extreme values but no further than 1.5 of the inter-quartile range.

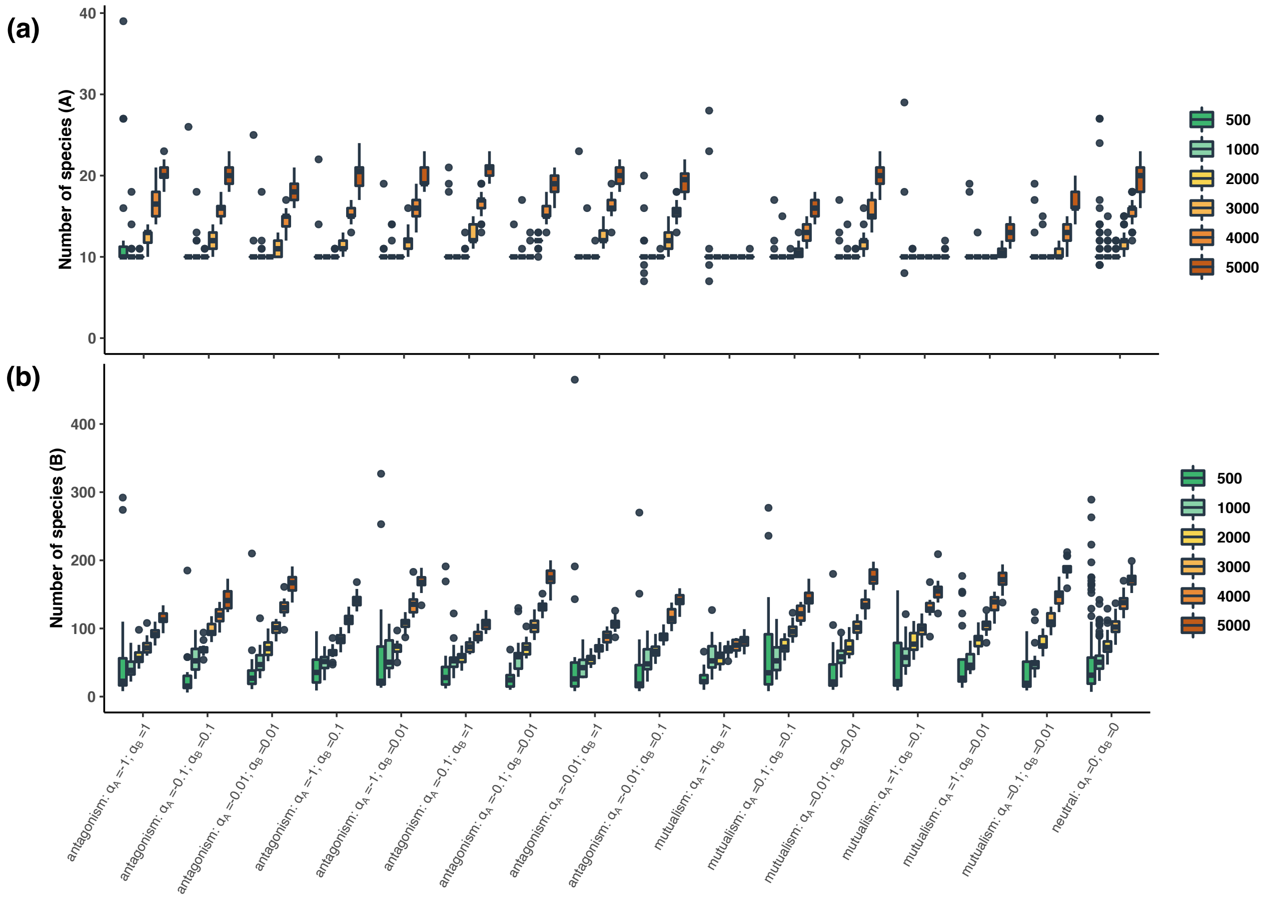

**Supplementary Figure 28: Realistic range of connectances (*i.e.* ratios of realized interactions) in networks simulated using *BipartiteEvol* (Maliet *et al.*, 2020) with sampling asymmetry.**

Connectances are represented as a function of the set of parameters ($\alpha_A$ and $\alpha_B$) and are colored according to the total number of pairs of individuals for each guild (from 500 to 5,000).

Boxplots present the median surrounded by the first and third quartiles, and whiskers extend to the extreme values but no further than 1.5 of the inter-quartile range.

Note that because we only sampled the 10% most abundant species, these asymmetrical networks had higher connectances than those from the original simulations.

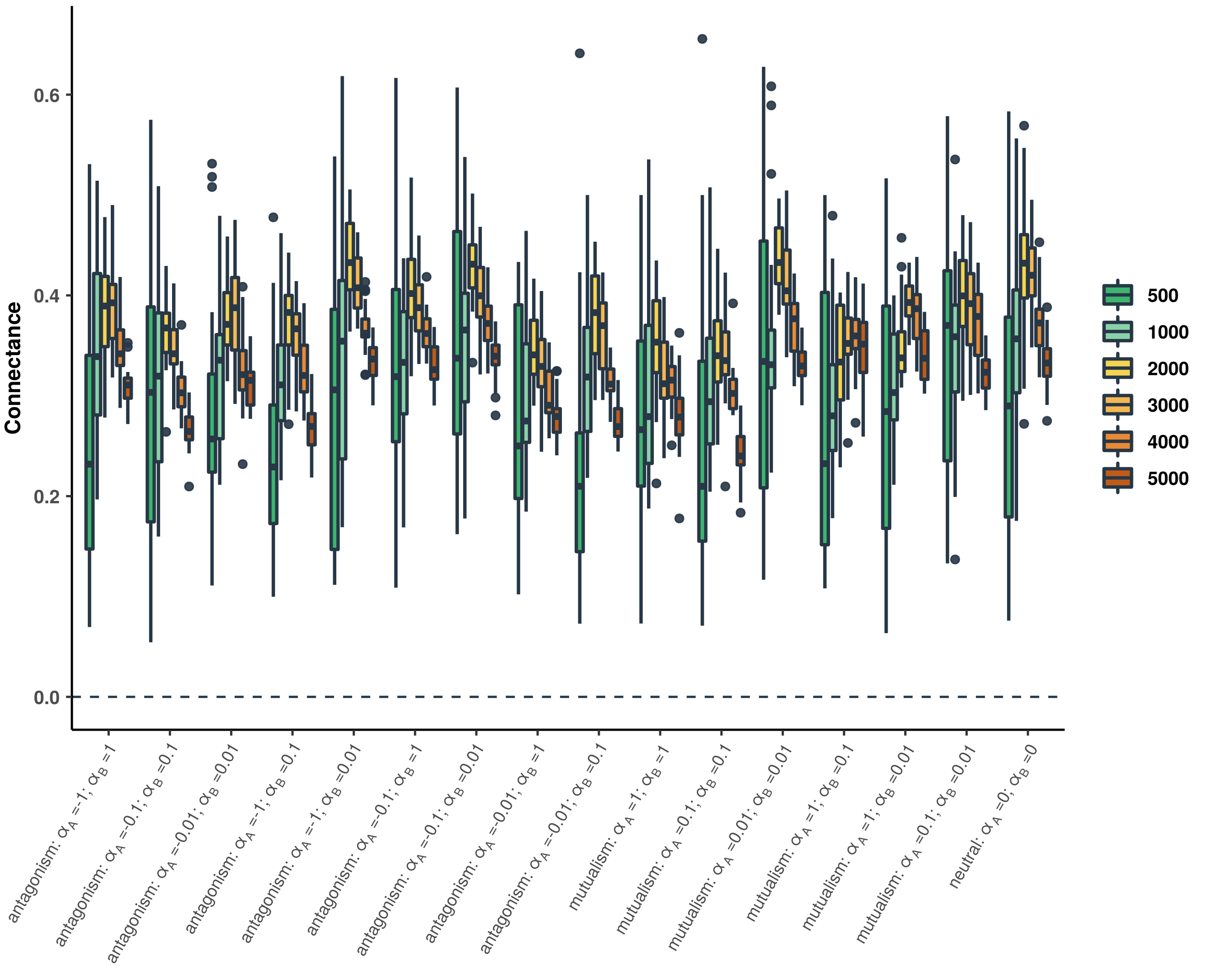

**Supplementary Figure 29: Phylogenetic signals in species interactions estimated using simple Mantel tests in guilds A (left) and B (right) simulated using *BipartiteEvol* (Maliet *et al.*, 2020) with sampling asymmetry.**

For each combination of parameters ($\alpha_A$ and $\alpha_B$), the bar indicates the percentage of simulated networks that present a significant positive correlation (in green; p-value<0.05 for the test of phylogenetic signal), a significant negative correlation (in red; p-value<0.05 for the test of negative phylogenetic signal), or no significant correlation (in yellow; both p-values>0.05).

The different panels in rows correspond to the 4 tested ecological distances (Jaccard or UniFrac distances computed with weighted or unweighted networks) and for each panel, the subpanels detail the results according to the network size (total number of species from guilds A and B) separated in three categories (<80, 80-140, or >140 species; see Table S6 for the number of simulated networks per category).

One-tailed Mantel tests based on the Pearson correlation (R) were performed using 10,000 permutations.

Importantly, these results indicated that Mantel tests seem robust to alternative sampling strategies such as the asymmetrical sampling of the guilds, which is particularly frequent when studying microbial symbiosis (Jacquemyn *et al.*, 2011; Martos *et al.*, 2012; Song *et al.*, 2020).

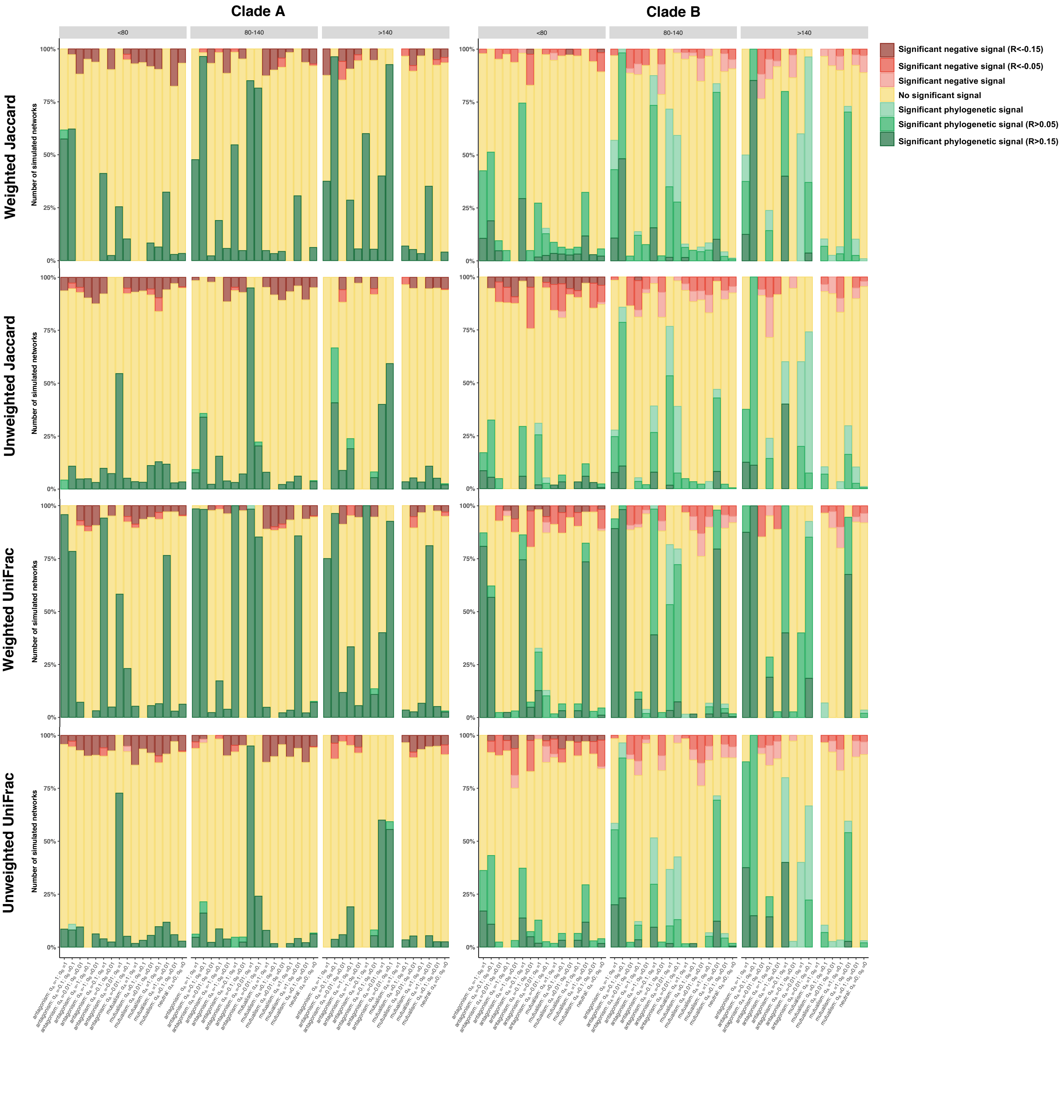

Clade A
Clade B
<80
80-140
>140
Weighted Jaccard
Unweighted Jaccard
Weighted UniFrac
Unweighted UniFrac
Number of simulated networks
100%
75%
50%
25%
0%
Significant negative signal (R<-0.15)
Significant negative signal (R<-0.05)
Significant negative signal
No significant signal
Significant phylogenetic signal
Significant phylogenetic signal (R>0.05)
Significant phylogenetic signal (R>0.15)

**Supplementary Figure 30: Phylogenetic signal estimated using the Phylogenetic bipartite linear model (PBLM; Ives & Godfray, 2006) in the networks simulated using *BipartiteEvol* (Maliet *et al.*, 2020) with sampling asymmetry.**

For each combination of parameters ($\alpha_A$ and $\alpha_B$), the bar indicates the percentage of simulated networks that present no significant (in yellow; $MSE \geq MSE_{star}$) or a significant (in green; $MSE < MSE_{star}$) phylogenetic signals. Phylogenetic signals are shaded from light green to dark green according to the strength of the signal (e.g. in dark green if $d_A > 0.15$ or $d_B > 0.15$). PBLM was run on the weighted networks (a) or on the unweighted networks (b).

For each panel, the subpanels detail the results according to the network size (total number of species from guilds A and B) separated in three categories (<80, 80-140, or >140 species). See Table S6 for the number of simulated networks per category.

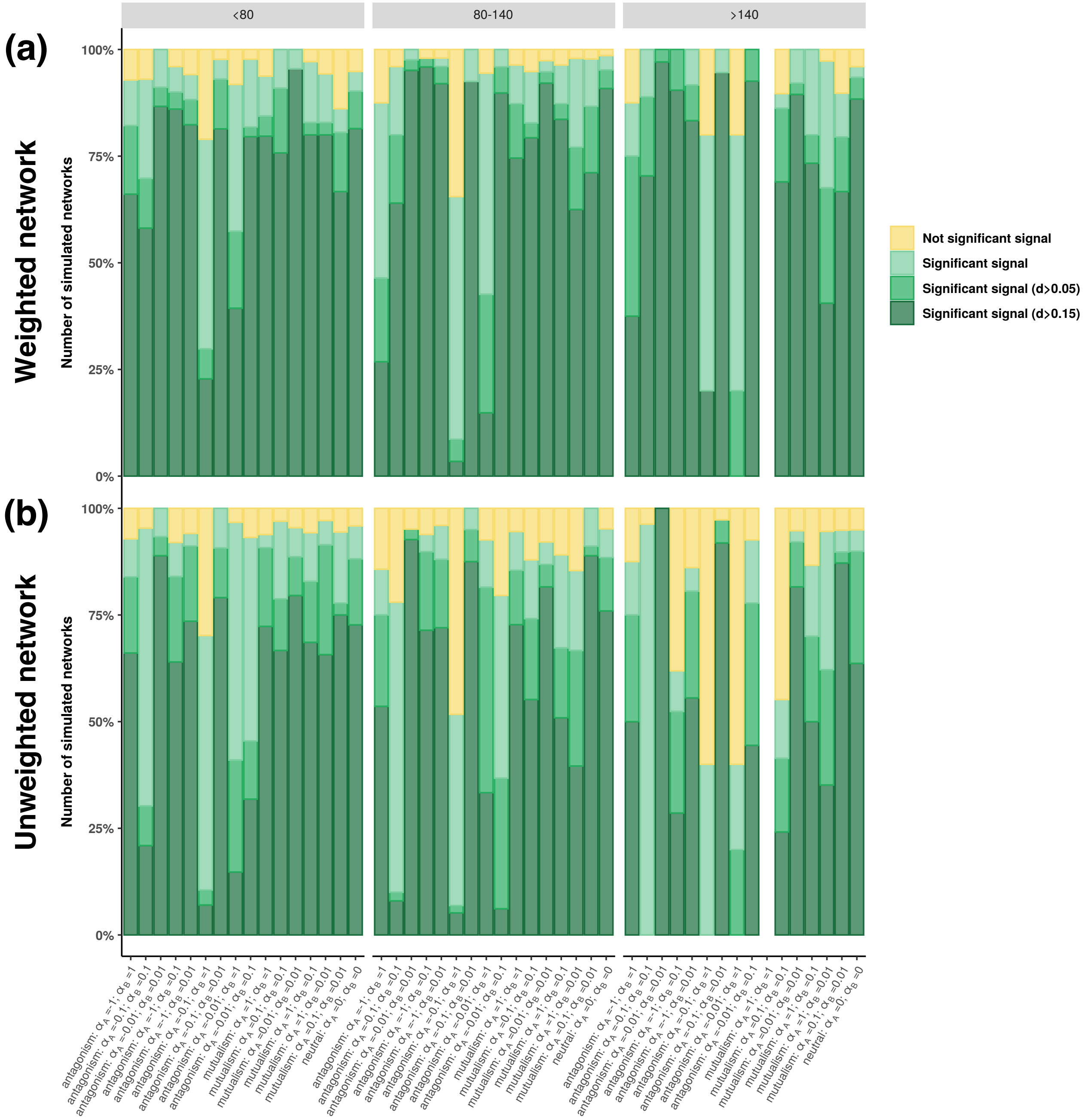

<80
80-140
>140
(a)
Weighted network
(b)
Unweighted network
Number of simulated networks
100%
75%
50%
25%
0%
Not significant signal
Significant signal
Significant signal (d>0.05)
Significant signal (d>0.15)

### Detecting clade-specific phylogenetic signal:

**Supplementary Figure 31: Example of clade-specific phylogenetic signals measured on the phylogeny of guild A (a) and B (b) in a *BipartiteEvol* simulation.**

The example was obtained by merging a simulation obtained with the parameters "mutualism (v)" into a neutral simulation. The asterisks represent where the "mutualistic clades" were merged in the "neutral backbone tree".

Each node with at least 10 descendent species is colored according to the result of the Mantel test (its significance and the strength of the Pearson correlation (R) when significant (in red)) performed on the corresponding sub-network. One-tailed simple Mantel tests were performed using the Pearson correlation and 100,000 permutations and their significance were evaluated while correcting for multiple testing (Bonferroni correction).

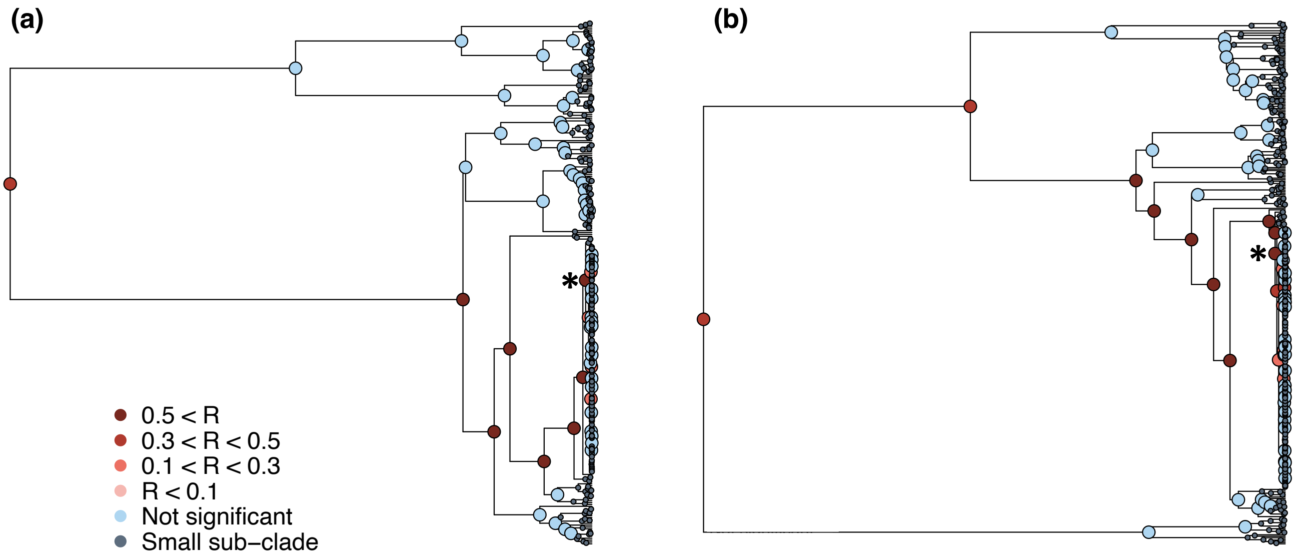

### Empirical application on an orchid-fungus interaction network:

**Supplementary Figure 32: The significant phylogenetic signal in the epiphytic subtribe Angraecineae seems robust to polytomies and subsampling in Angraecineae.**

Clade-specific analyses of phylogenetic signal performed on the orchid phylogeny where a polytomy has been added for the well-resolved *Angraecum* genus (a) and only 10 Angraecineae species were sampled (b).

Each node with at least 10 descendent species is colored according to the results of the Mantel test (its significance and the strength of the Pearson correlation (R)) performed on the corresponding sub-network. One-tailed simple Mantel tests were performed using the Pearson correlation and 100,000 permutations and their significance was evaluated while correcting for multiple testing (Bonferroni correction).

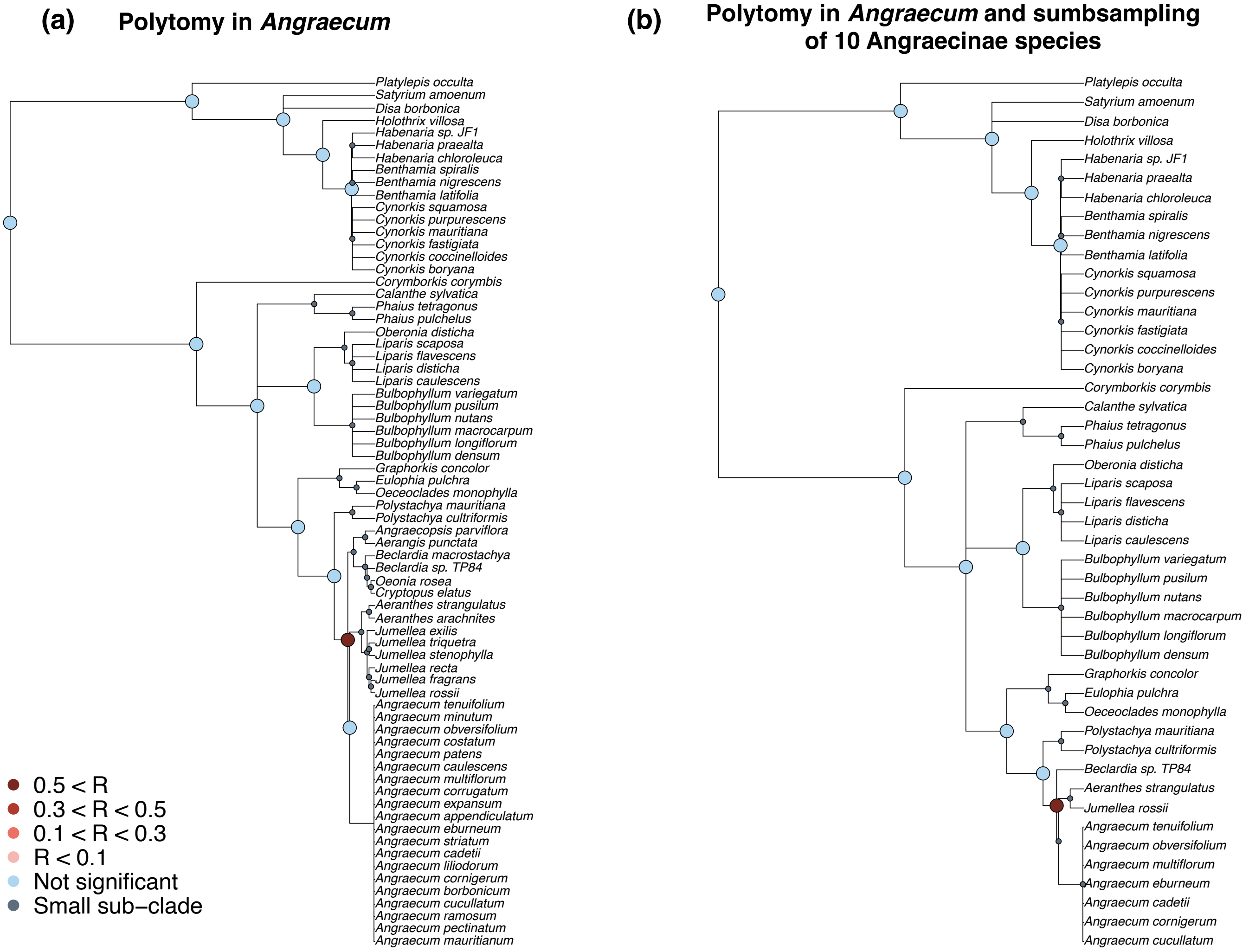
(a)
Polytomy in *Angraecum*
*Platylepis occulta*
*Satyrium amoenum*
*Disa borbonica*
*Holothrix villosa*
*Habenaria sp. JF1*
*Habenaria praealta*
*Habenaria chloroleuca*
*Benthamia spiralis*
*Benthamia nigrescens*
*Benthamia latifolia*
*Cynorkis squamosa*
*Cynorkis purpurescens*
*Cynorkis mauritiana*
*Cynorkis fastigiata*
*Cynorkis coccinelloides*
*Cynorkis boryana*
*Corymborkis corymbis*
*Calanthe sylvatica*
*Phaius tetragonus*
*Phaius pulchelus*
*Oberonia disticha*
*Liparis scaposa*
*Liparis flavescens*
*Liparis disticha*
*Liparis caulescens*
*Bulbophyllum variegatum*
*Bulbophyllum pusilum*
*Bulbophyllum nutans*
*Bulbophyllum macrocarpum*
*Bulbophyllum longiflorum*
*Bulbophyllum densum*
*Graphorkis concolor*
*Eulophia pulchra*
*Oeceoclades monophylla*
*Polystachya mauritiana*
*Polystachya cultriformis*
*Angraecopsis parviflora*
*Aerangis punctata*
*Beclardia macrostachya*
*Beclardia sp. TP84*
*Oeonia rosea*
*Cryptopus elatus*
*Aeranthes strangulatus*
*Aeranthes arachnites*
*Jumellea exilis*
*Jumellea triquetra*
*Jumellea stenophylla*
*Jumellea recta*
*Jumellea fragrans*
*Jumellea rossii*
*Angraecum tenuifolium*
*Angraecum minutum*
*Angraecum obversifolium*
*Angraecum costatum*
*Angraecum patens*
*Angraecum caulescens*
*Angraecum multiflorum*
*Angraecum corrugatum*
*Angraecum expansum*
*Angraecum appendiculatum*
*Angraecum eburneum*
*Angraecum striatum*
*Angraecum cadetii*
*Angraecum liliodorum*
*Angraecum cornigerum*
*Angraecum borbonicum*
*Angraecum cucullatum*
*Angraecum ramosum*
*Angraecum pectinatum*
*Angraecum mauritianum*
0.5 < R
0.3 < R < 0.5
0.1 < R < 0.3
R < 0.1
Not significant
Small sub−clade
(b)
Polytomy in *Angraecum* and sumbsampling of 10 Angraecinae species
*Platylepis occulta*
*Satyrium amoenum*
*Disa borbonica*
*Holothrix villosa*
*Habenaria sp. JF1*
*Habenaria praealta*
*Habenaria chloroleuca*
*Benthamia spiralis*
*Benthamia nigrescens*
*Benthamia latifolia*
*Cynorkis squamosa*
*Cynorkis purpurescens*
*Cynorkis mauritiana*
*Cynorkis fastigiata*
*Cynorkis coccinelloides*
*Cynorkis boryana*
*Corymborkis corymbis*
*Calanthe sylvatica*
*Phaius tetragonus*
*Phaius pulchelus*
*Oberonia disticha*
*Liparis scaposa*
*Liparis flavescens*
*Liparis disticha*
*Liparis caulescens*
*Bulbophyllum variegatum*
*Bulbophyllum pusilum*
*Bulbophyllum nutans*
*Bulbophyllum macrocarpum*
*Bulbophyllum longiflorum*
*Bulbophyllum densum*
*Graphorkis concolor*
*Eulophia pulchra*
*Oeceoclades monophylla*
*Polystachya mauritiana*
*Polystachya cultriformis*
*Beclardia sp. TP84*
*Aeranthes strangulatus*
*Jumellea rossii*
*Angraecum tenuifolium*
*Angraecum obversifolium*
*Angraecum multiflorum*
*Angraecum eburneum*
*Angraecum cadetii*
*Angraecum cornigerum*
*Angraecum cucullatum*

**Supplementary Figure 33: The significant phylogenetic signal in the epiphytic subtribe Angraecineae is robust to the arbitrary age we set for the polytomies of unresolved orchid genera.**

Clade-specific analyses of phylogenetic signal performed on the orchid phylogenies reconstructed with different ages for the polytomies corresponding to unresolved genera. We set 6 ages from 12 Myr ago to 2 Myr ago. For a given genus, if the arbitrary age of the polytomy was older than the divergence time with another sampled genus, we used this divergence time as the age of the polytomy instead.

For each orchid phylogeny, we reported its clade-specific phylogenetic signal. Each node with at least 10 descendent species is colored according to the results of the Mantel test (its significance and the strength of the Pearson correlation (R)) performed on the corresponding sub-network. One-tailed simple Mantel tests were performed using the Pearson correlation and 100,000 permutations; their significance was evaluated while correcting for multiple testing (Bonferroni correction).

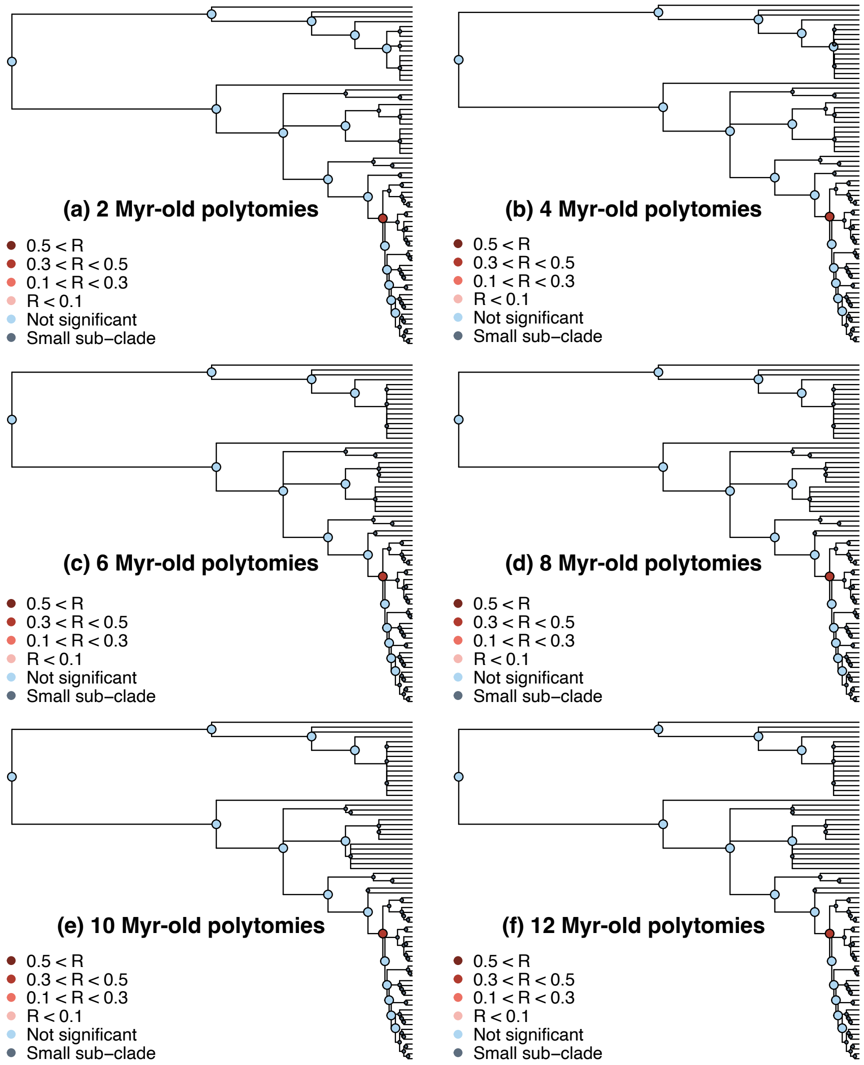
